# A selective inhibitor of oncogenic JNK signalling perturbs metastatic outgrowth of triple-negative breast cancer through metabolic blockade

**DOI:** 10.1101/2025.04.16.649243

**Authors:** Sharissa L Latham, Yolande EI O’Donnell, Oscar Girgis-Cook, Misaki S Clearwater, Nicole S Bryce, Ellie TY Mok, Sophie A Lynn, Jenny Ni, Stephanie Alfred, King Ho Leong, Lake-ee Quek, Kendelle J Murphy, Chiara Pantarelli, Iliya Dragutinovic, Marjan M Naeini, Claudia Thiel, Antonia Cadell, Monica Phimmachanh, Rashmi Sharma, Ewan Millar, Jeremy ZR Han, Jordan F Hastings, Martin Köhler, Tilman Brummer, Adelaide Young, Sandra O’Toole, Samantha R Oakes, Edna Hardeman, John Lock, Walter Kolch, Manuel H Taft, Max Nobis, Leonard Goldstein, Paul Timpson, Benjamin L Parker, Jeff Holst, Peter W Gunning, Thomas R Cox, Jonathan C Morris, David R Croucher

## Abstract

Although c-Jun N-terminal Kinase (JNK) represents an attractive anti-cancer target, its pleiotropic functionality limits the use of direct JNK inhibitors. Here, we identify a distinct subcellular pattern of JNK activity as a therapeutic vulnerability in breast cancer, where cytoplasmic JNK activity predicts poor survival outcomes, is elevated in triple-negative breast cancers (TNBC) and is essential for metastatic outgrowth. Mechanistic analyses reveal cytoplasmic JNK acts through multiple mechanisms, with downstream targets involved in cellular metabolism and cytoskeletal regulation. On this basis, we leveraged actin-based phenotypic drug-screening and identified K12, an indirect but selective inhibitor of cytoplasmic JNK that blocks TNBC metastatic outgrowth *in vivo*. We reveal that K12 inhibits glutaminase-1 and the pyruvate dehydrogenase complex, and that this poly-pharmacology overcomes pyruvate anaplerosis, a known resistance mechanism of existing glutaminase inhibitors. These findings demonstrate the potential of selectively targeting the oncogenic function of JNK, offering new treatment options for early-stage metastatic TNBC.

## Introduction

Triple negative breast cancer (TNBC) is a highly metastatic subtype of breast cancer, accounting for ∼15% of all breast cancer cases and disproportionately affecting young, premenopausal women of specific racial backgrounds^1,2^. Despite significant progress in treating TNBC over recent years, as seen with the incorporation of PARP inhibitors, Trop2-targeting antibody-drug conjugates, and immunotherapies into the TNBC treatment landscape^3^, there remain limited options for effectively targeting metastatic disease. TNBC tumours are distinguished by their increased propensity to develop visceral metastases^4,5^, and patients with this disease have a significantly increased risk of distant recurrence within two years of diagnosis^6^. As metastatic disease ultimately underpins high mortality rates in these patients^7^, there is a critical need for the identification and development of therapies that specifically target metastatic disease progression to improve TNBC patient outcomes.

The c-Jun N-terminal Kinases (JNK) are members of the mitogen activated protein kinase (MAPK) family of serine/threonine protein kinases and have emerged as key drivers of metastatic TNBC. This stems from several studies demonstrating the effectiveness of ATP-competitive JNK inhibitors in halting pulmonary metastases in immunocompromised and syngeneic mouse models of TNBC^8–11^. Whilst JNK is an attractive therapeutic target in this context, genetic knockout studies reveal that it also plays critical tumour-suppressive roles in normal breast tissue. Specifically, JNK has been shown to maintain normal mammary gland architecture through the regulation of gland branching and luminal clearance^12,13^, and to prevent the initiation of early tumorigenic events by maintaining genomic stability and DNA repair^14,15^. Furthermore, as JNK activity is also required for chemotherapy-induced apoptosis^16^, it is unlikely that direct JNK inhibitors could be incorporated within many existing standard-of-care treatment regimens. As such, for clinical success alternate JNK-targeting strategies will be required to specifically target the oncogenic functions of the JNK signalling pathway without perturbing its tumour-suppressive or apoptotic activities^17^.

In this study, we aimed to delineate the oncogenic functions of the JNK signalling pathway that drive metastatic progression within established TNBC models and utilise this understanding to identify selective inhibitors of oncogenic JNK signalling. Our results uncover spatially and prognostically distinct pools of JNK activity in breast tissue and demonstrate that JNK activity within the cytoplasm of TNBC cells is essential for metastatic outgrowth. Furthermore, our phenotypic drug screening approach led to the identification of K12, a small molecule that selectively inhibits the oncogenic functions of JNK and perturbs the growth of pulmonary metastases through its dual-targeting of glutaminase-1 (GLS) and the pyruvate dehydrogenase complex (PDHC). This study provides proof-of-principle data that discrete functions of this pleiotropic signalling pathway can be selectively targeted, opening up new strategies to treat early metastatic outgrowth and prevent relapse in TNBC patients.

## Results

### Spatially and prognostically distinct subcellular JNK activity in breast tissue

An immunohistochemical (IHC) assessment of JNK activity in breast cancer^18^ and TNBC^19^ patient cohorts revealed contrasting prognostic roles for nuclear and cytoplasmic JNK activity in breast tissue. Whilst we observed prominent phospho-JNK (Thr183/Tyr185, ppJNK) staining within the nuclei of luminal epithelial cells in normal breast tissue, both a loss of nuclear ppJNK and a gain of cytoplasmic ppJNK were observed to varying degrees in tumour tissue (**Fig. 1A**). Subtype-specific changes in JNK activity were also apparent in these cohorts (**Fig. 1B**), with TNBC tumours displaying both a significant loss of nuclear ppJNK (**Fig. 1C**) and gain of cytoplasmic ppJNK (**Fig 1D**). Survival analysis revealed that any deviation from the “normal” high nuclear/low cytoplasmic JNK activity profile was associated with poorer overall (**Fig. 1E**) and relapse-free survival (**Fig. 1F**). These associations between subcellular JNK activity and survival outcomes were upheld when TNBC patients were excluded from the analysis (**Fig. S1A-C**), demonstrating that the poor survival outcomes associated with this subtype were not biasing these results. Together, these data suggest distinct and opposing roles for JNK within the nucleus and cytoplasm of breast tissue, with nuclear JNK activity appearing to be tumour-suppressive and cytoplasmic JNK activity oncogenic, particularly in TNBC.

**Figure 1.**
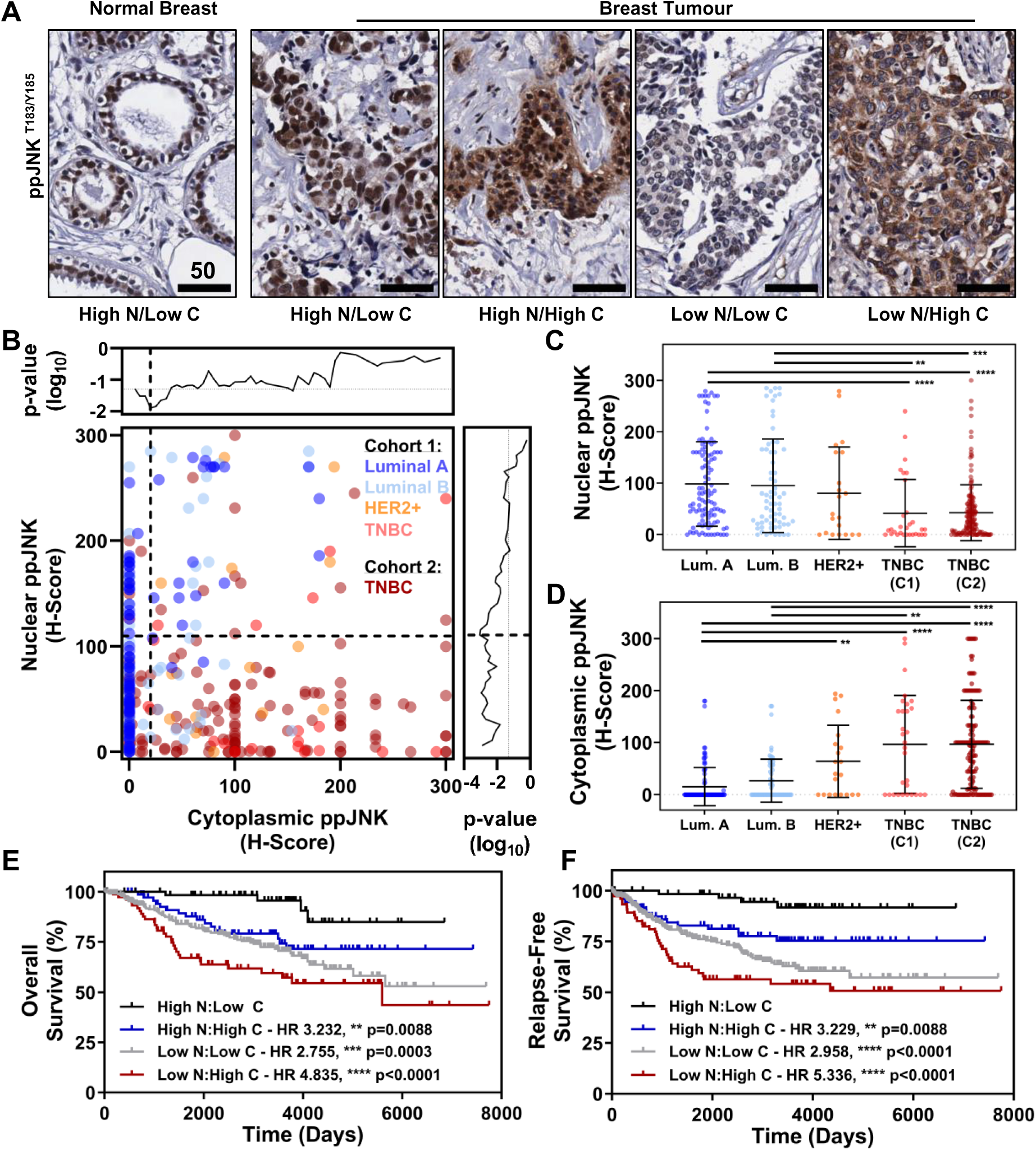
Prognostic significance of subcellular JNK activity in breast cancer. **(A)** Representative images of variable nuclear (N) and cytoplasmic (C) staining from IHC analyses of ppJNK (Thr183/Tyr185) in normal breast tissue and a pan-breast cancer cohort. Scale bars are in µm. **(B)** Nuclear vs cytoplasmic ppJNK H-Scores from both the pan-breast cancer (cohort 1: n=457) and TNBC validation (cohort 2: n=161) cohorts, classified according to subtype. H-Score thresholds defining low/high ppJNK staining were determined through p-value scanning (Gehan-Breslow-Wilcoxon test). Comparative analysis of **(C)** nuclear and **(D)** cytoplasmic ppJNK staining between breast cancer subtypes in the pan-breast cancer (C1) and TNBC validation (C2) cohorts (mean ± SD; Kruskal-Wallis with Dunns post-test). Kaplan-Meier analysis of **(E)** overall survival and **(F)** relapse-free survival in the pan-breast cancer cohort (log-rank test, Mantel-Haenszel hazard ratio). For both analyses, low nuclear JNK ≤ 110, high nuclear ppJNK >110, low cytoplasmic ppJNK ≤ 20, high cytoplasmic ppJNK >20. In all cases: * p<0.05, ** p<0.01, *** p<0.001 and *** p<0.0001.

### Direct JNK inhibitors block both oncogenic and tumour-suppressive JNK functions

Given the previous use of ATP-competitive, direct JNK inhibitors in models of TNBC^8–11^, we next adopted a syngeneic orthotopic mouse model to monitor the impact of the simultaneous JNK inhibition that would occur within both of these subcellular compartments in this context. This model utilised 4T1 TNBC cells, which have elevated levels of cytoplasmic ppJNK compared to normal breast tissue, which has strong nuclear JNK activity (**Fig. 2A**). Using CC-401, a direct JNK inhibitor previously used in MDA-MB-231 *in vivo* models of metastatic TNBC^8^, we show that although JNK inhibition had no effect on primary tumour growth (**Fig. 2B**) or end-point tumour burden (**Fig. 2C**), it significantly reduced the number of lung metastases (**Fig. 2D-E**). Whilst these data support the therapeutic potential of JNK inhibition in TNBC, examination of the contralateral native breast tissue revealed that CC-401 simultaneously perturbed normal mammary gland architecture (**Fig. 2F**), as seen by a significant increase in duct branch number (**Fig. 2G**) and width (**Fig. 2H**). Using DNA damage as an orthogonal read-out of tumour-suppressive JNK activity^15^ (**Fig. 2I**), we further demonstrate that CC-401 treatment also increased phospho-histone H2A.X (Ser139, γH2AX) staining within luminal epithelial cells, as measured by both positive pixel analysis (**Fig. 2J**) and average intensity (**Fig. 2K**). These observations closely parallel previously published *in vivo* JNK knockout models and support the role of nuclear JNK activity in preventing tumour initiation in normal breast tissue^12,15^. Further, these data demonstrate that direct JNK inhibitors are unlikely to yield clinical success in breast cancer, as they are unable to discriminate between these discrete subcellular JNK pools and instead block both the oncogenic and tumour-suppressive functions of JNK.

**Figure 2.**
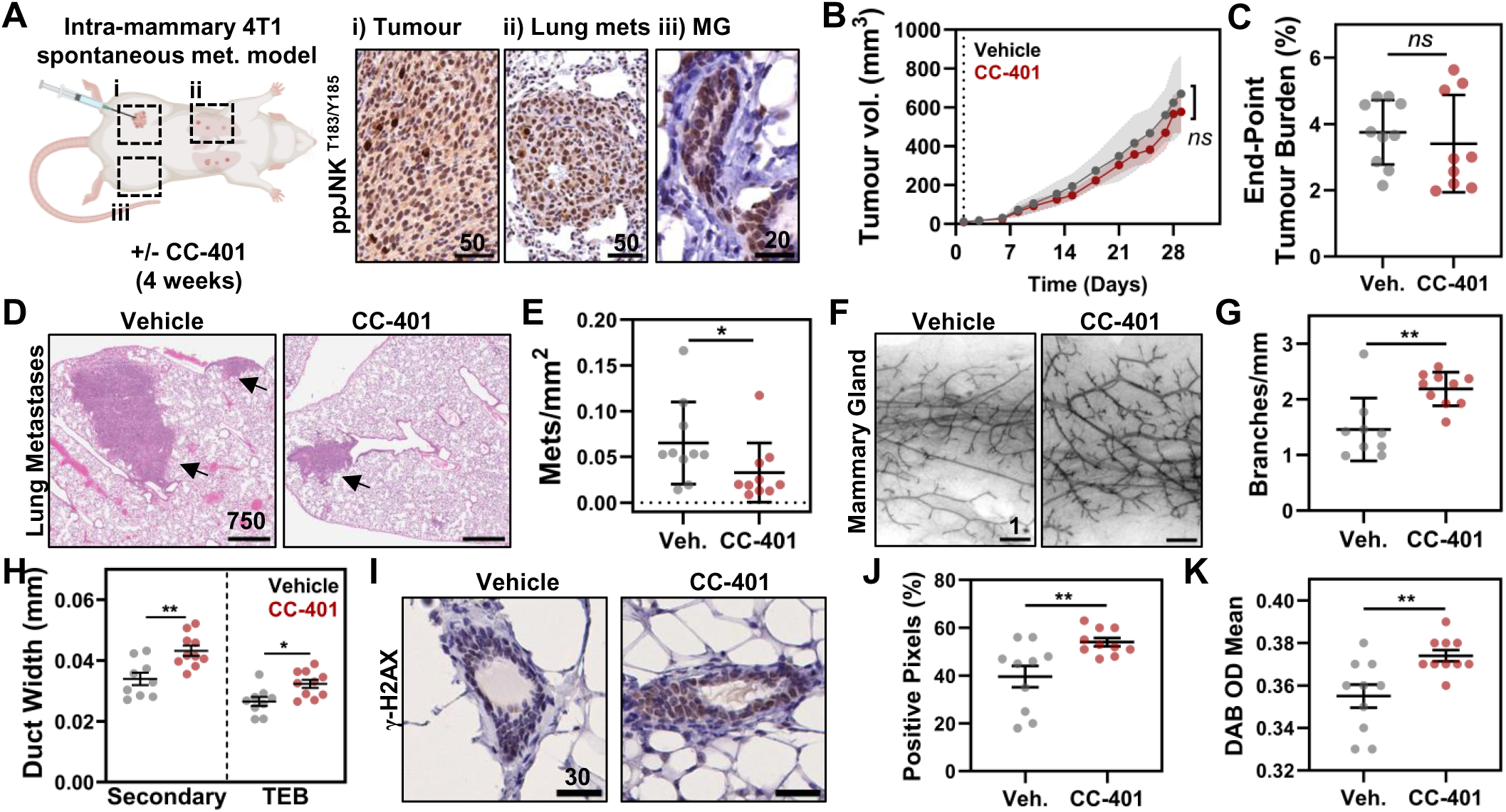
A direct JNK inhibitor blocks both oncogenic and tumour-suppressive JNK functions. **(A)** Schematic of the intra-mammary 4T1 mouse model, including representative images of ppJNK staining within i) 4T1 primary tumour, ii) lung metastases (mets) and iii) Balb/C native mammary glands (MG). **(B)** Primary tumour growth in vehicle and CC-401 (25 mg/kg) treated mice (mean ± SD, n=10; mixed effects analysis, Sidaks post-test). **(C)** Tumour burden at experimental endpoint (mean ± SD, n=10 vehicle/n=9 CC-401; t-test). **(D)** Representative images of H&E-stained lung metastases at experimental endpoint. **(E)** Quantitative assessment of lung metastases (mean ± SD, n=10). **(F)** Representative images of carmine-stained mammary glands. **(G)** Quantitative assessment of mammary gland branching (mean ± SD, n=9 vehicle/n=10 CC-401; Mann-Whitney test). **(H)** Analysis of secondary ductal branches and terminal end buds (TEB) in mammary glands (mean ± SEM, n=9 vehicle/n=10 CC-401; t-tests). **(I)** Representative images of γ-H2AX staining in mammary gland cross-sections. Quantitative assessment of γ-H2AX staining within the mammary gland luminal epithelial layer by **(J)** positive pixel analysis and **(K)** staining intensity (mean ± SEM, n=10; Mann-Whitney test). All scale bars are in µm.

### Cytoplasmic JNK inhibition blocks TNBC metastatic outgrowth

To validate this oncogenic role for cytoplasmic JNK in TNBC, we utilised genetically-encoded, doxycycline-inducible localisation-specific JNK inhibitors^20^ within the MDA-MB-231 cell line (**Fig. 3A**), which displays both cytoplasmic and nuclear JNK activity *in vivo* (**Fig. 3B**). These inhibitors incorporate an inhibitory peptide from the JNK scaffold protein JIP-1, fused to EGFP and either a nuclear export (C-JNKi) or localisation (N-JNKi) sequence. Using an orthotopic intraductal model, the effect of nuclear and cytoplasmic JNK inhibition on primary tumour growth and end-point metastatic burden was assessed in mice receiving normal vs doxycycline chow (**Fig. 3C**). Consistent with the impact of CC-401, no apparent effect of either the cytoplasmic or nuclear JNK inhibitor was observed on primary tumour growth (**Fig. 3D**), whilst the C-JNKi significantly inhibited metastatic burden within the lungs (**Fig. 3E-F**). Close examination of mice injected with C-JNKi expressing MDA-MB-231 cells, revealed the presence of pulmonary micro-metastases (**Fig. 3G**), suggesting that cytoplasmic JNK inhibition does not affect metastatic dissemination, but rather the ability of these cells to proliferate within the metastatic niche.

**Figure 3.**
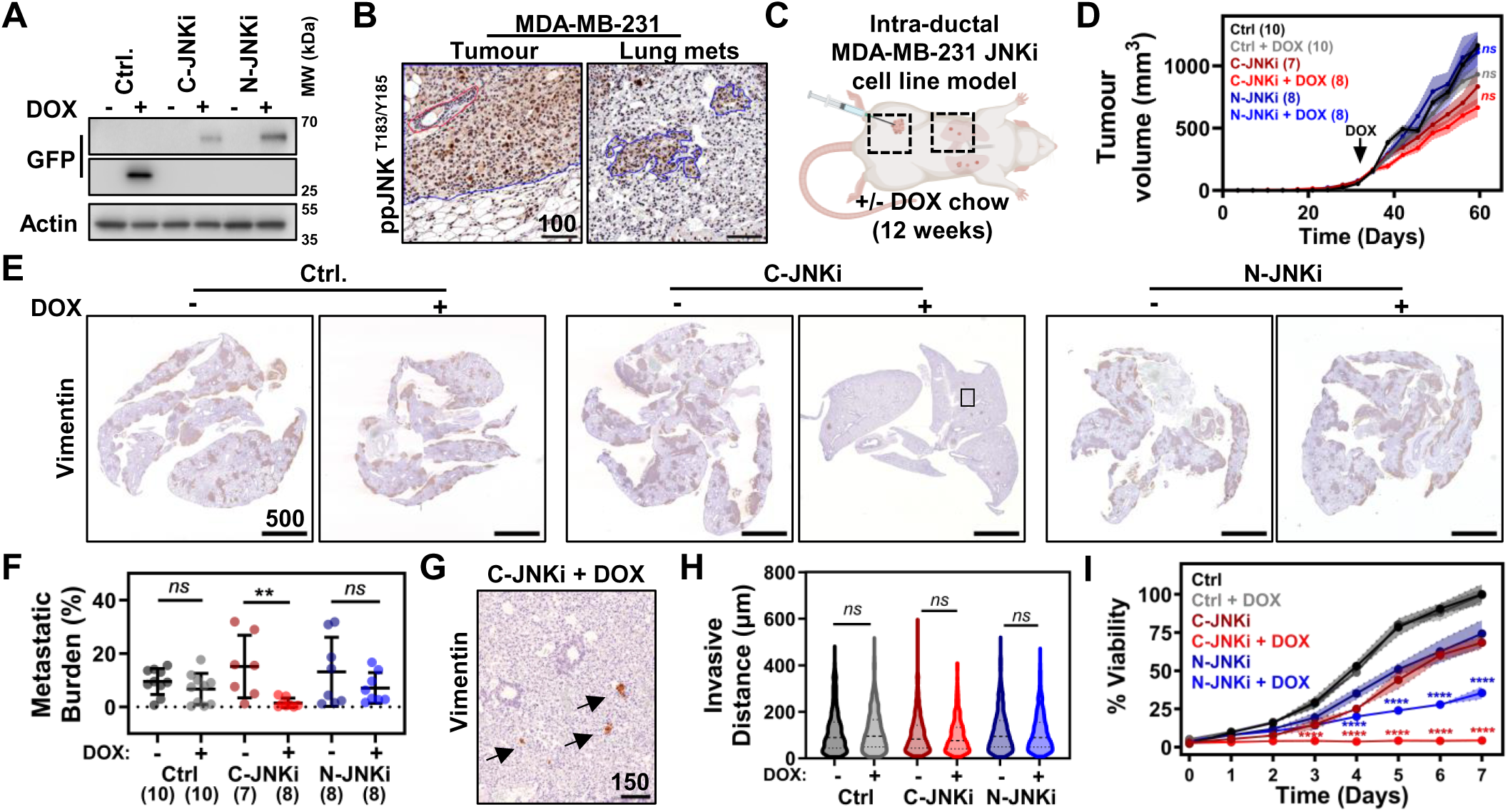
Cytoplasmic JNK activity drives TNBC metastatic outgrowth. **(A)** Representative Western blots of MDA-MB-231 cell lines stably expressing the GFP-fused control (Ctrl), cytoplasmic JNK inhibitor (C-JNKi) and nuclear JNK inhibitor (N-JNKi) constructs ± doxycycline. **(B)** Representative images of ppJNK staining within MDA-MB-231 primary tumour and lung metastases (mets). **(C)** Schematic representation of intra-ductal MDA-MB-231 mouse model in NSG mice with stable, subcellular JNK inhibitor-expressing cell lines. **(D)** Growth curves of primary tumours in mice receiving Ctrl, C-JNKi or N-JNKi MDA-MB-231 cells, fed normal or doxycycline chow (mean ± SD, n=7-10 as indicated; Kruskal-Wallis with Dunns post-test). **(E)** Representative images of vimentin-stained lung metastases from each experimental arm. **(F)** Assessment of metastatic burden (vimentin positive area/lung area, mean ± SD, n=7-10 as indicated; Kruskal-Wallis with Dunns post-test). **(G)** Representative image of vimentin-stained lung tissue from C-JNKi MDA-MB-231 recipient mouse, fed doxycycline chow. Image corresponds to the boxed area in panel E; arrows indicate micro-metastases. **(H)** Invasive distance of Ctrl, C-JNKi or N-JNKi MDA-MB-231 cells ± doxycycline in an organotypic invasion assay. Single-cell data are from a single representative experiment comprising 3 technical replicates per condition (Kruskal-Wallis with Dunn’s post-test). **(I)** Growth curves of Ctrl, C-JNKi or N-JNKi MDA-MB-231 cells cultured in 3D collagen matrices ± doxycycline. Data are from a single representative experiment with 6 technical replicates per condition, expressed relative to the Ctrl - doxycycline arm at day 7 (mean ± SD; mixed effects analysis, Tukey post-test).

To identify appropriate *in vitro* models to perform focused functional assays, we assessed the subcellular activity of JNK across a panel of breast cancer cell lines. Surprisingly, subcellular fractionation experiments showed that all breast cancer cell lines displayed only cytoplasmic JNK activity when grown on 2D-tissue culture plastic (**Fig. S2A**). However, the subtype-specific subcellular JNK activity profiles observed in patients could be recapitulated when these cell lines were cultured in 3D-collagen matrices, allowing the assembly of a cell line panel with varying levels of both cytoplasmic and nuclear JNK activity (**Fig. S2B**). This finding has important implications for all studies of JNK signalling and as such prompted us to perform functional assays within a 3D context. Therefore, using an organotypic matrix invasion assay, we could show that whilst the C-JNKi had no effect on MDA-MB-231 invasion (**Fig. 3H**), it abolished proliferation in 3D-collagen matrices (**Fig. 3I**). As expected, the C-JNKi had no effect on MCF7 proliferation, as these estrogen receptor (ER+) cells predominantly display nuclear JNK activity, with low cytoplasmic JNK staining in 3D-collagen matrices (**Fig. S2B-D**).

### Delineating the downstream functions of cytoplasmic JNK

To delineate the mechanisms downstream of cytoplasmic JNK in TNBC cells, we performed sequential phosphoproteomic and RNA-seq analyses on 3D cultures of MDA-MB-231 cell lines stably expressing the doxycycline-inducible Ctrl, C-JNKi or N-JNKi constructs (**Fig. 4A**). Whilst phosphoproteomic analyses revealed limited effects of the Ctrl and N-JNKi constructs on the MDA-MB-231 phosphoproteome, expression of the C-JNKi profoundly altered the phospho-landscape of these cells (**Fig. 4B**). To visualize this cytoplasmic JNK signalling network, the cellular localization (**Fig. S3A**) and function of each uniquely deregulated phosphoprotein were annotated from multiple online databases into a curated network map (**Fig. 4C**). Data were overlaid with the significantly deregulated phosphosites and potential direct substrates of cytoplasmic JNK (down-regulated sites with a putative JNK phosphorylation motif). This analysis highlights the breadth of JNK phospho-regulation across multiple cellular pathways, with phosphoproteins enriched in processes such as cytoskeletal regulation (24.2%), transcription (11.9%), RNA processing and biogenesis (9.6%), DNA/chromatin dynamics (8.2%), signal transduction (5.5%), translation (5.5%) and protein processing (5%). Along with these processes, phosphoproteins involved in apoptosis, pyruvate metabolism, nuclear membrane integrity and function, and autophagy were also enriched in this dataset, while all were significantly enriched with automated KEGG pathway and Gene Ontology analyses (**Fig. 4C**, **Fig. S3B-C**).

**Figure 4.**
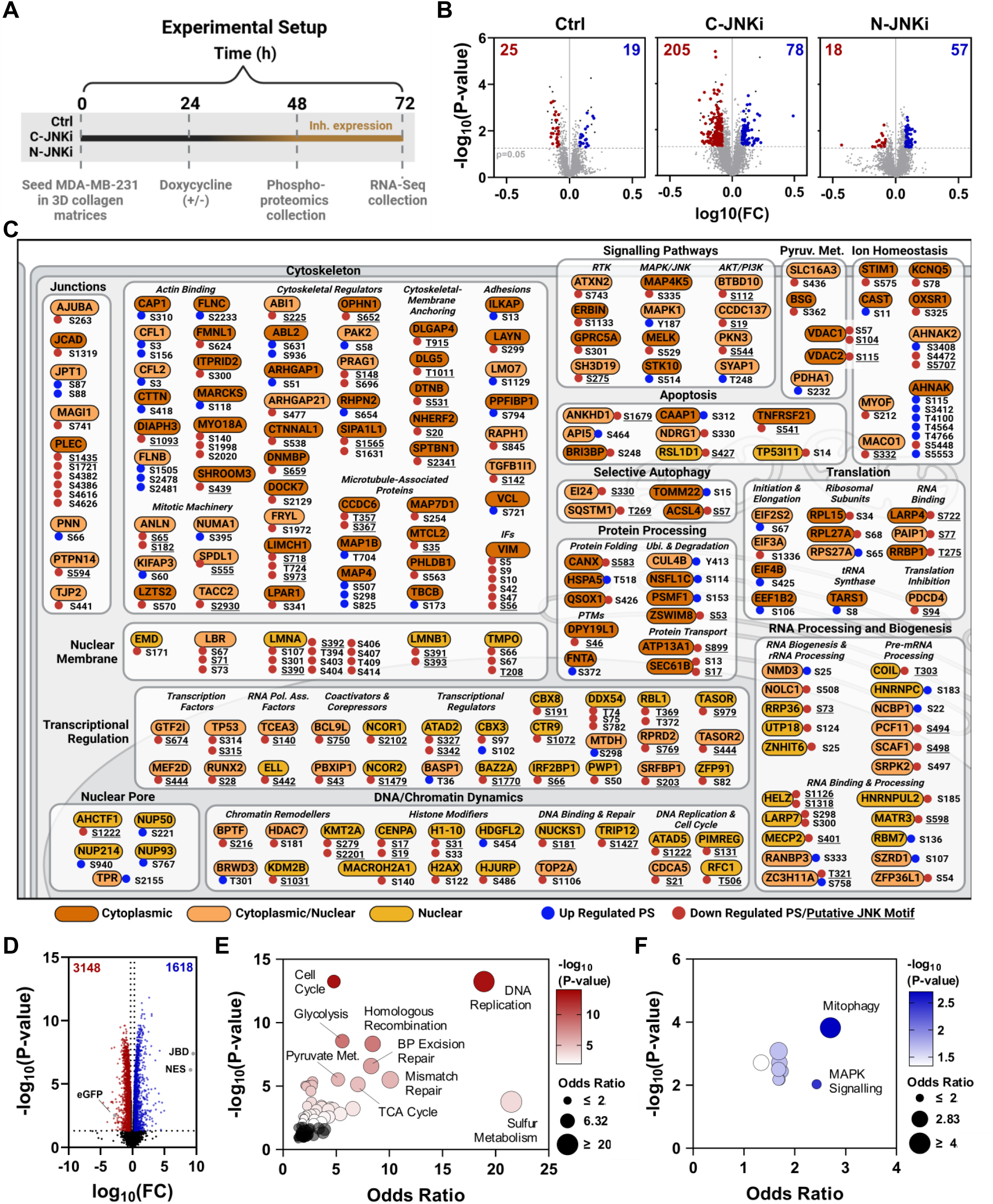
Phosphoproteomics and RNA-Seq analysis reveal downstream targets of cytoplasmic JNK. **(A)** Schematic of experimental setup used to identify downstream targets of cytoplasmic JNK signalling. Phosphoproteomics and RNA-Seq analyses of 3D cultures of MDA-MB-231 cell lines stably expressing the control (Ctrl), cytoplasmic JNK inhibitor (C-JNKi) and nuclear JNK inhibitor (N-JNKi) constructs ± doxycycline. **(B)** Volcano plots of the significantly deregulated phosphoproteins in each dataset (p-value <0.05, fold change (FC) >1.2, localisation probability >0.75). Significantly down-regulated (red) and up regulated (blue) phosphosites unique to each construct are depicted by colour. **(C)** Curated network map of the phosphosites significantly and uniquely deregulated by cytoplasmic JNK inhibition (91.16% coverage of dataset). Protein localisation was annotated from online databases and literature as cytoplasmic (brown), cytoplasmic and nuclear (orange), or nuclear (yellow). Phosphosites are represented as circles on the proteins, with an increase (blue) or decrease (red) in abundance depicted by colour. Significantly down-regulated phosphosites with a putative JNK phosphorylation motif (SP, TP or PxSP) are underlined. **(D)** Volcano plots of genes significantly deregulated by cytoplasmic JNK inhibition in RNA-Seq analysis (C-JNKi vs C-JNKi + doxycycline, corrected for Ctrl vs Ctrl + doxycycline; p-value <0.05, fold change >2). Balloon plot showing KEGG pathways associated with **(E)** significantly down-regulated genes and **(F)** significantly up-regulated genes.

In line with the canonical role of JNK in regulating gene expression through AP-1 transcription factors and the strong representation of transcriptional regulators in the C-JNKi phosphoproteomic dataset, RNA-seq analyses yielded 4766 genes significantly deregulated by C-JNKi expression (**Fig. 4D**). KEGG pathway analysis revealed that significantly down-regulated genes (**Fig. 4E**) are associated with DNA replication and repair, cell cycle and metabolism (glycolysis, pyruvate metabolism, TCA cycle and sulphur metabolism), whilst up-regulated genes are implicated in mitophagy and MAPK signalling (**Fig. 4F**). Together, these datasets suggest that cytoplasmic JNK mediates its oncogenic effects through multiple downstream mechanisms, with cytoskeletal regulation, DNA/chromatin dynamics, metabolism, selective autophagy, and signal transduction regulated at both transcriptional and post-translational levels.

### Identifying inhibitors of cytoplasmic JNK signalling through actin-based phenotypic screening

Given the prominent effect of cytoplasmic JNK inhibition on cytoskeletal proteins in phosphoproteomic analyses, immunofluorescence studies were undertaken to investigate the effect of C-JNKi expression on filamentous actin (F-actin) organisation. These analyses revealed drastic morphological changes in MDA-MB-231 cells expressing the C-JNKi, characterised by a significant reduction in cell size and a strong increase in cortical F-actin staining (**Fig. 5A**); observations supported by quantitative image analysis (**Fig. 5Bi**) and blinded scoring (**Fig. 5Bii, Fig. S4A**). This is consistent with the well reported role of JNK in cytoskeletal regulation, including studies undertaken in neurons using these inhibitory constructs^21,22^. Therefore, we next used this distinct morphological phenotype to interrogate a high-throughput actin-based drug screen of 114,400 structurally diverse compounds performed in SK-N-SH neuroblastoma cells^23^, in order to identify selective inhibitors of cytoplasmic JNK signalling. By first retrofitting the C-JNKi phenotype in MDA-MB-231 TNBC cells to this existing dataset through sampling of 18 compounds from 6 candidate clusters, we identified phenotypes comparable to the C-JNKi and assessed their effects on MDA-MB-231 morphology and F-actin organisation (**Fig. S4B**). These analyses revealed consistent overlap between the C-JNKi and compounds from clusters 2, 4 and 5 (**Fig. S4C-D**). As such, the 256 small molecules from these clusters were selected for further refinement screens to identify potent inhibitors of cytoplasmic JNK signalling (**Fig. 5C**, Stages 3-8).

**Figure 5.**
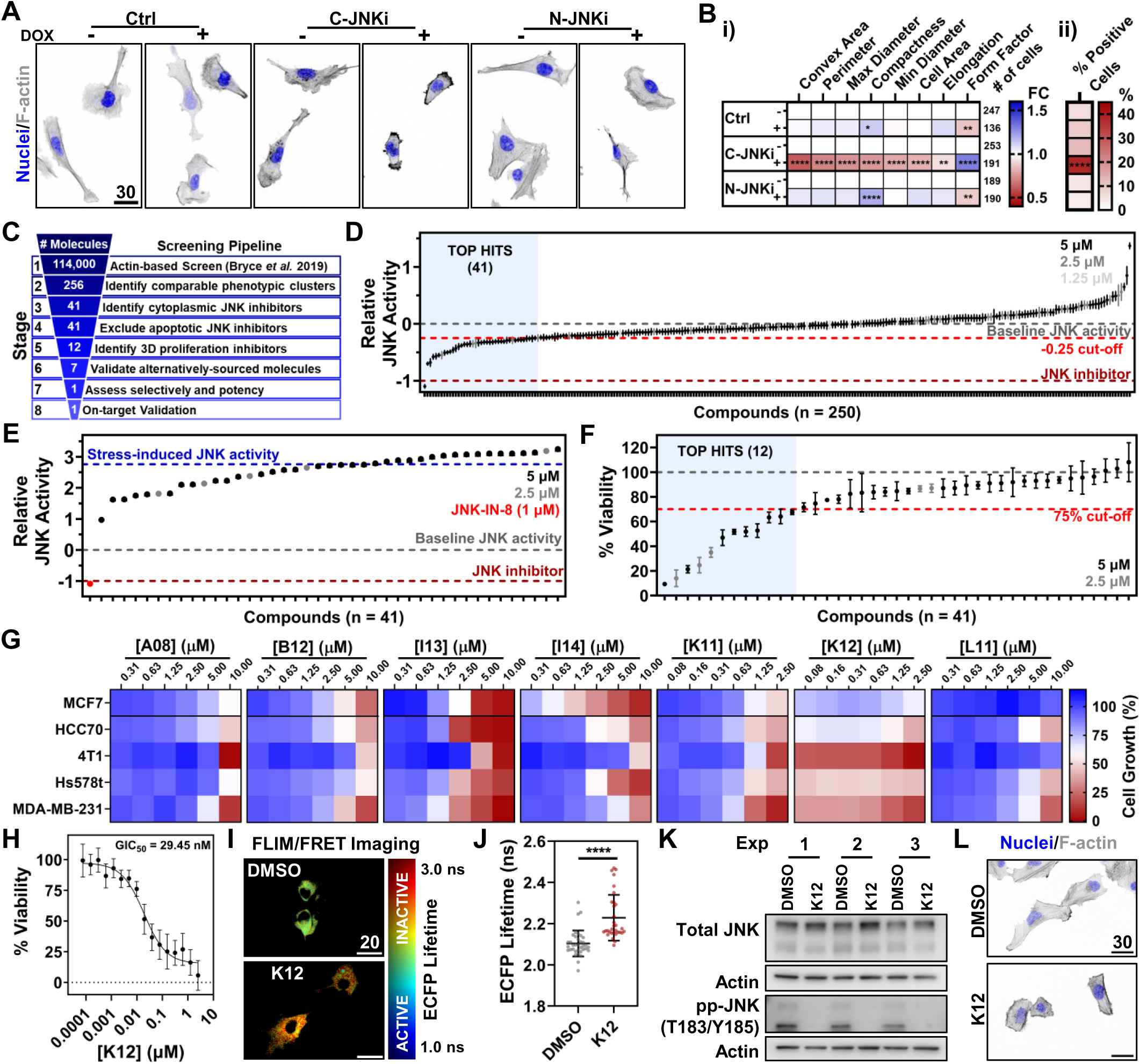
Identifying selective oncogenic JNK inhibitors through actin-based phenotypic screening. **(A)** Phalloidin staining of F-actin structures in MDA-MB-231 cell lines stably expressing the control (Ctrl), cytoplasmic JNK inhibitor (C-JNKi) and nuclear JNK inhibitor (N-JNKi) constructs ± doxycycline. **(B)** Summary of the cell morphology parameters significantly altered in C-JNKi expressing MDA-MB-231 cells treated with doxycycline. i) Quantitative analysis of morphological parameters assessed with CellProfiler 4.1.3., relative to the corresponding untreated (no doxycycline) control. Normalised data are pooled from 3 experiments, with total cell counts indicated to the right of the heatmap. ii) Blinded scoring of the phenotype enriched in C-JNKi expressing MDA-MB-231 cells treated with doxycycline from 3 independent experiments (small cells with cortical actin enrichment). **(C)** Schematic of the experimental pipeline used to identify selective cytoplasmic JNK inhibitors from within Bryce *et al.* phenotypic clusters 2, 4 and 5. **(D)** High content imaging of MDA-MB-231 JNK-KTR mClover cells following 24 h treatment with 256 compounds at doses ranging from 1.25-5 µM. Relative JNK activity (JNK-KTR cytoplasmic/nuclear ratio) was calculated from single cell data obtained from 3 technical replicates, normalized to DMSO and JNK-IN-8 controls (mean ± 95% CI. Data for 6 compounds were excluded due to biosensor mislocalisation. **(E)** High content imaging of MDA-MB-231 JNK-KTR mClover cells treated for 24 h with 2.5-5 µM of the 41 top hits, followed by anisomycin treatment (300 nM, 30 min) (mean ± 95% CI). **(F)** Viability of MDA-MB-231 cells in 3D collagen matrices treated for 72 h with 2.5-5 µM of the top 41 compounds. Data are from a representative experiment with 3 technical replicates per compound, normalised to the DMSO control (mean ± SD). **(G)** MCF7 (ER+), HCC70 (TNBC), 4T1 (murine TNBC), Hs578t (TNBC) and MDA-MB-231 (TNBC) breast cancer cell lines cultured in 3D collagen matrices were treated for 7 days with the 7 top hits at doses ranging from 0.08-10 µM, as indicated. Data shown are pooled from 3 independent experiments, relative to the respective untreated control. **(H)** MDA-MB-231 cells cultured in 3D collagen matrices were treated for 7 days with K12 doses ranging from 0.08 nM – 2.5 µM. Data shown are from a representative experiment comprising 6 technical replicates for each dose tested (Mean ± SD). **(I)** Representative FLIM images of MDA-MB-231 cells stably expressing the JNK-AR1-EV NES FRET biosensor cultured in 3D collagen matrices and treated with DMSO or K12 (2.5 µM) for 24 h. **(J)** Quantitative analysis of FLIM/FRET imaging from a representative experiment showing single cell data from DMSO (36 cells) and K12 (32 cells) treated 3D cultures (Mann-Whitney test). **(K)** Western blotting of MDA-MB-231 cell lines cultured in 3D collagen matrices treated with DMSO or K12 (2.5 µM) for 72 h. Samples are from 3 independent experiments (Exp 1-3). **(L)** Phalloidin staining of F-actin structures in MDA-MB-231 cell lines treated with DMSO or K12 (2.5 µM) for 24 h. In all cases: *p<0.05, ** p<0.01 and **** p<0.0001. All scale bars are in µm.

Exploiting our finding that breast cancer cells exclusively display cytoplasmic JNK activity when grown on 2D-tissue culture plastic (**Fig. S2A**), we utilised the JNK-Kinase Translocation Reporter^24^ (JNK-KTR, **Fig. S5A**) to measure single cell JNK activity by high content imaging (**Fig. S5B**) and screen for inhibitors of cytoplasmic JNK signalling. This approach yielded 41 compounds that inhibited cytoplasmic JNK activity below a cut-off of −0.25 (**Fig. 5D**). By treating these JNK-KTR MDA-MB-231 cells with anisomycin, a protein synthesis inhibitor that potently activates stress-induced JNK signalling, we subsequently demonstrated that none of the 41 compounds tested reduced stress-induced JNK signalling below baseline, as seen with the direct JNK inhibitor, JNK-IN-8 (**Fig. 5E**). Given our observation that cytoplasmic JNK inhibition suppressed proliferation in 3D-matrices, each compound was screened through 3D-collagen growth assays, yielding 12 compounds that reduced proliferation below a cut-off of 75% following 72 h treatment (**Fig. 5F**). Of these, the 11 compounds that could be sourced commercially were rescreened through stages 3-5 of our analysis pipeline (**Fig. S5C-E**), revealing 7 compounds that satisfied each assay. These compounds, nominally named A08, B11, I13, I14, K11, K12 and L11, were evaluated in 3D-proliferation assays across our breast cancer cell line panel (**Fig. 5G**). These analyses yielded K12 as the compound with the greatest potency and specificity for cell lines with cytoplasmic JNK activity, as demonstrated by a reduced effect in MCF7 cells, which have high nuclear and low cytoplasmic JNK activity in 3D collagen matrices (**Fig S2B**). Follow-up time course studies demonstrated a dose-dependent inhibitory effect of K12 on MDA-MB-231 3D-proliferation (**Fig. S5F**), with a half maximal Growth Inhibitory concentration (GIC_50_) value of ∼30 nM determined 7 days post treatment (**Fig. 5H**).

For orthogonal, on-target validation, we harnessed a JNK activity FRET biosensor (JNK-AR1-EV NES; **Fig. S5G**) and performed FLIM/FRET imaging of 3D-cultures to demonstrate that 24 h K12 treatment significantly increases eCFP (donor) fluorescence lifetime and thus significantly decreases JNK activity (**Fig. 5I-J**). Furthermore, 3D cultures showed a significant reduction in ppJNK levels following K12 treatment (**Fig. 5K, Fig. S5H**), whilst cellular and actin phenotypes induced by K12 treatment in 2D cultures parallel those driven by the C-JNKi (**Fig. 5L, Fig. S5I-J**). Together, these data demonstrate that K12 is a potent inhibitor of cytoplasmic JNK activity in TNBC cells, warranting further *in vivo* investigation as a metastasis-targeting agent.

### K12 selectively inhibits cytoplasmic JNK activity *in vivo* and blocks metastatic outgrowth

For the *in vivo* assessment of K12, maximum tolerated dose (MTD) studies were undertaken, with clinical signs and mortality evaluated in Swiss Albino mice for 72 h following single dose intraperitoneal (i.p.) administration. These analyses yielded an MTD of ≥3 mg/kg body weight. Further exposure studies revealed a dose-dependent increase in K12 plasma concentration following i.p. administration (**Fig. 6A**), with a peak plasma concentration (C_max_) of 1308.6 ng/mL and terminal half-life (T½) of 1.33 h following administration of K12 at 3 mg/kg. To validate the on-target effects of K12 *in vivo*, MDA-MB-231 JNK-AR1-NES cells were intravenously (i.v.) injected into NSG mice and allowed to colonise the lungs for 4 days prior to single-dose K12 treatment (3 mg/kg) and FLIM/FRET imaging of *ex vivo* lung tissue (**Fig. 6B**). These analyses revealed a significant increase in eCFP (donor) fluorescence lifetime 48 h after treatment (**Fig. 6C-D**), demonstrating that K12 is effective at inhibiting cytoplasmic JNK activity within disseminated TNBC cells *in vivo*.

**Figure 6.**
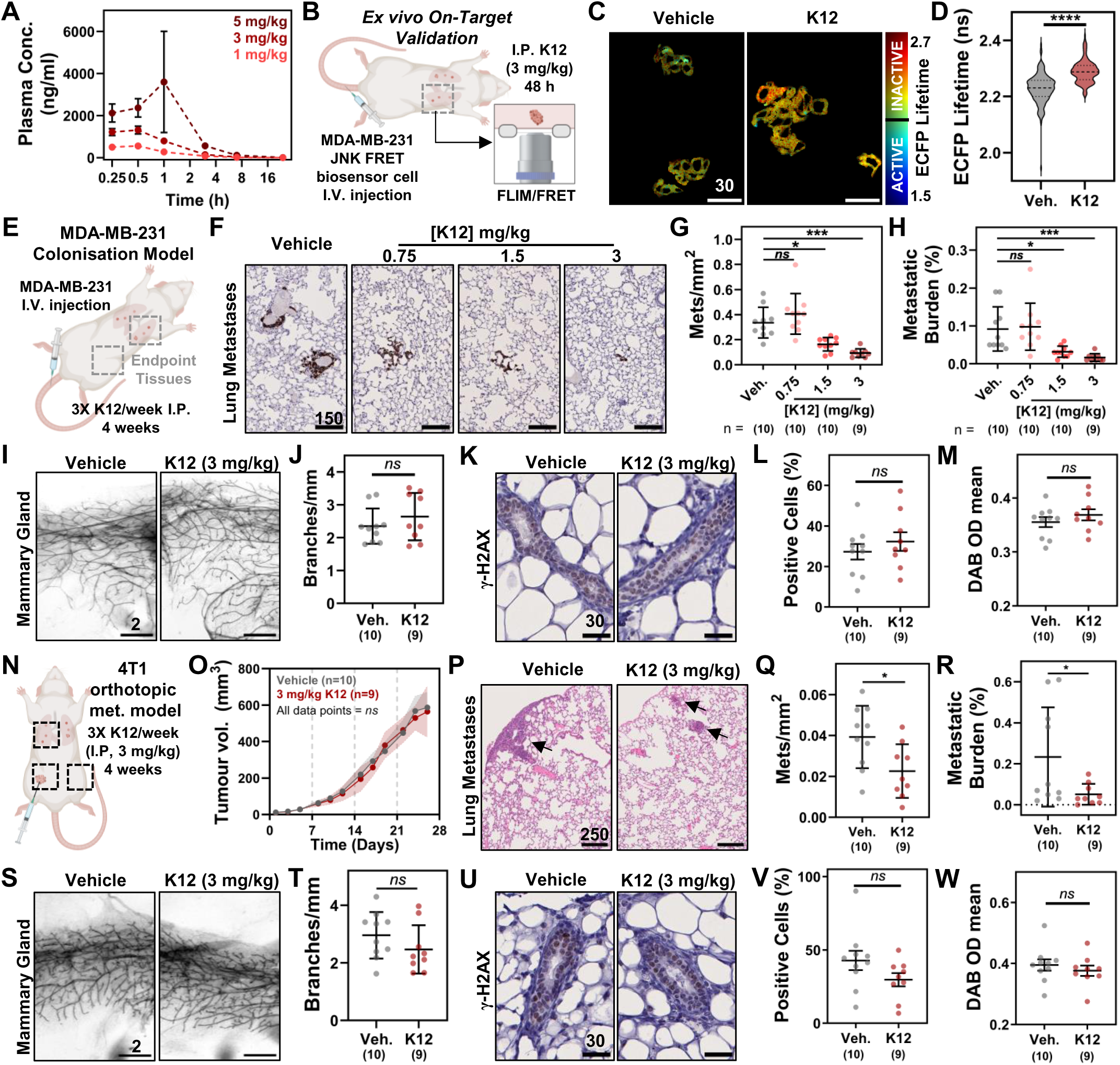
K12 selectively inhibits cytoplasmic JNK activity in disseminated TNBC cells and blocks metastatic outgrowth. **(A)** Plasma profile of K12 in Swiss albino mice following single intraperitoneal (I.P.) administration at 1, 3 or 5 mg/kg (mean ± SD, n=3). **(B)** Schematic of *ex vivo* FLIM/FRET imaging experiment used to validate on-target effects of K12 in disseminated MDA-MB-231 cells expressing the JNK FRET (JNK-AR1-EV NES) biosensor. **(C)** Representative *ex vivo* FLIM images of disseminated MDA-MB-231 tumour cells in NSG lung tissue, 48 h after I.P. administration of vehicle or K12 (3 mg/kg). **(D)** Quantitative analysis of *ex vivo* single-cell FLIM data from vehicle (n=171 cells) and K12 (3 mg/kg; n=177) treated mice (Mann-Whitney test). **(E)** Schematic representation of the MDA-MB-231 colonisation mouse model performed in NSG mice to evaluate K12 efficacy. **(F)** Representative images of vimentin-stained lung tissue from mice treated 3 times a week, for 4 weeks, with vehicle or K12 (0.75, 1.5 or 3 mg/kg). **(G)** Quantitative assessment of lung metastases (mean ± SD, n=9-10, as indicated). **(H)** Quantitative analysis of metastatic burden (mean ± SD, n=9-10, as indicated). **(I)** Representative images of carmine-stained mammary glands. **(J)** Quantitative assessment of mammary gland branching (mean ± SD, n=10 vehicle/n=9 K12; t-test). **(K)** Representative images of γ-H2AX staining in mammary gland cross-sections from vehicle and K12 (3 mg/kg) treated mice. Quantitative assessment of γ-H2AX staining within the mammary gland luminal epithelial layer (3 mg/kg; mean ± SEM, n=10 vehicle/n=9 K12; t-test) by **(L)** positive pixel analysis and **(M)** staining intensity. **(N)** Schematic representation of the 4T1 orthotopic model performed in BALB/c mice to validate K12 efficacy. **(O)** Tumour volume in mice treated with vehicle or K12 (3 mg/kg) 3 times a week, for 4 weeks (mean ± SD, n=10 vehicle/n=9 K12; mixed effects analysis, Sidaks post-test). **(P)** Representative images of H&E-stained lung tissue. Arrows indicate metastases. **(Q)** Quantitative assessment of lung metastases (mean ± SD, n=10 vehicle/n=9 K12; t-test). **(R)** Quantitative analysis of metastatic burden (mean ± SD, n=10 vehicle/n=9 K12; t-test). **(S)** Representative images of carmine-stained mammary glands. **(T)** Quantitative assessment of the number of branch points per mm of mammary gland (mean ± SD, 10 vehicle/9 K12 mice; t-test). **(U)** Representative images of γ-H2AX staining in mammary gland cross-sections. Quantitative assessment of γ-H2AX staining within the mammary gland luminal epithelial layer by **(V)** positive pixel analysis and **(W)** staining intensity (mean ± SEM, 10 vehicle/9 K12 mice; t-test).

To assess the efficacy of K12 for inhibiting TNBC metastatic outgrowth, MDA-MB-231 cells were i.v. injected into NSG mice 4 days prior to the commencement of treatment with vehicle or K12 (0.75, 1.5 or 3 mg/kg), three times a week for 4 weeks (**Fig. 6E**). Of note, no signs of treatment-induced toxicity were observed throughout the experiment (**Fig. S6A,** weight loss over time), or in histopathological analyses of kidney, spleen and liver samples at experimental endpoint (**Fig. S6B-D**). IHC analysis of lung tissue (**Fig. 6F**) did however reveal a dose-dependent effect of K12 on the number of lung metastases and metastatic burden within the lungs (**Fig. 6G-H**). To confirm the selectivity of K12 for oncogenic JNK activity, an analysis of branching and DNA damage was undertaken on mammary glands from mice treated with vehicle and 3 mg/kg K12. Unlike the direct JNK inhibitor CC-401, K12 treatment had no effect on the number of mammary gland branches (**Fig. 6I-J**), or the frequency or intensity of γH2AX staining (**Fig. 6K-M**), confirming that this compound does not impact the tumour suppressive functions of JNK.

Using the 4T1 intra-mammary model as an orthogonal approach for assessing K12 *in vivo* efficacy and selectivity in an immune-competent setting (**Fig. 6N**), we again observed no signs of K12-induced weight loss during the treatment course (**Fig. S6E**). In line with our previous assessment of cytoplasmic JNK inhibition *in vivo* (**Fig. 2**, **Fig. 3**), K12 had no effect on primary tumour growth (**Fig. 6O**) or end-point tumour burden (**Fig. S6F**) but significantly inhibited the number of lung metastases and overall metastatic burden within the lungs (**Fig. 6P-R**). Consistent with the MDA-MB-231 model, K12 elicited no effect on mammary gland branching (**Fig. 6S-T**) or γH2AX staining (**Fig. 6U-W**) in this immune-competent model. Together, these data demonstrate that K12 is a selective inhibitor of cytoplasmic, oncogenic JNK activity and effective at blocking metastatic TNBC progression *in vivo*.

### Target identification studies reveal glutaminase-1 to be a direct target of K12

As K12 (**Fig. 7A**) was identified through phenotypic drug screening, its direct protein target(s) were unknown. To resolve this, kinome profiling was performed using the Eurofins DiscoverX KINOME*scan* active site-directed competition binding assay. This evaluation of a single high dose treatment of K12 (2.5 μM) across 468 kinases revealed that K12 is not a direct JNK inhibitor, nor does it mediate its effects through any kinase tested (**Fig. S7**). Turning to affinity purification-mass spectrometry (AP-MS) for target identification, medicinal chemistry was used to generate a K12 affinity probe. For this, K12 (**Fig. S8A**) and a series of structural analogues (**Fig. S8B**) were synthesised and screened through 3D proliferation assays (**Fig. S8C**), revealing the C5-position of the thiophene ring as the optimal site for sepharose bead conjugation (**Fig. S8D**). Through AP-MS, we identified 8 potential protein targets that were both significantly enriched on K12 vs negative control beads and also competed off the K12 beads by excess free K12 (**Fig. 7B-C**). Targets were validated using siRNA-mediated knockdown in our established MDA-MB-231 JNK-KTR high content imaging and 3D-growth assays (**Fig. 7D-E**), which uncovered glutaminase-1 (GLS) as a likely functional target of K12, given its ability to regulate cytoplasmic JNK activity in 2D cultures as well as proliferation in 3D matrices.

**Figure 7.**
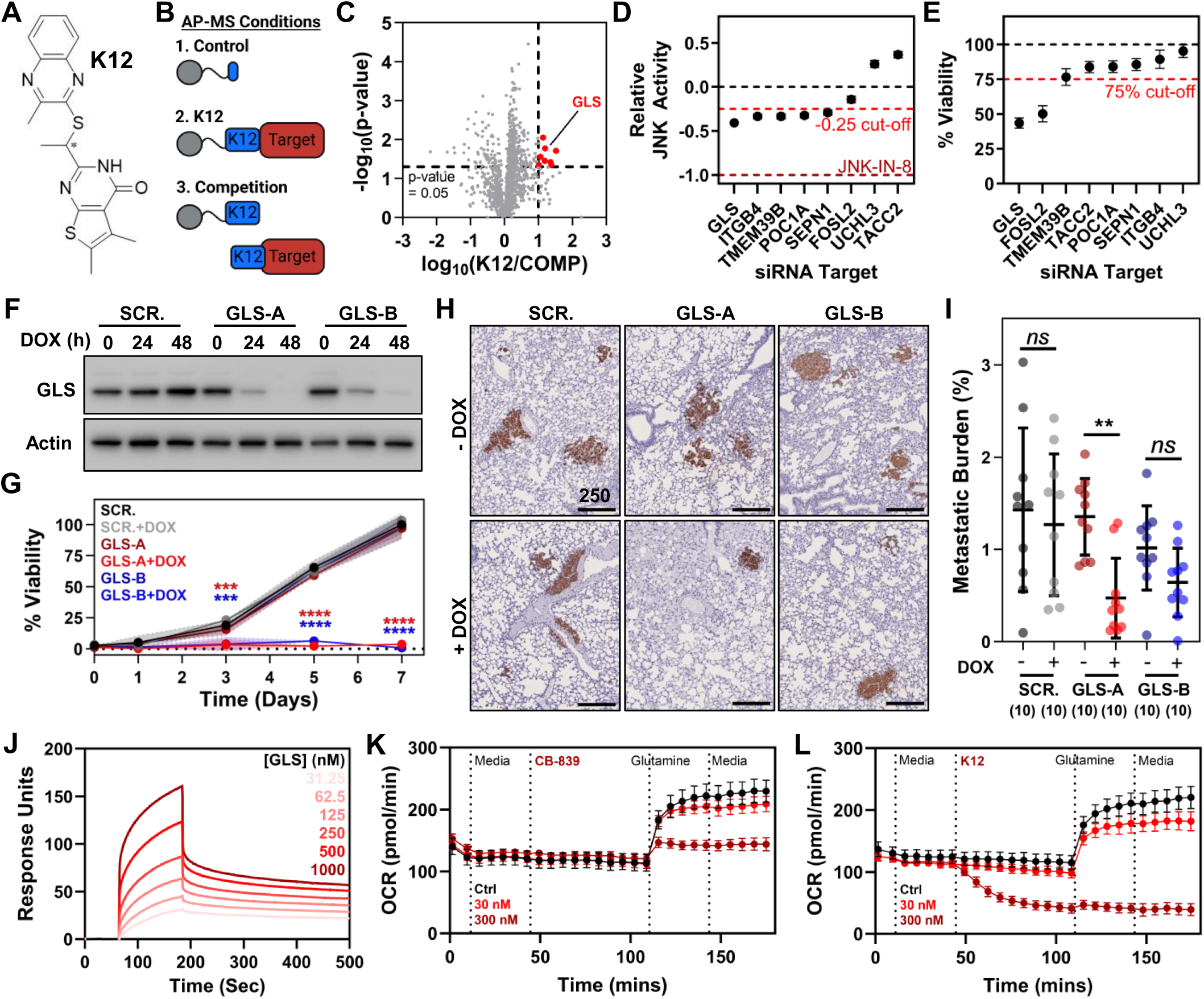
Glutaminase 1 is a direct target of K12. **(A)** Chemical structure of K12 (stereocenter indicated by *). **(B)** Schematic outlining the 3 experimental arms used for affinity purification-mass spectrometry (AP-MS) analysis. **(C)** Volcano plot of affinity-mass spectrometry analysis. Data were initially filtered for proteins significantly enriched on K12 beads vs control beads (p<0.05). Data presented depict the selective removal of proteins bound to K12 beads with excess free K12 (300 nM). Proteins of interest with log_2_ fold change > 1 and p-value < 0.05 are shown in red. **(D)** High content imaging of MDA-MB-231 JNK-KTR mClover cells treated for 72 h with the siRNA (15 nM) indicated. Relative JNK activity (JNK-KTR cytoplasmic/nuclear ratio) was calculated from single-cell data obtained from 3 technical replicates, normalised to negative control siRNA and JNK-IN-8 controls (mean ± 95% CI). **(E)** Viability of MDA-MB-231 cells treated for 24 h with the siRNA (10 nM) indicated and cultured for a further 72 h in 3D collagen matrices. Data are from a representative experiment with 6 technical replicates, normalised to the negative control siRNA condition (mean ± SD). **(F)** Western blot validation of MDA-MB-231 cell lines stably expressing doxycycline-inducible negative control (SCR.) or GLS targeting (GLS-A and GLS-B) shRNA. **(G)** Growth curves of these shRNA expressing cell lines cultured in 3D collagen matrices ± doxycycline. Data are from a representative experiment, expressed relative to the SCR. - doxycycline control at day 7 (mean ± SD; mixed effects analysis, Tukey post-test). **(H)** Representative images of vimentin-stained lung tissue from mice tail-vein injected with these shRNA expressing cell lines, followed by normal or doxycycline chow for 4 weeks post-injection. **(I)** Assessment of metastatic burden (vimentin positive area/lung area, mean ± SD, n=10; non-parametric test, one-way ANOVA with Sidaks post-test). **(J)** Surface plasmon resonance sensorgram showing the dose-dependent binding of glutaminase 1 (31.25-1000 nM) to immobilised K12. **(K)** Seahorse bioanalyzer assessment of oxygen consumption rates (OCR) in MDA-MB-231 cells treated with CB-839 (30 nM and 300 nM) followed by glutamine (2 mM). **(L)** Seahorse bioanalyzer assessment of oxygen consumption rates in MDA-MB-231 treated with K12 (30 nM and 300 nM) followed by glutamine (2 mM).

GLS is a mitochondrial enzyme that hydrolyses glutamine to glutamate. TNBC tumours are known to be addicted to glutamine metabolism^25^, with the upregulation of GLS and a number of related enzymes and transporters already observed in TNBC^26^. To validate GLS as a direct target of K12, MDA-MB-231 cell lines stably expressing doxycycline-inducible negative control (SCR.) and GLS-targeting (GLS-A and GLS-B) shRNA were generated. Western blot analyses showed that it is the low molecular weight GAC isoform of GLS that is abundant in MDA-MB-231 cells, consistent with literature examining TNBC cells and tumours^27,28^, and that the protein was efficiently knocked down with doxycycline treatment of GLS-A and GLS-B shRNA expressing cells (**Fig. 7F**). Whilst GLS depletion recapitulated the inhibitory effects of K12 and the C-JNKi in 3D-growth assays (**Fig. 7G**), a less striking effect was observed within an *in vivo* colonisation assay, with a significant decrease in metastatic burden only observed in MDA-MB-231 expressing the GLS-A shRNA construct (**Fig. 7H-I**). To further validate a direct interaction between GLS and K12, surface plasmon resonance analysis with recombinant GLS was performed. Although model fitting with these data was complicated by the presence of the two K12 enantiomers, they do demonstrate direct and dose-dependent binding of GLS to K12 (**Fig. 7J**). These data are supported by *in silico* docking studies (**Fig. S9A-C**), which show binding of the K12-S enantiomer within an allosteric binding pocket of the GLS homo-tetramer. Importantly. this is also the binding site for CB-839^29^, a well-characterised GLS inhibitor that has progressed to Phase I/II clinical trials for the treatment of various tumours^30–32^.

On this basis, modified Seahorse XF assays were undertaken to confirm that K12 elicits an inhibitory effect on GLS-dependent metabolic processes. Through direct comparisons with CB-839 (**Fig. 7K**), it was apparent that low doses of K12 (30 nM) were as effective as equimolar doses of CB-839 (30 nM) at inhibiting glutamine-dependent oxygen consumption (**Fig. 7L**). However, these data also demonstrated that high doses of K12 (300 nM) had a markedly distinct effect from CB-839, inducing a rapid glutamine-independent decrease in oxygen consumption rate (**Fig. 7L**). These Seahorse data, along with the modest effects of GLS knockdown in MDA-MB-231 *in vivo* metastasis models, suggest that GLS inhibition alone does not account for the full anti-metastatic effects of K12 and that this compound is likely acting through additional protein targets.

### Dual targeting of glutaminase-1 and the pyruvate dehydrogenase complex by K12 overcomes pyruvate anaplerosis

To directly analyse glutaminase inhibition, we undertook ^13^C_5_-glutamine tracing in MDA-MB-231 cells incubated with DMSO, CB-839 or K12 for 6 h (**Fig. 8A**). Metabolomics analyses revealed that whilst CB-839 effectively blocks GLS-dependent glutamine hydrolysis, K12 elicits a modest but significant effect on GLS activity, as seen by a reduction in the ratio of m+5 glutamate/glutamine isotopologues (**Fig. 8B**) and a significant decrease in m+5 α-ketoglutarate (**Fig. S10A**). This effect was observed at all doses of K12 tested, indicating that GLS is potentially a high affinity target of K12. In support of this, lower levels of glutamine-derived carbons were metabolised from m+5 α-ketoglutarate into m+4 succinate during both CB-839 and K12 treatment (**Fig. S10B**), with this reduction also observed in m+4 fumarate and m+4 malate, suggesting that both drugs block glutamine entry into oxidative TCA cycle (**Fig. S10C-D**). Intriguingly though, there was a compensatory increase in m+3 fumarate and m+3 malate in K12 only, suggestive of reductive carboxylation (**Fig. 8A** and **Fig. S10C-D**). Indeed, analysis of citrate isotopologues confirmed that K12 induced reductive carboxylation, as indicated by a dose dependent increase in the ratio of reductive m+5 citrate compared to oxidative (TCA-derived) m+4 citrate (**Fig. 8C**). These data were not observed with CB-839, suggesting that K12 acts upon additional target(s) to CB-839 at higher doses.

**Figure 8.**
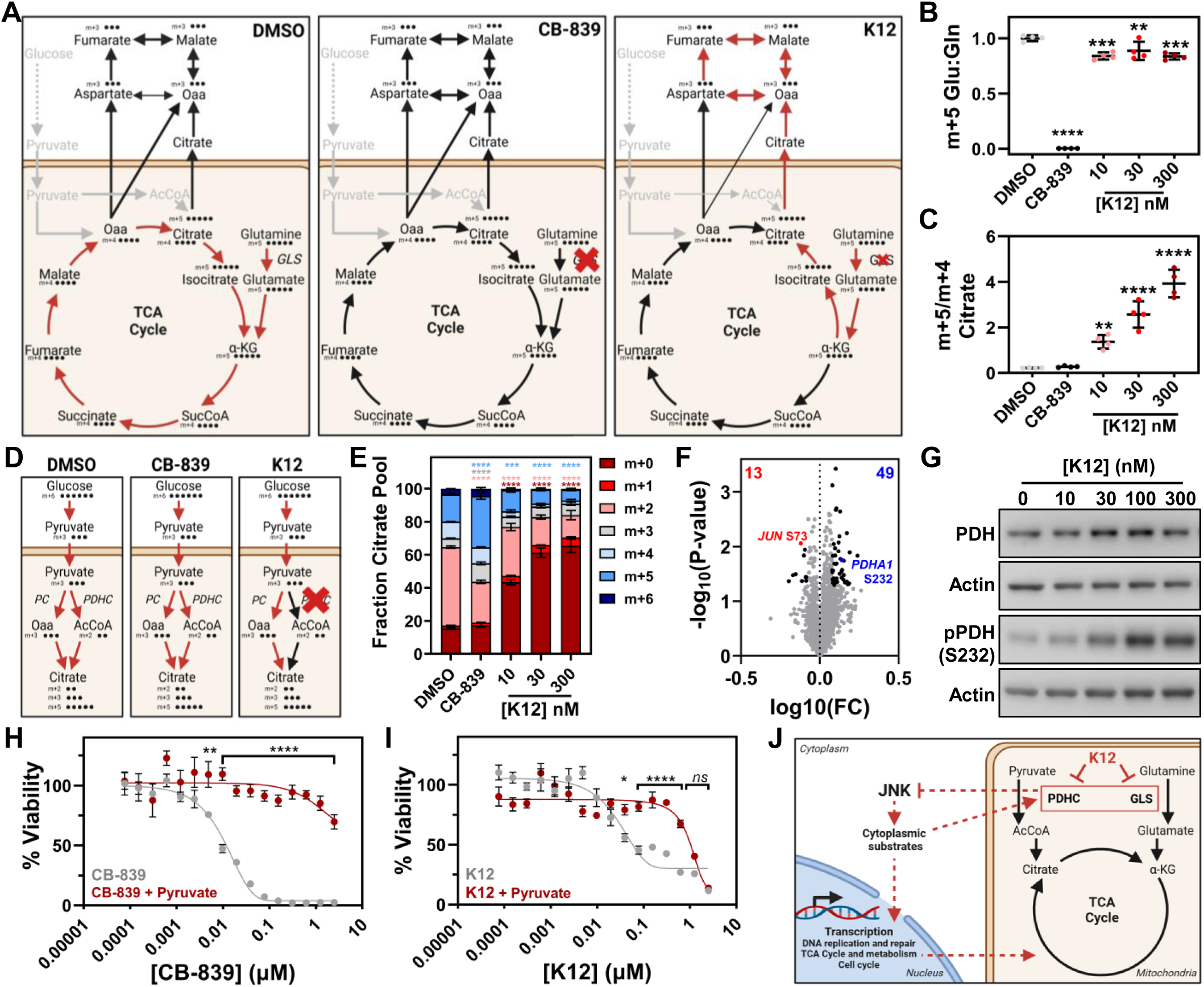
K12 overcomes pyruvate anaplerosis through low affinity effects on the pyruvate dehydrogenase complex. **(A)** Schematic representation of metabolomics studies in MDA-MB-231 cells depicting U-^13^C_5_-glutamine carbon transfer between TCA intermediates following treatment with DMSO, CB-839 (300 nM) or K12 (10, 30 and 300 nM). Red arrows depict prominent tracing patterns for each condition and red crosses depict relative GLS inhibition. **(B)** Ratio of m+5 glutamine to glutamate (Glu:Gln) in ^13^C_5_-glutamine tracing experiments as a readout of GLS activity (n=4, mean ± SD; one-way ANOVA with Dunnett’s post-test). **(C)** Ratio of m+5 citrate to m+4 citrate levels in U-^13^C_5_-glutamine tracing experiments as a readout of reductive carboxylation (n=4, mean ± SD; one-way ANOVA with Dunnett’s post-test). **(D)** Schematic representation of metabolomics studies in MDA-MB-231 cells depicting U-^13^C_6_-glucose carbon transfer, tracing glucose entry into the TCA cycle following treatment with DMSO, CB-839 (300 nM) or K12 (10, 30 and 300 nM). Red arrows depict prominent tracing patterns for each condition and the red cross depicts PDHC inhibition. **(E)** Effect of DMSO, CB-839 (300 nM) and K12 (10, 30 and 300 nM) on ^13^C fraction enrichment of citrate after 6 h incubation with U-^13^C_6_-glucose (n=4, mean ± SD; mixed effects analysis with Dunnett’s post-test). Unlabelled metabolite (^12^C) is denoted by m+0, whilst m+x denotes a metabolite with x number of carbons labelled with ^13^C. **(F)** Volcano plot of phosphoproteomics analyses performed with MDA-MB-231 cells following treatment with K12 (300 nM) for 24 h. Significantly down-regulated sites shown in red and up regulated sites shown in blue. **(G)** Western blotting of MDA-MB-231 cells treated with K12 (10, 30, 100 and 300 nM) for 1 h. **(H)** MDA-MB-231 cells cultured in 3D collagen matrices were treated for 7 days with increasing doses of CB-839 ± Na-Pyruvate (1 mM). Data shown are from a representative experiment comprising 6 technical replicates (mean ± SD; mixed effects analysis with Sidaks post-test). **(I)** MDA-MB-231 cells cultured in 3D collagen matrices were treated for 7 days with increasing doses of K12 ± Na-Pyruvate (1 mM). Data shown are from a representative experiment comprising 6 technical replicates (mean ± SD; mixed effects analysis with Sidaks post-test). **(J)** Schematic representation of the mechanism of K12.

One striking observation was that total citrate was depleted to extremely low levels with K12 treatment (**Fig. S10E**). Since glucose contributes substantially to citrate metabolite levels, we next undertook ^13^C_6_-glucose tracing in MDA-MB-231 cells incubated with DMSO, CB-839 or K12 for 6 h (**Fig. 8D**). Metabolomics analysis confirmed the effects of K12 on total citrate levels (**Fig. S10F**), while also revealing a clear dose dependent effect of K12 blocking m+2 and m+5 citrate production (**Fig. 8E**). The citrate m+2 isotopologue is generated via the PDHC (m+2 AcCoA), the m+3 via pyruvate carboxylase (m+3 Oaa) and the m+5 is generated by both pathways together (**Fig. 8D**). Therefore, the loss of citrate m+2 and m+5, with no change in m+3, suggests that K12 blocks PDHC activity.

PDHC activity is tightly regulated by phosphorylation of the pyruvate dehydrogenase (PDH) E1α subunit, with phosphorylation at Ser232, Ser293 or Ser300 rendering the complex inactive^33^. As our earlier phosphoproteomic analyses had identified a significant increase in PDH (*PDHA1*) Ser232 phosphorylation following cytoplasmic JNK inhibition (**Fig. 4C**), phosphoproteomics were repeated on vehicle vs K12 treated (300 nM) MDA-MB-231 3D cultures. Consistently, these results showed significantly increased PDH Ser232 phosphorylation following K12 treatment, along with a concomitant reduction in c-Jun Ser73 phosphorylation (**Fig. 8F**). Follow-up Western blot analyses further confirmed that this effect on PDH was a dose-dependent (**Fig. 8G, Fig. S10G**) and rapid effect of K12 (**Fig. S10H**), with maximal Ser232 phosphorylation observed within 30 min of treatment.

While the mechanism of PDHC phosphorylation and the direct target of K12 within the PDHC was not identified by this analysis, this dual targeting of K12 represents a highly serendipitous finding, as pyruvate anaplerosis is known to drive resistance to the GLS inhibitors CB-839 and BPTES through the replenishment of TCA cycle intermediates^34^. In support of this, our metabolomics data clearly showed that CB-839 treatment resulted in the conversion of more pyruvate into citrate (m+5 and m+3) compared to K12 or DMSO control (**Fig. 8E**). This effect becomes more evident in 3D-growth assays, where the inhibitory effect of CB-839 on MDA-MB-231 proliferation was significantly ablated following pyruvate supplementation (**Fig. 8H**). Remarkably, we show that high doses of K12 retained their anti-proliferative effects on MDA-MB-231 cells in the presence of pyruvate (**Fig. 8I**), indicating that PDHC targeting by K12 overcomes this known mechanism of resistance observed with existing GLS inhibitors.

Together, these data show that K12 directly inhibits GLS and rapidly blocks PDHC activity. Furthermore, the data in Figure 7D demonstrate that GLS depletion leads to decreased cytoplasmic JNK activity, which would in turn drive PDH Ser232 phosphorylation (**Fig. 4C**) and the downregulation of genes implicated in pyruvate metabolism, glycolysis and the TCA cycle (**Fig. 4D-E**). Collectively, this suggests that K12 treatment disrupts a feedback loop where cytoplasmic JNK activity drives a metabolic phenotype that in turn promotes the activation of further JNK activity (**Fig. 8J**), with the resulting metabolic blockade likely contributing to the *in vivo* impact of this novel oncogenic JNK inhibitor upon disseminated TNBC cells.

## Discussion

Whilst there has been significant interest in therapeutically targeting JNK signalling across various indications, including different cancers, direct JNK inhibitors have not progressed through clinical trials^17^. This is largely due to the pleiotropic roles of the JNK pathway and the multitude of downstream targets that JNK influences. In breast tissue alone, JNK has been characterised as tumour-suppressive^12–15,35^, tumour-promoting^9,36^, a driver of metastatic disease^8–11^, and an essential regulator of apoptosis-mediated drug response^16^. Our data from the 4T1 orthotopic model with the first generation JNK inhibitor CC-401 highlight these complexities and are the first to demonstrate that direct JNK inhibitors simultaneously inhibit metastatic outgrowth and promote events that may lead to tumour initiation within the native mammary gland. Whilst second-generation and emerging JNK inhibitors are designed with these pleiotropic functions in mind, they aim to overcome these complexities by preferentially targeting the JNK1 and JNK2 isoforms^37,38^. Although this approach is supported by a significant body of literature^39–41^, the effects of JNK1 and JNK2 are highly tissue- and tumour-specific^42–44^, and functional redundancy at this level of the pathway has been described^45,46^. On this basis, alternative strategies are needed to target this oncogenic signalling pathway in breast cancer patients.

Our study, driven by IHC analyses in clinical cohorts, demonstrates that the discrete functions of JNK in breast tissue are regulated by two spatially distinct subcellar pools, active under basal conditions; a nuclear tumour-suppressive JNK network involved in maintaining mammary gland architecture and genome stability, and a cytoplasmic oncogenic JNK network critical for the proliferation of disseminated TNBC cells at secondary sites. The existence of this cytoplasmic JNK pool aligns with published IHC analyses of JNK phosphorylation in TNBC patients^47^, while our assessment of its function was enabled by genetically encoded localisation-specific JNK inhibitors, originally developed for examining subcellular JNK signalling in neurons^20,21^. In the context of neurons, nuclear JNK activity is shown to be essential for apoptosis^48^, whilst constitutively active cytoplasmic JNK signalling regulates the architecture of neuronal processes through its effects on the microtubule and actin cytoskeletons^20–22,49^. These effects of cytoplasmic JNK activity on cytoskeletal organisation and cellular architecture parallel our observations in MDA-MB-231 TNBC cells, with ∼25% of the phosphosites deregulated by cytoplasmic JNK inhibition implicated in cytoskeletal regulation. Utilising unbiased phosphoproteomic and RNA-seq analyses, our results demonstrate that the subcellular targets of cytoplasmic JNK signalling extend well beyond the cytoskeleton, suggesting that multiple pathways ultimately contribute to the metastatic phenotype observed *in vivo*.

Nonetheless, the profound effect of cytoplasmic JNK inhibition upon cytoskeletal organisation and cell morphology allowed us to harness an actin-based phenotypic screen of 114,400 structurally diverse compounds^23^ and identify K12, a selective inhibitor of oncogenic JNK signalling. Phenotypic screening is an extremely productive drug discovery strategy, with retrospective analyses demonstrating that it yielded the majority of FDA-approved first-in-class drugs between 1999 and 2008^50^. Unlike target-based drug discovery, phenotypic screening allows for the unbiased identification of novel compounds based on a desired biological outcome and facilitates the discovery of drugs that elicit their effects through multiple protein targets^51^. Our data support that it is the poly-pharmacology of K12, leading to dual targeting effects on both GLS and the PDHC, which improves upon the modest effect seen with GLS knockdown *in vivo*. Importantly, these drug characteristics were unlikely to have been identified through a purely target-driven approach. Furthermore, the identification of a compound targeting metabolic enzymes through a cytoskeletal driven phenotypic screen, underscores the intertwined nature of these cellular pathways and the power of phenotypic screening to drive discoveries in a potentially unexpected manner.

Although there is a well-documented role of JNK in metabolic regulation^52^, and a strong literature basis linking both PDHC regulation and GLS to JNK signalling, our findings are distinct from published works. For instance, experiments on rat-derived neurons have found that stress-induced JNK activation inhibits oxidative metabolism through the direct phosphorylation of PDH^53,54^, whereas we observed indirectly increased PDH phosphorylation in response to the inhibition of basal cytoplasmic JNK activity. Given that PDH Ser232 is predominantly phosphorylated by pyruvate dehydrogenase kinase 1 (PDK1)^33,55^, our data suggest that cytoplasmic JNK modulates PDHC activity through a yet uncharacterised PDK1-dependent mechanism. Regarding GLS, the literature-based connection to JNK is strongly focused on the transcriptional regulation of GLS through c-Jun^56^. This is supported by pharmacological JNK inhibition studies where SP600125 treatment has been shown to decrease GLS expression at both the mRNA and protein levels, and by this reduce cellular GLS activity^57^. To our knowledge, our study is the first to demonstrate the converse effect, whereby GLS inhibition blocks JNK activity. Whilst this precise mechanism remains to be elucidated, our results suggest that this indirect effect of K12 on JNK activity likely sustains blockade of these essential energy-generating metabolic pathways through both post-translational and transcriptional mechanisms.

GLS has been extensively investigated as a therapeutic target in TNBC due to its high expression in these tumours and their well-reported dependence on glutamine metabolism^25^. Interest has been further driven by the development of CB-839, a potent, orally bioavailable inhibitor of GLS that has progressed to Phase I/II clinical trials for the treatment of various tumours, including TNBC^58,59^. Although our metabolomics analyses suggested that CB-839 is more effective than K12 at inhibiting glutamine hydrolysis, Seahorse target engagement assays showed that equimolar doses of K12 were just as effective at inhibiting glutamine-dependent oxygen consumption as CB-839. One explanation for this is compensatory glutamate dehydrogenase reactions that may occur in the K12 treated condition, which would increase glutamate levels and mask the inhibitory effect of K12 on GLS. Further studies are required to resolve this and to decipher the mechanism by which K12 treatment drives reductive carboxylation at high doses. It is intriguing to speculate that this may be due to the direct targeting of proteins within the α-ketoglutarate dehydrogenase complex, particularly as this complex shares protein components and regulatory mechanisms with the PDHC. Regardless, our results indicate that PDHC targeting by K12 ensures that K12 retains its anti-proliferative effects in the presence of pyruvate supplementation. This is particularly pertinent given that pyruvate anaplerosis is a key mechanism of resistance for CB-839^34^, rendering the poly-pharmacology of K12 a distinguishing feature that enhances its *in vivo* efficacy.

In summary, this study has demonstrated that cytoplasmic JNK signalling is responsible for the oncogenic functions of JNK and is critical for driving TNBC metastatic outgrowth. Further, we show that whilst direct JNK inhibitors are not suitable for use in this disease context, alternative strategies can be identified to target discrete downstream functions of this oncogenic signalling pathway. We envision that a therapeutic agent targeting the oncogenic functions of JNK in TNBC, such as K12, could be used in a similar manner to the highly successful adjuvant use of endocrine therapy and CDK4/6 inhibitors in ER+ breast cancer, which prevent relapse by inhibiting the growth of disseminated metastatic cells. While further drug development would be required to achieve long-term safety profiles for a prophylactic therapeutic approach aimed at preventing metastatic relapse, these proof-of-principle data with K12 open up new avenues for therapeutically targeting early-stage metastatic disease in TNBC patients.

## Acknowledgements

This work was supported by the National Health and Medical Research Council of Australia (APP1148099 to D.R.C and T.R.C; APP2011728 to D.R.C and S.L.L), Cancer Institute NSW (13/FRL/1-02 to D.R.C), National Breast Cancer Foundation (IIRS-18-121 to D.R.C and T.R.C; IIRS-20-032 to S.L.L, D.R.C and J.C.M), (H2020-MSCA-COFUND-2019 No. 945425 “DevelopMed” to R.S.) and Research Ireland (Frontiers for the Future Program grant 22/FFP-A/10729 to W.K.) funding.

## Author contributions

Conceptualization: S.L.L., D.R.C.

Methodology: S.L.L., O.G-C., N.S.B, L.Q., S.O’T., S.O., E.H., J.L., M.N., P.T., B.L.P., J.H., P.W.G., T.R.C., J.C.M., D.R.C.

Investigation: S.L.L. Y.E.I.O. O.G-C., M.S.C., N.S.B., E.T.Y.M., S.A.L., J.N., S.A., K.H.L., L.Q.,

K.J.M., C.P., I.D., M.M.N., C.T., A.C., M.P., R.S., E.M., J.Z.R.H., J.F.H., M.K., T.B., A.Y., S.O’T., S.R.O., J.L., M.H.T., M.N., L.G., B.L.P., D.R.C.

Visualization: S.L.L., Y.E.I.O., C.P., D.R.C.

Funding Acquisition: S.L.L., J.C.M., T.R.C., D.R.C.

Project Administration: S.L.L., J.C.M., D.R.C.

Supervision: S.L.L., E.H., W.K., P.T., J.H., P.W.G., T.R.C., J.C.M., D.R.C.

Writing – original draft: S.L.L., Y.E.I.O., D.R.C.

Writing – review and editing: all authors.

## Data and materials availability

AP-MS (PXD061936) and phospho-proteomic data (PXD063010) are available in ProteomeXchange via the PRIDE database. RNA-Seq data will be made available in the Gene Expression Omnibus (GEO) database prior to publication. Metabolomics data is available through GitHub (https://github.com/lakeeeq/20250405_CroucherTCATracingData). Requests regarding compounds identified in the original actin-based phenotypic screen by Bryce et al. should be directed to lead contact of that article (J.G.L). All other data are available from the lead contact (D.R.C) upon reasonable request.

## Materials and Methods

### Patient cohort analysis

JNK activity was assessed in published pan-breast cancer (457 patients)^18^ and TNBC (161 patients) TMAs^19^ provided by S.O’T. (Garvan Institute). IHC of paraffin-embedded sections was performed following heat-induced epitope retrieval using the rabbit polyclonal ppJNK (T183/Y185) antibody (Cat# AF1205, R&D Systems), as described in detail below. Samples were scored by a breast pathologist (E.M.), who calculated H-scores based on the percentage of tumour cells in the samples expressing nuclear or cytoplasmic ppJNK, and the intensity of the IHC signal. For both cohorts, an H-score cut-off of 100 was identified to distinguish between low and high nuclear ppJNK, whilst an H-score cut-off of 20 separated low and high cytoplasmic ppJNK. Molecular subtyping of the TNBC cohort was from van Geldermalsen *et al.*^26^.

### Immunohistochemistry (IHC)

All IHC sample preparation and staining was performed at the Garvan Institute Immunohistochemistry facility. IHC was performed on formalin-fixed paraffin-embedded sections using the Leica BOND RX (Leica, Germany). Briefly, samples were dewaxed and rehydrated in an ascending ethanol series (70-100%), prior to heat-induced epitope retrieval at 100°C for 20 min with a citrate (pH 6.0) epitope retrieval solution (Cat# AR9961; Leica). Primary antibodies: ppJNK (T183/Y185; Cat# AF1205; R&D Systems, 1:150-:300), Vimentin (V9; Cat# 14-9897-82; Invitrogen; 1:1000), Pan-Cytokeratin (C11; Cat# 4545; Cell Signaling Technology: 1:300), Phospho-Histone H2A.X (γH2AX, Ser139; Cat# 9718; Cell Signaling Technology; 1:500) or Phospho-Histone H3 (Ser10; Cat# 9701, Cell Signaling Technology; 1:100), were diluted in Leica antibody diluent and incubated for 30-60 min. Antibody staining was completed with the Leica Refine and DAKO Envision reagents. Samples were counterstained on the Leica Autostainer XL and coverslips applied with the Leica CV5030 glass coverslipper. Bright-field images were taken with the Aperio CS2 Slide Scanner (Leica) and NanoZoomer S210 (Hamamatsu Photonics, Japan). All quantification was performed using QuPath software (v 0.4.3).

### Tissue culture and stable cell line generation

Human MDA-MB-231, Hs578t, HCC70, MDA-MB-175 and MCF7 cell lines were purchased from ATCC. 4T1 cells and MDA-MB-231 tdTomato luciferase cells were provided by T.R.C. The MDA-MB-231, Hs578t, MDA-MB-175 and MCF7 lines were maintained in RPMI-1640 media supplemented with 10% fetal bovine serum (FBS), 1X penicillin-streptomycin (P/S), 20 mM HEPES and 140 IU insulin. HCC70 were cultured in 10% FBS, 1X P/S, 20 mM HEPES, 1 mM Na-Pyruvate and 9 g/L D-glucose. Telomerase-immortalized fibroblasts (TIFs-provided by P.T.), 4T1 and HEK293t cells were grown in Dulbecco’s Modified Eagle Medium (MEM) with 10% FBS and 1X P/S. All cell lines were maintained under standard culture conditions (5% CO_2_, 37°C). All human cell lines were validated with STR-profiling at the Garvan Molecular Genetics facility.

The pSH570MK Tet-On retroviral plasmid was provided by Prof. Tilman Brummer, Martin Koehler and Sebastian Herzog and contains the elements for tetracycline-inducible gene expression described previously^60^. To interrogate the discrete subcellular JNK pools, GFP-JBD (control), GFP-NES-JBD (C-JNKi) and GFP-NLS-JBD constructs provided by Prof. Eleanor Coffey were subcloned into pSH570MK using NotI and PacI restriction sites. For the production of stable cell lines, Plat-E cells were transfected with these constructs using jetPRIME transfection reagent (Polyplus), according to the manufacturer’s instructions. Viral supernatants were collected after 48 h, filtered and combined with 8 μg/mL polybrene. MDA-MB-231 stably expressing the ecotropic receptor were transduced with viral supernatant. Positive cells were selected with puromycin (1 μg/mL) and validated by Western blotting.

For the generation of stable MDA-MB-231 JNK-KTR-Clover cells, the pLentiPGK Puro DEST JNK-KTR-Clover was kindly provided by Markus Covert (Addgene #59151). For inducible GLS knockdown lines, stable cell lines were generated using SMARTvector Inducible Lentiviral shRNA (Horizon Discovery; GLS-A: GACTAAGATTCAACAAACT, GLS-B: CAGTCTGGAGGAAAGG TTG). In both cases, HEK293t cells were transfected with lentiviral vectors alongside pMDLg/pRRE (Addgene #12251), pRSV-Rev (Addgene #12253) and pMD.G (Addgene #12259) third generation packaging plasmids^61^ using Lipofectamine 2000 DNA transfection reagent (Thermo Fisher Scientific), according to the manufacturer’s instructions. Viral supernatants were collected after 48 h, filtered and combined with 8 μg/mL polybrene. MDA-MB-231 were transduced with packaged lentivirus and selected with puromycin (1 μg/mL). MDA-MB-231 JNK-KTR-Clover cells were further subjected to cell sorting using the FACS Aria^TM^ III Cell Sorter (BD Biosciences).

For FLIM-FRET imaging of JNK activity, the pCAGG-JNKAR1EV-NES was kindly provided by Dr Kazuhiro Aoki^62^, utilising source material from Prof. Jun-ichi Miyazaki^63^. For the production of stable cell lines, MDA-MB-231 cells were transfected with these constructs using jetPRIME transfection reagent, according to the manufacturer’s instructions. G418 selection pressure (2 mg/mL) was applied for 10 days post transfection and pure populations were isolated with cell sorting using the FACS Aria^TM^ III Cell Sorter.

### Preparation of collagen I from rat tails

Collagen I was extracted from rat tails, as previously described^64^. Briefly, manually extracted collagen tendons were solubilised in 0.5 M acetic acid for ≥48 hours at 4°C. Collagen was precipitated with the addition of 10% (*w/v*) NaCl, filtered through cloth wipers and pelleted by centrifugation at 10,000 RPM for 30 min at 4°C. Collagen precipitate was re-dissolved in 0.25 M acetic acid overnight at 4°C, dialysed in 17.4 mM acetic acid over 3-5 days and collagen I was collected by centrifugation at 14,000 RPM for 1.5 h at 4°C. Collagen concentration was measured using the Sircol^TM^ Soluble Collagen Assay Kit (Biocolor) and adjusted using 17.4 mM acetic acid to 2.5 mg/mL. All 2D experiments were performed on tissue culture plastic or glassware coated with 0.08 mg/mL collagen.

### Organotypic invasion assay

Organotypic matrices were generated as previously described^64^. Briefly, TIFs were incorporated into a collagen solution consisting of 74% (*v/v*) 2.5 mg/mL acid-extracted collagen, 8.9% (*v/v*) 10X MEM, 8.9% (*v/v*) 0.22M NaOH and 8.9% (*v/v*) FBS at a density of ∼1×10^5^ cells/mL, plated in 6-well plates and polymerised at 37°C. TIF-containing matrices were allowed to contract in culture for 12 days before being seeded under 2×10^5^ MDA-MB-231 cells in 24-well plates. After 3 days, matrices were placed on top of sterile mesh grid within media +/- doxycycline (2 μg/mL), creating an air-liquid interface to promote invasion. Matrices were collected after 14 days, fixed in 10% (*v/v*) neutral buffered formalin and processed for IHC analysis, as described above. Tumour cells were distinguished from TIFs using pan-cytokeratin staining and invasive distance was quantified in 2 cross sections of 3 replicate matrices (per experiment) using QuPath software (v 0.4.3).

### 3D-proliferation assay

For 3D-proliferation, TNBC cell lines were incorporated into a collagen solution consisting of 74% (*v/v*) 2.5 mg/mL acid-extracted collagen, 8.9% (*v/v*) 10X MEM, 8.9% (*v/v*) 0.22M NaOH and 8.9% (*v/v*) FBS. For 3-day treatment experiments, MDA-MB-231 cells were seeded at a density of 2.5×10^3^ cells/well. For 7-day treatment experiments, MDA-MB-231 and 4T1 were seeded at a density of 1.25×10^3^ cell/well, whilst Hs578t, HCC70 and MCF7 cells were seeded at 7.5×10^3^ cell/well. Cell-collagen mix (100 μL) was plated in 96-well plates and set at 37°C for 20 min before the addition of equal parts complete culture media. In the case of subcellular JNK inhibitor or drug treatment experiments, doxycycline (2 μg/mL) and drug treatments commenced 1 day post seeding. For GLS knockdown studies, cells were treated with doxycycline 2 days prior to seeding in collagen. In the majority of cases, cell viability was measured using a CellTiter 96® AQeous MTS assay (Promega) at indicated time points, with absorbance measured at 490 nm using a FluoSTAR Omega plate reader (BMG Labtech). Values from collagen-only controls were subtracted at each time point and data were normalised to untreated controls. Following the identification of metabolic targets for K12, relative cell numbers of MDA-MB-231 tdTomato Luciferase cells were measured with the addition of luciferin (1:100) and an endpoint luminescence readout on the FluoSTAR Omega plate reader.

### Cell lysates and Western blotting

For 2D subcellular fractionation experiments, sub-confluent cell monolayers cultured in 10 cm petri dishes were washed twice in ice cold 1X PBS and lysed in 0.1% (*v/v*) NP-40/1X PBS containing Na-orthovanadate and protease inhibitors (Cat#P8340; Merck). Whole cell lysate (WCL) fractions were collected, samples were spun for 10 sec on a tabletop centrifuge and cytoplasmic fractions (supernatant) were separated from nuclear fractions (pellet). Nuclear fractions were washed and resuspended in lysis buffer, then WCL and nuclear fractions were sonicated and all samples were denatured at 95°C in 1X NuPAGE LDS sample buffer (Cat# NP0007; Thermo Fisher Scientific).

For 2D samples, cells cultured in 6-well plates were washed twice in ice cold 1X PBS and lysed in normal lysis buffer containing 50 mM Tris-HCl (pH 7.4), 150 mM NaCl, 1 mM EDTA, 1% Triton X-100, Na-orthovanadate and protease inhibitors. For 3D samples, 5×10^5^ cells were cultured in 1 mL collagen plugs with a final FBS concentration of 0.25% (*v/v*). At experimental endpoint, collagen plugs were washed in ice cold 1X PBS and snap frozen. Plugs were then thawed in ice-cold 1X RIPA buffer containing 50 mM Tris-HCl (pH 7.4), 150 mM NaCl, 1% (*v/v*) NP-40, 0.5% (*w/v*) Na-deoxycholate, 0.1% (*w/v*) SDS, Na-orthovanadate and protease inhibitors, and sonicated for 30 sec on ice. Lysates were collected after centrifugation at 10,000 g for 5 min. All lysates were denatured at 95°C in LDS sample buffer.

All SDS gel electrophoresis and Western blotting were performed with the NuPAGE SDS PAGE and NuPAGE 4-12% Bis-Tris Precast Gels (Cat# NP0323; Thermo Fisher Scientific) according to the manufacturer’s instructions. Primary antibodies used in this study include: mouse monoclonal anti-β-actin (AC-15; Cat# A1978; Sigma Aldrich), mouse monoclonal anti-GAPDH (14C10; Cat# 2118; Cell Signaling Technology), rabbit polyclonal anti-JNK (Cat# 9252; Cell Signaling Technology), rabbit polyclonal anti-phospho-JNK (T183/Y185; Cat# 9251; Cell Signaling Technology), rabbit monoclonal anti-GLS (E4T9Q; Cat# 49363; Cell Signalling Technology), rabbit monoclonal anti-PDH (C54G1; Cat# 3205; Cell Signalling Technology), rabbit polyclonal anti-phospho-PDH (Ser232; Cat# 15289; Cell Signalling Technology) and mouse monoclonal anti-Lamin A/C (ARC5001-08; Cat#A19524; AbClonal). Anti-mouse-HRP (Cat# 7076; Cell Signaling Technology) and anti-rabbit-HRP (Cat# 7074; Cell Signaling Technology) secondary antibodies were used alongside Western lightning ultra chemiluminescent substrate (Cat# NEL113001; PerkinElmer) for signal detection.

### Phosphoproteomics Mass Spectrometry

Phosphoproteomics was performed on MDA-MB-231 cells cultured in 3D collagen matrices. For these experiments, 5×10^5^ cells were cultured in 1 mL collagen plugs with a final FBS concentration of 0.5% (*v/v*). Cells were either treated with doxycycline (2 μg/mL) or K12 (300 nM) for 24 h. At experimental endpoint, collagen plugs were washed in ice-cold 1X PBS and snap frozen. When thawed, samples were kept on ice, incubated in lysis buffer containing 100 mM TEAB buffer (pH 8.5), 10% Na-deoxycholate and Na-orthovanadate, sonicated and lysates were collected after centrifugation at 10,000 xg for 5 min. Subsequent mass spectrometry sample preparation and analysis steps were undertaken by The Australian Proteome Analysis Facility (APAF).

Vacuum-dried supernatants were processed using S-Trap columns (Protifi, USA). Samples were solubilized in 10% SDS and 100_mM TEAB, reduced with 10_mM dithiothreitol (56_°C, 30_min), and alkylated using 25_mM iodoacetamide (RT, dark, 30_min). Acidification was performed by adding 12% phosphoric acid to a final concentration of ∼1.2%, followed by dilution with S-Trap binding buffer (90% methanol, 100_mM TEAB, pH 7.55). Samples were loaded onto S-Trap columns, washed, and digested on-column with trypsin (∼20_μg/sample) for 2_h at 47_°C. Peptides were eluted sequentially with TEAB, 0.2% formic acid, and 50% acetonitrile containing 0.2% formic acid. Eluates were dried and reconstituted in 200_mM HEPES (pH 8.8), and peptide concentrations were determined using the Pierce colorimetric peptide assay.

Equal peptide amounts from each sample were labelled using 10-plex Tandem Mass Tags (TMT, Thermo Scientific, USA) in two batches. TMT reagents were reconstituted in anhydrous acetonitrile and added to individual peptide samples, followed by incubation for 1_h at room temperature. Labelling was quenched with 5% hydroxylamine (15_min, RT). To assess labelling efficiency and normalize pooling, a ‘label check’ was performed by mixing small aliquots of each labelled sample and analyzing by LC-MS/MS. Based on normalization factors, labelled peptides were pooled at a 1:1 ratio and desalted using C18 SepPak columns.

Phosphopeptides were enriched using a metal-affinity-based phosphopeptide enrichment kit (Thermo Scientific, USA). Peptides were reconstituted in binding buffer, incubated with activated resin (30_min), washed, and eluted. The flow-through (global proteome) was retained for further analysis. Eluted phosphopeptides were immediately dried and stored. Enriched phosphopeptides were fractionated using SDB-RPS StageTips into 10 fractions using a stepwise gradient of acetonitrile. Dried fractions were reconstituted in 0.1% formic acid for LC-MS/MS analysis. Global proteome samples were fractionated using the Pierce High-pH Reversed-Phase Peptide Fractionation Kit (Thermo Scientific, USA), following manufacturer’s instructions, and reconstituted in 0.1% formic acid before LC-MS/MS.

Phosphoproteomics data were evaluated using MaxQuant (v1.6.10.43), with MS/MS data processed using the Andromeda search engine within the MaxQuant software against all *Homo sapien* protein FASTA sequences in the Uniprot database. Pairwise-relative abundance comparisons were performed between Ctrl vs Ctrl+Dox, C-JNKi vs C-JNKi+Dox, N-JNKi vs N-JNKi+Dox, with significantly deregulated phosphosites defined as those with a localisation probability > 0.75, log fold change ≥ 1.2 and p-value ≤ 0.05. Cross-comparison between the different analyses revealed phosphosites uniquely deregulated by each of the inhibitor constructs. To capture the effects of cytoplasmic JNK inhibition on the MDA-MB-231 phosphoproteome, KEGG pathway analyses were performed using the EnrichR webtool and gene ontology analyses were performed using the ShinyGo (v0.81) webtool. The phophoproteomics data have been deposited to the ProteomeXchange Consortium via the PRIDE^65^ partner repository with the dataset identifier PXD063010

### RNA-sequencing

MDA-MB-231 cells were grown in 3D collagen matrices, as described above. After 48 h of doxycycline (2 μg/mL) treatment, RNA was extracted after homogenisation by PowerLyzer tissue disruption with the RNeasy Tissue Mini Kit (Quagen). RNA concentration and integrity were assessed using the Qubit™ 3 Fluorometer (ThermoFisher Scientific) and the 4200 Tapestation (Agilient). Sequencing libraries were subsequently prepared using the KAPA RNA HyperPrep Kit with RiboErase (Roche) and sequencing was performed on a S4 300 cycle flow cell on the NovaSeq 6000 System (Illumina) by the Garvan Genomics Platform at the Garvan Institute of Medical Research. Sequence reads were aligned using STAR (v2.7.3a) to the GRCh38 assembly with the gene, transcript, and exon features of Ensembl (release 105) gene model. Expression was estimated using RSEM^66^, and differentially expressed genes were identified using edgeR package. The EnrichR webtool was used for KEGG pathways analyses.

### Confocal microscopy and cellular phenotyping

MDA-MB-231 cells were seeded in collagen-coated black 96-well plates at a density of 250 cell/well. For analysis of cells stably expressing the inducible, localisation specific JNK inhibitors, cells were treated for 24 h with 2 μg/mL doxycycline. For evaluation of compounds sampled from clusters 2, 4, 5, 9, 11 and 18 of the Bryce et al. study^23^, MDA-MB-231 cells were treated with 10 μM of each compound for 24 h. Cells were subsequently fixed in 4% PFA for 20 min, before undergoing permeabilisation with 0.1% (*v/v*) Triton-X in PBS. Cells were washed in PBS and incubated with Phalloidin-iFluor 647 (Abcam), as per the manufacturer’s instructions. After DAPI counterstaining, z-stack images were collected on the Leica SP8 confocal microscope using a 40X/1.30 Oil objective. Confocal microscopy was performed at the Garvan Institute Imaging facility.

For image analysis, maximum intensity projections were generated with ImageJ and analysed with CellProfiler 4.1.3 using an optimized pipeline for quantification of the following 2D morphological parameters: cell area, perimeter, min Feret diameter, max Feret diameter, convex area, compactness (perimeter^2^ /4*π*Area), elongation (min/max diameter) and form factor (4*π*Area/perimeter^2^). Data were manually verified, ensuring that inaccurately demarcated/segmented objects were excluded from further analyses. In the case of doxycycline treated experiments, only GFP-positive cells were analysed. Once the morphological features associated with cytoplasmic JNK inhibition were defined, blinded scoring was undertaken to assess phenotype frequency across treatment conditions. With reference images provided to guide phenotype identification, the number of “small cells with cortical actin enrichment” were counted in de-identified images and the percentage of phenotype positive cells was calculated per well (5 images/well, 4 wells/condition). Samples were unblinded once the quantification of all conditions was completed.

### High content imaging

The 256 compounds identified in clusters 2, 4 and 5 by Bryce *et al.*^23^ were obtained from the Walter and Eliza Hall Screening Lab WECC diversity library. For the initial drug screen that sought to identify inhibitors of cytoplasmic JNK activity, 2.5×10^3^ MDA-MB-231 JNK-KTR mClover cells were seeded in Cellstar μclear cell culture 96-well F-bottom microplates (Greiner Bio-One) in 100 μL of growth media and cultured for 24 h. Each of the 256 compounds was tested in triplicate at a concentration of 5 µM in a final volume of 150 μL. Each plate also contained triplicate wells of the following three controls: 1) baseline control, which had a matched final concentration of DMSO, 2) JNK inhibitor control, where cells were treated with 10 µM JNK-IN-8 for 30 min, and 3) JNK activator control, where cells were treated with 300 nM anisomycin for 30 min. Cells were counterstained with 1 µg/mL Hoechst 33342 for 30 min, fixed with 4% paraformaldehyde for 30 min, washed twice with PBS and imaged on the Cellomics Arrayscan VTI HCS Reader.

Imaging and parameter optimisation, cell gating and preliminary data analysis was conducted in the native HCS Studio 2.0 Cell Analysis Software (6.0.3.4020), as previously described^67^. Briefly, a nuclear mask was generated using the Hoechst counterstain, and a 6-pixel cytoplasmic ring was generated in all directions excluding neighbouring nuclei. These masks were used to determine the average intensity of the JNK-KTR biosensor within the nucleus and cytoplasm and in turn calculate the cytoplasmic-to-nuclear ratio of the biosensor, which is directly proportional to JNK kinase activity^24^. Compound screening was repeated at lower doses in instances where crystalline artefacts were observed in images. Six compounds were eliminated from the screen on the basis that they caused mis-localisation of the biosensor between the two compartments. Single cell data were collated from the triplicate wells and outliers were removed using the ROUT method (Q = 0.1%). Data were normalised to the baseline and JNK inhibitor controls from the respective plate and mean values were collected for each treated condition. Compounds were ranked in ascending order of their relative JNK activity and top hits were identified as those with a mean value below the arbitrary cut-off of 0.25.

For the secondary drug screen that sought to exclude inhibitors of stress-induced JNK activity, each of the top 41 hits was tested as described above, with the addition of 300 nM anisomycin to each well 30 mins prior to fixation. Samples were counterstained, fixed, imaged and analysed as described for the initial screen. Inhibitors of stress-induced JNK signalling were defined as any compounds that resembled JNK-IN-8 and reduced relative JNK activity below baseline levels.

For siRNA screens, 750 MDA-MB-231 JNK-KTR mClover cells were seeded in Cellstar μclear cell culture 96-well F-bottom microplates (Greiner Bio-One) in 150 μL of growth media and cultured for 24 h. A custom library of siGENOME SMART pool siRNA (Millenium Science) were diluted to 20 µM according to the manufacturer’s instructions. Lipofection was performed using jetPRIME transfection reagent according to the manufacturer’s instructions. Briefly, siRNA were diluted to 200 nM in 50 μL jetPRIME buffer prior to the addition of 1 μL jetPRIME transfection reagent. Samples were mixed and incubated for 10 min at RT before being added to plates at a final concentration of 15 nM and incubated for 72 h. Samples were counterstained, fixed, imaged and analysed as described for the drug screens above. Data were normalised to the cells treated with the negative control siRNA ± JNK-IN-8 (10 µM, 30 min) and potential targets were ranked in ascending order of their relative JNK activity, with top hits identified as those with a mean value below the arbitrary cut-off of 0.25.

### K12 Formulation Development, Maximum Tolerated Dose and Toxicokinetics Studies

Formulation, maximum tolerated dose (MTD) and exposure studies were performed by Aragen Life Sciences Private Limited (Hyderbad, India). K12 was dissolved directly in N-methyl-2-pyrrolidone (NMP) and stored at −80°C. K12 solutions were then prepared immediately prior to i.p. injection, with final formulations comprising 0.05% (*w/v*) compound in 1 part NMP (10% *w/v* final conc.), 4 parts Kolliphor HS15 (40% *w/v* final conc.) and 5 parts 1% Hydroxypropyl methylcellulose E3 prepared in phosphate buffer (pH 6.8; 0.5% *w/v* final conc.). Compounds were omitted from vehicle solutions. For MTD studies, vehicle control and escalating doses of K12 were evaluated in Swiss albino mice. Specifically, 3 male and 3 female mice received a single i.p. injection of 10 mL/kg vehicle or each dose formulation and were monitored for clinical signs and morbidity/mortality for up to 72 h, with daily body weights measured and blood collected at end-point. From this, a maximum tolerable dose of ≥3 mg/kg was determined for K12. For TK studies, 6 male mice received a single i.p. injection of K12 (3 doses tested up to the MTD). Blood was collected in Li-Heparin tubes at 0.25, 0.5, 1, 3, 7 and 24 h post-dose (3 animals/time-point), plasma was separated and stored at −80°C for exposure analysis of plasma concentration (ng/mL).

### Mouse models

Animal experiments were conducted in accordance with the Garvan/St Vincent’s Animal Ethics Committee guidelines (19/08, 21/15 and 24/15) and in compliance with the Australian code of practice for the care and use of animals for scientific purposes. Unless specified otherwise, mice were bred at the Australian BioResources (ABR) Pty Ltd breeding and holding facility and transferred to the Garvan Institute at 6-8 weeks of age. For each model, mice were housed in individual ventilated cages in a 12-hour light/dark cycle and fed *ad libitum*. Experiments were initiated after an acclimatization period of at least seven days, at which point mice were monitored by weight and tumor measurement (where applicable) three times per week.

### 4T1 orthotopic model

BalbC/AusJ mice received intra-mammary injection of 1.0×10^5^ 4T1 cells in 100µL of dPBS into the left-fourth mammary fat pad. Tumours were palpable (∼5 mm^3^) 7 days post-injection at which point mice were randomly allocated to treatment groups (10 mice/treatment arm). Two experiments were conducted using this model, the first comparing vehicle (3% DMSO in PBS) with CC-401 (25 mg/kg) treatment and the second comparing vehicle (10% NMP, 40% Kollophor HS15, 0.5% hydroxypropyl methylcellulose E3 in phosphate buffer) with K12 (3 mg/kg). In both cases, treatments were administered thrice weekly by intraperitoneal (i.p.) injection, rotating between quadrants. Body weight and tumour size were monitored twice weekly, and mice were sacrificed at experimental endpoint (4 weeks post injection), at which point tumour burden was determined (mouse weight/tumour weight). Primary tumours, mammary glands and lungs were collected for follow-up IHC analysis. In the case of the vehicle vs K12 experiment, kidney, spleen and liver were also collected for drug toxicity assessment by a clinical pathologist (S.O’T).

For mammary gland analysis, mammary gland whole mounts were generated by spreading glands on a Superfrost^TM^ glass slide prior to fixation in 10% neutral buffered formalin. Glands were defatted in acetone overnight prior to overnight staining in carmine alum solution (0.2% *v/v* carmine, 0.5% *v/v* aluminum sulfate). Whole mounts were dehydrated with a graded ethanol series, cleared with Slide Brite xylene substitute and stored in methyl-salicylate prior to imaging with a Leica MZ 12 Dissecting Microscope. ImageJ software was used to measure the number of branches per mm of duct, as well as the width of secondary ductal branches and terminal end buds. Once imaged, mammary glands were rehydrated in a graded ethanol series and processed for IHC alongside primary tumours and lung tissue. For the analysis of pulmonary metastases, H&E sections were manually annotated using QuPath software and both the number of metastases per lung area and metastatic burden (total met area/lung area) were calculated for each mouse.

### Intraductal xenograft model with inducible, localisation specific JNK inhibitors

For intra-ductal injections, immune-compromised NOD.Cg-Prkdc^scid^ IL2rg^tm1Wjl^/SzJ(Ausb) (NSG) female mice were anesthetized and 8×10^4^ MDA-MB-231 cells stably expressing the respective control or JNK inhibitor construct (20 mice/cell line; 3 cell lines) were directly injected into the nipple of the left, 4^th^ mammary gland. Body weight and tumour size were monitored twice weekly. Once tumours reached 100 mm^3^, mice were randomized and allocated control or DOX-containing chow. Variations in mouse numbers between groups are underpinned by an absence of tumour growth (all but one case) or premature ethical endpoint (single case). Mice were sacrificed at ethical endpoint, defined as when the primary tumour exceeded a tumour volume/body weight ratio of 10% (*v/w*), or upon signs of respiratory distress. Primary tumours and lungs were excised, formalin fixed and paraffin embedded for IHC analysis, as described above. In this case, pulmonary metastases were stained with a vimentin antibody and metastatic burden (total vimentin area/lung area) was annotated through automated cell detection analyses using QuPath software.

### MDA-MB-231 colonisation models

For K12 *in vivo* analyses, female 8-week-old immune-compromised NSG mice received an i.v. injection of 1.5×10^5^ MDA-MB-231 cells in PBS. Mice were randomly allocated to treatment groups (10/treatment; 4 treatments), with thrice weekly vehicle and K12 (0.75, 1.5 and 3 mg/kg) i.p. injections commencing one day post cell-injection. Body weight was monitored twice weekly, and mice were sacrificed at experimental endpoint (4 weeks post injection), at which point mammary glands, lungs, kidney, live and spleen were collected for IHC analysis, as described above. Pulmonary metastases were stained with a vimentin antibody and the number of metastases per lung area and metastatic burden (total met area/lung area) were calculated for each mouse. For drug toxicity assessment, kidney, liver and spleen samples were evaluated by a clinical pathologist (S.O’T).

For the *in vivo* analysis of control and GLS shRNA knockdown MDA-MB-231 cells, immune-compromised NSG mice received an i.v. injection of 1.5×10^5^ cells in PBS (3 cell lines; 20 mice per line). Mice were allocated control or DOX-containing chow and sacrificed at experimental endpoint (4 weeks post injection), at which point lung tissue was collected for IHC analysis, as described above. In this case, pulmonary metastases were stained with a vimentin antibody and metastatic burden (total vimentin area/lung area) was annotated through automated cell detection analyses using QuPath software.

### FLIM/FRET Multiphoton imaging

For *in vitro* on-target validation experiments, 1×10^5^ cells were seeded in 2 mL collagen plugs within a Cellvis glass bottom imaging dish (Cat# D35-20-1.5-N), cultured overnight in complete growth media and then treated with DMSO or 2.5 μM K12 for 24 h. For *ex vivo* on-target validation experiments, 2×10^5^ cells were i.v. injected into NOD.Cg-Prkdc^scid^ IL2rg^tm1Wjl^/SzJ(Ausb) mice and metastases were allowed to develop for 4 days prior to a single i.p. injection of vehicle or K12 (3 mg/kg). Lungs were harvested 48 h post treatment. In both cases, images were acquired with the Leica DMI 6000 SP8 inverted confocal microscope using a 25x 0.95 numerical aperture (NA) water immersion objective (HC FLUOTAR L 25x/0.95 W VISIR 0.17). The Ti:Sapphire femtosecond laser (Chameleon Ultra II, Coherent) excitation source operating at 80 MHz was tuned to a wavelength of 840 nm and signals were collected with the HyD-RLD detectors using 483/40 and 435/40 nm band pass emission filters for enhanced cyan fluorescent protein (ECFP) and collagen I second-harmonic generation (SHG), respectively. Alternatively, a Leica Stellaris 8 set up was utilized equipped with 25x 1.00 numerical aperture (NA) water immersion objective (HC IRAPO L 25x/1.00 W motCORR), 4 tunable HyD-RLD hybrid detectors and two multiphoton lasers, an InSight X3 and MaiTai DeepSee (SpectraPhysics). Imaging was performed using the 80 MHz InSight X3 laser at 840 nm and HyD-RLD detectors were tuned to 440-510 and 410-430 emission windows for cyan fluorescent protein (ECFP) and collagen I second-harmonic generation (SHG) detection respectively. Images were acquired at 512×512 pixels, a line rate of 700 Hz and a total of 203 frames per image. Motion correction was performed using Galene (v2.0.2)^68^ and single-cell analysis was performed using FLIMfit (v5.1.1)^68^, as described previously^69^. FLIM/FRET imaging was performed at the Garvan Institute Imaging facility and ACRF INCITe centre.

### K12 Synthesis, Bead Design and Generation

The synthesis of K12 is summarised in **Fig. S9A**. Commercially available 2-chloropropionitrile 1 and ethyl 2-amino-4,5-dimethylthiophene-3-carboxylate 2 were reacted in the presence of dry HCl gas to generate the pyrimidinone 3. Coupling of thione 4^70^ and pyrimidinone 3 in ethanol afforded K12 in 83% yield. See supplementary information for full synthetic procedures. Using the K12 synthesis, 6 analogues were prepared to investigate the structure-activity relationship of the scaffold (**Fig. S9B**). Changes to the C5 position had the least impact on activity in 3D growth assays (**Fig. S9C**). Therefore, it was decided to pursue this position for bead conjugation (**Fig. S9D**). K12 linker-compound 7 was synthesised with a terminal propargyl group. Following the procedure developed by Finn and co-workers^71^, this molecule was coupled to azide-functionalised agarose bead 8 in a “click” reaction to generate K12 affinity bead 9. This methodology was repeated using a negative control substrate 10 to generate negative control bead 11. See supplementary information for full synthetic procedures.

### Affinity Purification-Mass Spectrometry (AP-MS)

Sepharose beads with immobilised K12, and control beads, were incubated with lysates prepared from MDA-MB-231 breast cancer cells. This was performed in the presence and absence of free K12 (300 nM) to competitively remove specific binding partners. Bound proteins were digested off the beads in 2 M urea containing 1 mM tris(2-carboxyethyl)phosphine and 5 mM 2-chloroacetamide in 100 mM Tris pH 8.5 using 0.4 µg of trypsin overnight at 37°C while shaking at 1600 rpm. The peptides were purified using reversed-phase sulfonate polystyrene-divinylbenzene microcolumns. Peptides were separated on a Dionex 3500 nanoUHPLC, coupled to an Orbitrap Lumos mass spectrometer via electrospray ionization in positive mode with 1.9 kV at 275 °C and RF set to 30%. Separation is achieved on a 50 cm × 75 µm column packed with C18AQ (1.9 µm) over 40 min at a flow rate of 300 nL/min. Peptides were eluted over a linear gradient of 3–40% Buffer B (Buffer A: 0.1% v/v formic acid; Buffer B: 80% v/v acetonitrile, 0.1% v/v FA) and the column was maintained at 50°C. The instrument was operated in data-independent acquisition (DIA) mode, with an MS1 spectrum acquired over the mass range 350–1,400 m/z (60,000 resolution, 100% automatic gain control (AGC), and 45 ms maximum injection time) followed by sequential MS/MS spectra across 13.7 m/z isolation windows with 1 m/z overlap covering the full mass range. MS/MS data will be acquired with higher-energy collisional dissociation (HCD) fragmentation (15,000 resolution, 2000% AGC, 55 ms maximum injection time, and normalized collision energy 30%). Data were processed in Spectronaut v17.6.230428.55965 with default setting against the Homo sapien protein FASTA sequences in the Uniprot database and filtered to 1% FDR at the PSM, peptide and protein level. Statistical analysis was performed in Perseus^72^ with 2-way t-tests and correction for multiple hypothesis testing with Benjamini Hochberg and a q-value < 0.05 used as statistical cut-off.

### Molecular modelling

Protein structures were taken from the Research Collaboratory for Structural Bioinformatics Protein Data Bank archive (RCSB PDB): accession code 5HL1 for GLS (Crystal structure of glutaminase C in complex with inhibitor CB-839)^73^. All rigid docking calculations were performed using Glide with the OPLS3 forcefield provided within the Maestro interface of the Schrodinger software suite (version 13.8)^74–77^. The receptor structure was prepared and minimised within the Protein Preparation Wizard. In each case, the receptor grid used was a 10×10×10 Å^3^ inner box defined by the centre of the co-crystallised ligand of the protein used. Each docking run was carried out using the standard settings for Glide, with the following modifications; XP (extra precision) scoring function was used and the number of poses written per ligand was increased from 1 to 5. All ligands were drawn in Maestro using the 2D sketching tool then prepared using the Ligand Preparation module. This implements the OPLS3 forcefield and Epik to determine the low energy conformation and the likely ionisation state between pH 5-9^77–79^. All images of crystal structures or docking were generated using PyMOL Molecular Graphics System, Version 2.5.5 Schrödinger, LLC.

### Surface Plasmon Resonance (SPR)

SPR was performed on the Biacore T200 (Cytiva) using high-affinity streptavidin (SA) sensor chips. Biotinylated K12 (100 nM) was immobilised according to manufacturer’s instructions and a kinetic analysis was performed with recombinant GLS (Abcam, ab1980650) diluted in HBS-EP+ buffer. Regeneration steps were performed with 10 mM Glycine (pH 1.5).

### Seahorse Assays

Oxygen consumption rate (OCR) and extracellular acidification rate (ECAR) were measured by the Seahorse XFe24 Analyzer. MDA-MB-231 cells (80,000 in 100 μL/well) were seeded in a Seahorse XF24 cell culture microplate, allowed to settle at room temperature for 1 h. Following attachment, 150 µL normal growth media was added and cells were incubated overnight at 37°C. The Seahorse XFe24 sensor cartridge was hydrated with 1 mL sterile Milli-Q water/well, with all sensors fully submerged, and then placed in a 37°C non-CO_2_ humidified incubator. Milli-Q water was replaced with 1 mL warm XF calibrant solution 1 h prior to the assay and incubated in a non-CO_2_ incubator at 37°C. Growth media in the cell microplate was replaced by a final volume of 500 μL XF RPMI media and the cells were incubated in a non-CO_2_ incubator at 37°C for 1 hour before the assay. Treatment for each injection portal was prepared to the final concentration and loaded to the corresponding portal (50 μL each) before loading onto the analyser for the assay. The measurement of OCR and ECAR commenced under basal conditions and after sequential addition of the following: XF RPMI media (added after 15 min), CB-839 or K12 (0 or 30 or 300 nM; added after 30 mins), glutamine (2 mM, added after 1 h) and XF RPMI media (added after 30 min). Upon completion of the assay run, cells from each well were collected by RIPA buffer and the protein content was quantified by an BCA assay for result normalisation.

### Metabolite extraction and targeted metabolomics for glutamine and glucose tracing

MDA-MB-231 cells were seeded at 1 × 10^6^ cells per well in a 6-well plate (Thermo) in complete RPMI media and allowed to adhere overnight. DMSO (vehicle control; 0.2%), K12 (10, 30 or 300 nM) or CB-839 (10, 30, or 300 nM) were added to cells for a 2 h incubation. Media was removed and cells were washed with PBS prior to incubation with the same vehicle control, K12 or CB-839 concentrations in glutamine/glucose free RPMI containing 10% dialysed FBS (Gibco, #30067-334) supplemented with either 2 mM ^13^C_5_-glutamine (Sigma) or 11.1 mM ^13^C_6_-glucose (Sigma) for 6 hours. After 6 h, media was removed, cells were gently rinsed twice with cold PBS and the plates were immediately stored in the freezer at −80 °C until metabolite extraction.

Metabolites were extracted using a modified Folch method (chloroform:methanol:water at 2:1:1 *v/v* ratio)^25^. Cells were lysed by rinsing and scraping the wells twice with 300 μL methanol:water (1:1 *v/v*) containing internal standards, with the cell slurry combined in a microfuge tube. After 600 μL chloroform was added, the cell slurry was vortexed and then centrifuged (15000 g, 4°C, 10 min) to induce phase separation. The top aqueous layer was transferred into fresh microfuge tube and dried completely in a SpeedVac (Savant, Thermo Scientific) without heat over 3 h. Dried extracts were stored in freezer awaiting LC-MS analysis. Extracts were resuspended by vortex in 60 μL buffer (2.5% acetonitrile, 10 mM ammonium acetate and 10 mM ammonium hydroxide). After centrifuging, 50 μL supernatant was transferred into HPLC vials. Targeted metabolomics for glucose tracing^80^ and glutamine tracing^81^ was carried out as described previously using LC-MS/MS.

For central carbon metabolites (glycolysis and TCA cycle), samples were assayed on a Sciex QTRAP 6500+ MS coupled to Agilent 1260 Infinity LC. Reverse phase ion pairing LC separation was achieved using Synergi™ 2.5 µm Hydro-RP (Phenomenex, 100 Å, LC Column 100 x 2 mm (00D-4387-B0)^82^, using mobile phase Buffer A 3% acetonitrile, 15 mM acetic acid and 10 mM tributylamine (ion pairing), and Buffer B 100% acetonitrile. For amino acids, glutamine-tracing samples were assayed on a Sciex QTRAP 6500+ MS coupled to Shimadzu Nexera LC-40 UHPLC. HILIC-based LC separation was achieved using an Poroshell 120 HILIC-Z (Agilent, 2.1×100 mm, 2.7uM), using mobile phase Buffer A 5% acetonitrile, 20mM ammonium hydroxide and 20 mM ammonium acetate, and Buffer B 100% acetonitrile. Chromatographic alignment, peak identification and integration were carried out using MSConvert (Proteowizard) and in-house MATLAB scripts^83^. The total peak area of metabolites detected by both LC-MS platforms was normalised to the protein concentration (Lowry assay) of cell pellets collected post metabolite extraction.

### Statistical Analyses

Data were analysed and represented with GraphPad Prism 10 software, unless specified elsewhere. For patient data, p-value scanning was undertaken with the Gehen-Breslow-Wilcoxon test to determine H-score cut-offs for low vs high nuclear and cytoplasmic JNK activity. Survival data were evaluated with the Kaplan-Meier method and log-rank test, with data displaying the Mantel-Haenszel hazard ratio. For *in vivo* drug analyses, ROUTS outlier analysis (Q = 1%) was performed on metastatic burden data, with any outlier’s identified removed from all other analyses. For *in vitro* and *in vivo* datasets, data distribution was determined with the D’Agostino & Pearson test or Kolmogorov-Smirnov test (in the case of low n), which determined the appropriate test for group comparisons. Where two conditions were being compared, parametric t-tests were performed if data followed a gaussian distribution or non-parametric Mann-Whitney tests were performed if data were not normally distributed. Where more than two conditions were being compared and all datasets fit a gaussian distribution, one-way ANOVAs were performed with either a Dunnett’s or Sidak’s post-test. For data that did not fit a normal distribution, the Kruskal-Wallis test was performed with a Dunn’s post-test. For datasets with multiple comparisons, two-way ANOVA analysis was performed using the Geisser-Greenhouse correction followed by a Tukey or Sidak’s multiple comparisons post-test. In all cases: ns = not significant, * p<0.05, ** p<0.01, *** p<0.001 and **** p<0.0001.

## Supplementary Information

### Supplementary Methods

– Chemical Synthesis

### Supplementary Figure Legends

**Supplementary Figure 1. Prognostic significance of subcellular JNK activity in breast cancer (related to Fig. 1). (A)** Nuclear vs cytoplasmic ppJNK H-Scores for samples analysed in the pan-breast cancer cohort, excluding those belonging to the TNBC subtype. Kaplan-Meier analysis (log-rank test, Mantel-Haenszel hazard ratio) of **(B)** overall survival and **(C)** relapse-free survival in the pan-breast cancer cohort. For both analyses, low nuclear JNK ≤ 110, high nuclear ppJNK >110, low cytoplasmic ppJNK ≤ 20, high cytoplasmic ppJNK >20. In all cases: * p<0.05, ** p<0.01, *** p<0.001 and *** p<0.0001.

**Supplementary Figure 2. Effect of 2D vs 3D culture on subcellular JNK activity in breast cancer cell lines (related to Fig. 3). (A)** Subcellular fractionation of cells cultured in 2D including whole cell lysates (WCL), cytoplasmic and nuclear fractions were collected from ER+ MCF and MDA-MB-175 cell lines, and TNBC HCC70, 4T1, Hs578t and MDA-MB-231 cell lines. Western blot analyses show Lamin A/C as a marker of the nuclear compartment and GAPDH as a marker of the cytoplasmic compartment, along with total JNK and ppJNK (T183/Y185). (B) IHC analysis of ppJNK (T183/Y185) in MC7, HCC70, 4T1, Hs578t and MDA-MB-231 cell lines cultured in 3D collagen matrices. **(C)** Representative Western blots of MCF7 cell lines stably expressing the GFP-fused control (Ctrl), cytoplasmic JNK inhibitor (C-JNKi) and nuclear JNK inhibitor (N-JNKi) constructs ± doxycycline. **(D)** Growth curves of these cell lines cultured in 3D collagen matrices ± doxycycline. Data are from a single representative experiment with 5-6 technical replicates per condition, expressed relative to the Ctrl without doxycycline treatment condition at day 7 (mean ± SD; mixed effects analysis with Tukey post-test).

**Supplementary Figure 3. Phoshoproteomics analysis defines the downstream targets of cytoplasmic JNK (related to Fig. 4). (A)** Manual curation of the subcellular localisation of each phosphoprotein significantly deregulated by expression of the C-JNKi construct (C-JNKi + DOX). Proteins are first classed as cytoplasmic (CYT), shuttling (CYT/NUC), nuclear (NUC) or secreted (SEC). Cytoplasmic proteins with a known localisation pattern (not simply cytosolic) were then subcategorized based on their distribution at the plasma membrane (PM), cytoskeleton (CSK), endoplasmic reticulum (ER), mitochondria (MT), golgi (GOL), intracellular vesicles (VES), junctions (JNC), centrosome (CTS), focal adhesion (FA), endosomes (END), stress granules (SG), autophagosomes (AUT) or P-bodies (PBD). Nuclear proteins with a known localisation pattern were then subcategorised based on their distribution at the nucleolus (NCL), nuclear membrane (NME), nuclear pore complex (NPC), nuclear speckles (NSP), nuclear bodies (NB) or Cajal bodies (CB). **(B)** KEGG pathway analysis of phosphoproteins significantly deregulated by expression of the C-JNKi (C-JNKi + DOX; fold change > 1.2, p-value < 0.05). **(C)** Gene ontology analysis of the phosphosites significantly down-regulated (left) and up-regulated (right) by expression of the C-JNKi construct (C-JNKi + DOX).

**Supplementary Figure 4. Analysis of F-actin phenotypes for actin-based phenotypic screening (related to Fig. 5). (A)** Blinded scoring of the percentage of cells with a “C-JNKi phenotype”, defined by small cells with cortical actin enrichment, performed in MDA-MA-231 cells stably expressing the Ctrl, C-JNKi and N-JNKi constructs ± doxycycline (mean ± SD; one-way ANOVA with Sidaks post-test). **(B)** Phalloidin staining of F-actin structures in MDA-MB-231 cells treated with compounds sampled from Bryce *et al.* phenotypic clusters 2, 4, 5, 9, 18 and 21 (10 µM, 24 h). Scale bars represent 30 µm. **(C)** Blinded scoring of the percentage of cells with a “C-JNKi phenotype”, defined by small cells with cortical actin enrichment, performed in MDA-MA-231 cells treated with compounds sampled from Bryce *et al.* phenotypic clusters 2, 4, 5, 9, 18 and 21 (10 µM, 24 h). Data show mean ± SD from 4 technical replicates (Kruskal-Wallis with Dunns post-test). **(D)** (i) Quantitative analysis of morphological parameters assessed with CellProfiler 4.1.3. Heatmaps show the mean fold change for each parameter relative to the DMSO control. Data are calculated from 20 images from 4 technical replicates from a single experiment. Numbers on the right indicate the overlap between each compound and the C-JNKi + doxycycline condition shown in Fig. 5B. ii) Blinded scoring of phenotype positive cells (small cells with cortical actin enrichment) in 4 technical replicates from a single experiment. In all cases: ns = not significant, * p<0.05, ** p<0.01, *** p<0.001 and *** p<0.0001.

**Supplementary Figure 5. Screen developed to identify selective inhibitors of cytoplasmic JNK signalling (related to Fig. 5). (A)** Representative high-content images of MDA-MB-231 cells expressing the JNK-KTR mClover biosensor following treatment with DMSO, JNK-IN-8 (10 µM, 2 h) and anisomycin (300 nM, 2 h). **(B)** High content analysis of single cell JNK activity in MDA-MB-231 cells expressing the JNK-KTR mClover biosensor following treatment with DMSO, JNK-IN-8 (10 µM, 2 h) and anisomycin (300 nM, 30 min). **(C)** High content imaging of MDA-MB-231 JNK-KTR mClover cells treated for 24 h with 2.5-5 µM of the 11 commercially available top hit compounds. Relative JNK activity (JNK-KTR cytoplasmic/nuclear ratio) was calculated from single cell data obtained from 3 technical replicates, normalized to DMSO and JNK-IN-8 controls (mean ± 95% CI). **(D)** High content imaging of MDA-MB-231 JNK-KTR mClover cells treated for 24 h with 2.5-5 µM of the 11 commercially available top hit compounds, followed by anisomycin treatment (300 nM, 30 min). Relative JNK activity (JNK-KTR cytoplasmic/nuclear ratio) was calculated from single cell data obtained from 3 technical replicates, normalized to DMSO and JNK-IN-8 controls (mean ± 95% CI). **(E)** Heatmap showing the mean cell viability obtained from MDA-MB-231 cells treated with varying doses of the 11 top hit compounds (0.156-10 µM, 72 h). Data are from a representative experiment performed in triplicate and normalized to the untreated control. **(F)** MDA-MB-231 cells cultured in 3D collagen matrices were treated for 7 days with increasing doses of K12 doses. Data shown are from a representative experiment where cells were seeded on day 0, treated with K12 on day 1 and cell viability was measured every 48 h from 2 days post treatment (mean ± SD). **(G)** Schematic representation of JNK-AR1-EV NES biosensor used for FLIM/FRET imaging of single cell cytoplasmic JNK activity. **(H)** Densitometry analysis of ppJNK/total JNK from Western blots shown in Fig. 5K. Values are normalised to actin loading controls and then to the DMSO control (mean ± SD, n=3). **(I)** Quantitative analysis of morphological parameters assessed with CellProfiler 4.1.3. Heatmaps show the mean fold change for each parameter relative to the DMSO control. **(J)** Blinded scoring of the percentage of cells with a “C-JNKi phenotype”, defined by small cells with cortical actin enrichment, in MDA-MA-231 cells treated with K12 (2.5 µM, 24 h).

**Supplementary Figure 6. Extended evaluation of K12 in NSG and BALB/c murine models (related to Fig. 6). (A)** Assessment of NSG mouse weight throughout the course of the MDA-MB-231 colonisation model used to evaluate K12 (0.75, 1.5 and 3 mg/kg) efficacy following 4 weeks of thrice weekly i.p. injections (mean ± SD; mixed effects analysis with Tukey post-test). **(B)** Representative H&E-stained kidney sections from mice at experimental endpoint. **(C)** Representative H&E-stained spleen sections f. **(D)** Representative H&E-stained liver sections. **(E)** Analysis of the size of MDA-MB-231 pulmonary metastases in vehicle vs K12 (0.75, 1.5 and 3 mg/kg) treated NSG mice after 4 weeks of thrice weekly i.p. treatment (Kruskal-Wallis with Dunns post-test). **(F)** Assessment of BALB/c mouse weight throughout the course of the 4T1 colonisation model used to evaluate K12 (3 mg/kg) efficacy following 4 weeks of thrice weekly i.p. injections (mean ± SD; mixed effects analysis with Sidaks post-test). **(G)** Analysis of end-point tumour burden (mean ± SD; Mann-Whitney test). In all cases ns = not significant.

**Supplementary Figure 7. Kinome profiling of K12 (related to Fig. 7).** Eurofins KINOMEscan assay assessing the effect of 2.5 µM K12 across 468 human kinases.

**Supplementary Figure 8. Synthesis of K12, K12 structural analogues and K12 affinity beads (related to Fig. 7). (A)** Scheme depicting K12 synthesis. **(B)** Six K12 analogues were synthesized to investigate the structure activity relationship of the scaffold and identify sites for bead conjugation. **(C)** Assessment of MDA-MB-231 cell viability following treatment with the original batch of K12 (*), newly synthesized K12 (^#^) or the six K12 analogues (K12-1 through to K12-6). Heatmap shows the mean change (n=6) in cell viability relative to untreated DMSO controls. **(D)** Synthesis of K12 affinity beads (top) and negative control beads (bottom) used in AP-MS experiments.

**Supplementary Figure 9. Molecular docking of K12 enantiomers into the allosteric binding pocket of the GLS homotetrameric structure (PDB: 5HL1; related to Fig. 7). (A)** GLS (blue/grey) with K12-S (green) docked into the allosteric binding pocket. **(B)** K12-R (cyan) docked into the allosteric binding pocket of the GLS homotetramer (blue/grey). **(C)** CB-839 (magenta) bound to the allosteric binding pocket of the glutaminase-1 homotetramer (blue/grey).

**Supplementary Figure 10. Evaluation of K12 on glutamine-and glucose-dependent metabolic pathways (related to Fig. 8). (A-E)** Metabolomics was used to trace U-^13^C_5_-glutamine for 6 h in MDA-MB-231 cells following treatment with DMSO control, CB-839 (300 nM) or K12 (10, 30 or 300 nM). Unlabelled metabolite (^12^C) is denoted by m+0, whilst m+x denotes a metabolite with x number of carbons labelled with ^13^C. **(A)** Total peak area of α-KG. **(B)** Total peak area of succinate. **(C)** Total peak area of Fumarate. **(D)** Total peak area of Malate. **(E)** Total peak area of Citrate. **(F-G)** Metabolomics was used to trace U-^13^C_6_-glucose for 6 h in MDA-MB-231 cells following treatment with DMSO control, CB-839 (300 nM) or K12 (10, 30 or 300 nM). Unlabelled metabolite (^12^C) is denoted by m+0, whilst m+x denotes a metabolite with x number of carbons labelled with ^13^C. **(F)** Total peak area of citrate. **(G)** Densitometry analysis of PDH1 and phospo-PDH1 in MDA-MB-231 cells treated with K12 (10, 30, 100 and 300 nM) for 1 h. Data show mean± SD from 3 independent experiments (Kruskal-Wallis with Dunns post-test; * = p<0.05). **(H)** Western blotting in MDA-MB-231 cells treated with K12 (300nM) for the indicated intervals.

## References

1. Bauer, K.R., Brown, M., Cress, R.D., Parise, C.A., and Caggiano, V. (2007). Descriptive analysis of estrogen receptor (ER)-negative, progesterone receptor (PR)-negative, and HER2-negative invasive breast cancer, the so-called triple-negative phenotype: a population-based study from the California cancer Registry. Cancer 109, 1721–1728. 10.1002/cncr.22618.

2. Trivers, K.F., Lund, M.J., Porter, P.L., Liff, J.M., Flagg, E.W., Coates, R.J., and Eley, J.W. (2009). The epidemiology of triple-negative breast cancer, including race. Cancer Causes Control 20, 1071–1082. 10.1007/s10552-009-9331-1.

3. Morrison, L., and Okines, A. (2023). Systemic Therapy for Metastatic Triple Negative Breast Cancer: Current Treatments and Future Directions. Cancers (Basel) 15. 10.3390/cancers15153801.

4. Foulkes, W.D., Smith, I.E., and Reis-Filho, J.S. (2010). Triple-negative breast cancer. N Engl J Med 363, 1938–1948. 10.1056/NEJMra1001389.

5. Wang, R., Zhu, Y., Liu, X., Liao, X., He, J., and Niu, L. (2019). The Clinicopathological features and survival outcomes of patients with different metastatic sites in stage IV breast cancer. BMC Cancer 19, 1091. 10.1186/s12885-019-6311-z.

6. Lin, N.U., Vanderplas, A., Hughes, M.E., Theriault, R.L., Edge, S.B., Wong, Y.N., Blayney, D.W., Niland, J.C., Winer, E.P., and Weeks, J.C. (2012). Clinicopathologic features, patterns of recurrence, and survival among women with triple-negative breast cancer in the National Comprehensive Cancer Network. Cancer 118, 5463–5472. 10.1002/cncr.27581.

7. Kesireddy, M., Elsayed, L., Shostrom, V.K., Agarwal, P., Asif, S., Yellala, A., and Krishnamurthy, J. (2024). Overall Survival and Prognostic Factors in Metastatic Triple-Negative Breast Cancer: A National Cancer Database Analysis. Cancers (Basel) 16. 10.3390/cancers16101791.

8. Insua-Rodriguez, J., Pein, M., Hongu, T., Meier, J., Descot, A., Lowy, C.M., De Braekeleer, E., Sinn, H.P., Spaich, S., Sutterlin, M., et al. (2018). Stress signaling in breast cancer cells induces matrix components that promote chemoresistant metastasis. EMBO Mol Med 10. 10.15252/emmm.201809003.

9. Semba, T., Wang, X., Xie, X., Cohen, E.N., Reuben, J.M., Dalby, K.N., Long, J.P., Phi, L.T.H., Tripathy, D., and Ueno, N.T. (2022). Identification of the JNK-Active Triple-Negative Breast Cancer Cluster Associated With an Immunosuppressive Tumor Microenvironment. J Natl Cancer Inst 114, 97–108. 10.1093/jnci/djab128.

10. Nasrazadani, A., and Van Den Berg, C.L. (2011). c-Jun N-terminal Kinase 2 Regulates Multiple Receptor Tyrosine Kinase Pathways in Mouse Mammary Tumor Growth and Metastasis. Genes Cancer 2, 31–45. 10.1177/1947601911400901.

11. Pein, M., Insua-Rodriguez, J., Hongu, T., Riedel, A., Meier, J., Wiedmann, L., Decker, K., Essers, M.A.G., Sinn, H.P., Spaich, S., et al. (2020). Metastasis-initiating cells induce and exploit a fibroblast niche to fuel malignant colonization of the lungs. Nat Commun 11, 1494. 10.1038/s41467-020-15188-x.

12. Cellurale, C., Girnius, N., Jiang, F., Cavanagh-Kyros, J., Lu, S., Garlick, D.S., Mercurio, A.M., and Davis, R.J. (2012). Role of JNK in mammary gland development and breast cancer. Cancer Res 72, 472–481. 10.1158/0008-5472.CAN-11-1628.

13. Girnius, N., and Davis, R.J. (2017). JNK Promotes Epithelial Cell Anoikis by Transcriptional and Post-translational Regulation of BH3-Only Proteins. Cell Rep 21, 1910–1921. 10.1016/j.celrep.2017.10.067.

14. Cellurale, C., Weston, C.R., Reilly, J., Garlick, D.S., Jerry, D.J., Sluss, H.K., and Davis, R.J. (2010). Role of JNK in a Trp53-dependent mouse model of breast cancer. PLoS One 5, e12469. 10.1371/journal.pone.0012469.

15. Girnius, N., Edwards, Y.J., Garlick, D.S., and Davis, R.J. (2018). The cJUN NH(2)-terminal kinase (JNK) signaling pathway promotes genome stability and prevents tumor initiation. Elife 7. 10.7554/eLife.36389.

16. Ashenden, M., van Weverwijk, A., Murugaesu, N., Fearns, A., Campbell, J., Gao, Q., Iravani, M., and Isacke, C.M. (2017). An In Vivo Functional Screen Identifies JNK Signaling As a Modulator of Chemotherapeutic Response in Breast Cancer. Mol Cancer Ther 16, 1967–1978. 10.1158/1535-7163.MCT-16-0731.

17. Latham, S.L., O’Donnell, Y.E.I., and Croucher, D.R. (2022). Non-kinase targeting of oncogenic c-Jun N-terminal kinase (JNK) signaling: the future of clinically viable cancer treatments. Biochem Soc Trans 50, 1823–1836. 10.1042/BST20220808.

18. Tan, E.Y., Yan, M., Campo, L., Han, C., Takano, E., Turley, H., Candiloro, I., Pezzella, F., Gatter, K.C., Millar, E.K., et al. (2009). The key hypoxia regulated gene CAIX is upregulated in basal-like breast tumours and is associated with resistance to chemotherapy. Br J Cancer 100, 405–411. 10.1038/sj.bjc.6604844.

19. Beckers, R.K., Selinger, C.I., Vilain, R., Madore, J., Wilmott, J.S., Harvey, K., Holliday, A., Cooper, C.L., Robbins, E., Gillett, D., et al. (2016). Programmed death ligand 1 expression in triple-negative breast cancer is associated with tumour-infiltrating lymphocytes and improved outcome. Histopathology 69, 25–34. 10.1111/his.12904.

20. Bjorkblom, B., Ostman, N., Hongisto, V., Komarovski, V., Filen, J.J., Nyman, T.A., Kallunki, T., Courtney, M.J., and Coffey, E.T. (2005). Constitutively active cytoplasmic c-Jun N-terminal kinase 1 is a dominant regulator of dendritic architecture: role of microtubule-associated protein 2 as an effector. J Neurosci 25, 6350–6361. 10.1523/JNEUROSCI.1517-05.2005.

21. Tararuk, T., Ostman, N., Li, W., Bjorkblom, B., Padzik, A., Zdrojewska, J., Hongisto, V., Herdegen, T., Konopka, W., Courtney, M.J., and Coffey, E.T. (2006). JNK1 phosphorylation of SCG10 determines microtubule dynamics and axodendritic length. J Cell Biol 173, 265–277. 10.1083/jcb.200511055.

22. Bjorkblom, B., Padzik, A., Mohammad, H., Westerlund, N., Komulainen, E., Hollos, P., Parviainen, L., Papageorgiou, A.C., Iljin, K., Kallioniemi, O., et al. (2012). c-Jun N-terminal kinase phosphorylation of MARCKSL1 determines actin stability and migration in neurons and in cancer cells. Mol Cell Biol 32, 3513–3526. 10.1128/MCB.00713-12.

23. Bryce, N.S., Failes, T.W., Stehn, J.R., Baker, K., Zahler, S., Arzhaeva, Y., Bischof, L., Lyons, C., Dedova, I., Arndt, G.M., et al. (2019). High-Content Imaging of Unbiased Chemical Perturbations Reveals that the Phenotypic Plasticity of the Actin Cytoskeleton Is Constrained. Cell Syst 9, 496–507 e495. 10.1016/j.cels.2019.09.002.

24. Regot, S., Hughey, J.J., Bajar, B.T., Carrasco, S., and Covert, M.W. (2014). High-sensitivity measurements of multiple kinase activities in live single cells. Cell 157, 1724–1734. 10.1016/j.cell.2014.04.039.

25. Quek, L.E., van Geldermalsen, M., Guan, Y.F., Wahi, K., Mayoh, C., Balaban, S., Pang, A., Wang, Q., Cowley, M.J., Brown, K.K., et al. (2022). Glutamine addiction promotes glucose oxidation in triple-negative breast cancer. Oncogene 41, 4066–4078. 10.1038/s41388-022-02408-5.

26. van Geldermalsen, M., Wang, Q., Nagarajah, R., Marshall, A.D., Thoeng, A., Gao, D., Ritchie, W., Feng, Y., Bailey, C.G., Deng, N., et al. (2016). ASCT2/SLC1A5 controls glutamine uptake and tumour growth in triple-negative basal-like breast cancer. Oncogene 35, 3201–3208. 10.1038/onc.2015.381.

27. Wang, J.B., Erickson, J.W., Fuji, R., Ramachandran, S., Gao, P., Dinavahi, R., Wilson, K.F., Ambrosio, A.L., Dias, S.M., Dang, C.V., and Cerione, R.A. (2010). Targeting mitochondrial glutaminase activity inhibits oncogenic transformation. Cancer Cell 18, 207–219. 10.1016/j.ccr.2010.08.009.

28. Cassago, A., Ferreira, A.P., Ferreira, I.M., Fornezari, C., Gomes, E.R., Greene, K.S., Pereira, H.M., Garratt, R.C., Dias, S.M., and Ambrosio, A.L. (2012). Mitochondrial localization and structure-based phosphate activation mechanism of Glutaminase C with implications for cancer metabolism. Proc Natl Acad Sci U S A 109, 1092–1097. 10.1073/pnas.1112495109.

29. Huang, Q., Stalnecker, C., Zhang, C., McDermott, L.A., Iyer, P., O’Neill, J., Reimer, S., Cerione, R.A., and Katt, W.P. (2018). Characterization of the interactions of potent allosteric inhibitors with glutaminase C, a key enzyme in cancer cell glutamine metabolism. J Biol Chem 293, 3535–3545. 10.1074/jbc.M117.810101.

30. Vogl, D.T., Younes, A., Stewart, K., Orford, K.W., Bennett, M., Siegel, D., and Berdeja, J.G. (2015). Phase 1 Study of CB-839, a First-in-Class, Glutaminase Inhibitor in Patients with Multiple Myeloma and Lymphoma. Blood 126, 3059–3059. 10.1182/blood.V126.23.3059.3059.

31. Wang, E.S., Frankfurt, O., Orford, K.W., Bennett, M., Flinn, I.W., Maris, M., and Konopleva, M. (2015). Phase 1 Study of CB-839, a First-in-Class, Orally Administered Small Molecule Inhibitor of Glutaminase in Patients with Relapsed/Refractory Leukemia. Blood 126, 2566–2566. 10.1182/blood.V126.23.2566.2566.

32. Tannir, N.M., Agarwal, N., Porta, C., Lawrence, N.J., Motzer, R., McGregor, B., Lee, R.J., Jain, R.K., Davis, N., Appleman, L.J., et al. (2022). Efficacy and Safety of Telaglenastat Plus Cabozantinib vs Placebo Plus Cabozantinib in Patients With Advanced Renal Cell Carcinoma: The CANTATA Randomized Clinical Trial. JAMA Oncol 8, 1411–1418. 10.1001/jamaoncol.2022.3511.

33. Kolobova, E., Tuganova, A., Boulatnikov, I., and Popov, K.M. (2001). Regulation of pyruvate dehydrogenase activity through phosphorylation at multiple sites. Biochem J 358, 69–77. 10.1042/0264-6021:3580069.

34. Singleton, D.C., Dechaume, A.L., Murray, P.M., Katt, W.P., Baguley, B.C., and Leung, E.Y. (2020). Pyruvate anaplerosis is a mechanism of resistance to pharmacological glutaminase inhibition in triple-receptor negative breast cancer. BMC Cancer 20, 470. 10.1186/s12885-020-06885-3.

35. Itah, Z., Chaudhry, S., Raju Ponny, S., Aydemir, O., Lee, A., Cavanagh-Kyros, J., Tournier, C., Muller, W.J., and Davis, R.J. (2023). HER2-driven breast cancer suppression by the JNK signaling pathway. Proc Natl Acad Sci U S A 120, e2218373120. 10.1073/pnas.2218373120.

36. Xie, X., Kaoud, T.S., Edupuganti, R., Zhang, T., Kogawa, T., Zhao, Y., Chauhan, G.B., Giannoukos, D.N., Qi, Y., Tripathy, D., et al. (2017). c-Jun N-terminal kinase promotes stem cell phenotype in triple-negative breast cancer through upregulation of Notch1 via activation of c-Jun. Oncogene 36, 2599–2608. 10.1038/onc.2016.417.

37. Plantevin Krenitsky, V., Nadolny, L., Delgado, M., Ayala, L., Clareen, S.S., Hilgraf, R., Albers, R., Hegde, S., D’Sidocky, N., Sapienza, J., et al. (2012). Discovery of CC-930, an orally active anti-fibrotic JNK inhibitor. Bioorg Med Chem Lett 22, 1433–1438. 10.1016/j.bmcl.2011.12.027.

38. Nagy, M.A., Hilgraf, R., Mortensen, D.S., Elsner, J., Norris, S., Tikhe, J., Yoon, W., Paisner, D., Delgado, M., Erdman, P., et al. (2021). Discovery of the c-Jun N-Terminal Kinase Inhibitor CC-90001. J Med Chem 64, 18193–18208. 10.1021/acs.jmedchem.1c01716.

39. Sabapathy, K., Hochedlinger, K., Nam, S.Y., Bauer, A., Karin, M., and Wagner, E.F. (2004). Distinct roles for JNK1 and JNK2 in regulating JNK activity and c-Jun-dependent cell proliferation. Mol Cell 15, 713–725. 10.1016/j.molcel.2004.08.028.

40. Tafolla, E., Wang, S., Wong, B., Leong, J., and Kapila, Y.L. (2005). JNK1 and JNK2 oppositely regulate p53 in signaling linked to apoptosis triggered by an altered fibronectin matrix: JNK links FAK and p53. J Biol Chem 280, 19992–19999. 10.1074/jbc.M500331200.

41. Tuncman, G., Hirosumi, J., Solinas, G., Chang, L., Karin, M., and Hotamisligil, G.S. (2006). Functional in vivo interactions between JNK1 and JNK2 isoforms in obesity and insulin resistance. Proc Natl Acad Sci U S A 103, 10741–10746. 10.1073/pnas.0603509103.

42. Hui, L., Zatloukal, K., Scheuch, H., Stepniak, E., and Wagner, E.F. (2008). Proliferation of human HCC cells and chemically induced mouse liver cancers requires JNK1-dependent p21 downregulation. J Clin Invest 118, 3943–3953. 10.1172/JCI37156.

43. She, Q.B., Chen, N., Bode, A.M., Flavell, R.A., and Dong, Z. (2002). Deficiency of c-Jun-NH(2)-terminal kinase-1 in mice enhances skin tumor development by 12-O-tetradecanoylphorbol-13-acetate. Cancer Res 62, 1343–1348.

44. Sakurai, T., Maeda, S., Chang, L., and Karin, M. (2006). Loss of hepatic NF-kappa B activity enhances chemical hepatocarcinogenesis through sustained c-Jun N-terminal kinase 1 activation. Proc Natl Acad Sci U S A 103, 10544–10551. 10.1073/pnas.0603499103.

45. Davis, R.J. (2000). Signal transduction by the JNK group of MAP kinases. Cell 103, 239–252. 10.1016/s0092-8674(00)00116-1.

46. Jaeschke, A., Karasarides, M., Ventura, J.J., Ehrhardt, A., Zhang, C., Flavell, R.A., Shokat, K.M., and Davis, R.J. (2006). JNK2 is a positive regulator of the cJun transcription factor. Mol Cell 23, 899–911. 10.1016/j.molcel.2006.07.028.

47. Wang, X., Chao, L., Li, X., Ma, G., Chen, L., Zang, Y., and Zhou, G. (2010). Elevated expression of phosphorylated c-Jun NH2-terminal kinase in basal-like and “triple-negative” breast cancers. Hum Pathol 41, 401–406. 10.1016/j.humpath.2009.08.018.

48. Bjorkblom, B., Vainio, J.C., Hongisto, V., Herdegen, T., Courtney, M.J., and Coffey, E.T. (2008). All JNKs can kill, but nuclear localization is critical for neuronal death. J Biol Chem 283, 19704–19713. 10.1074/jbc.M707744200.

49. Oliva, A.A., Jr., Atkins, C.M., Copenagle, L., and Banker, G.A. (2006). Activated c-Jun N-terminal kinase is required for axon formation. J Neurosci 26, 9462–9470. 10.1523/JNEUROSCI.2625-06.2006.

50. Swinney, D.C., and Anthony, J. (2011). How were new medicines discovered? Nat Rev Drug Discov 10, 507–519. 10.1038/nrd3480.

51. Vincent, F., Nueda, A., Lee, J., Schenone, M., Prunotto, M., and Mercola, M. (2022). Phenotypic drug discovery: recent successes, lessons learned and new directions. Nat Rev Drug Discov 21, 899–914. 10.1038/s41573-022-00472-w.

52. Papa, S., Choy, P.M., and Bubici, C. (2019). The ERK and JNK pathways in the regulation of metabolic reprogramming. Oncogene 38, 2223–2240. 10.1038/s41388-018-0582-8.

53. Zhou, Q., Lam, P.Y., Han, D., and Cadenas, E. (2008). c-Jun N-terminal kinase regulates mitochondrial bioenergetics by modulating pyruvate dehydrogenase activity in primary cortical neurons. J Neurochem 104, 325–335. 10.1111/j.1471-4159.2007.04957.x.

54. Zhou, Q., Lam, P.Y., Han, D., and Cadenas, E. (2009). Activation of c-Jun-N-terminal kinase and decline of mitochondrial pyruvate dehydrogenase activity during brain aging. FEBS Lett 583, 1132–1140. 10.1016/j.febslet.2009.02.043.

55. Korotchkina, L.G., and Patel, M.S. (2001). Site specificity of four pyruvate dehydrogenase kinase isoenzymes toward the three phosphorylation sites of human pyruvate dehydrogenase. J Biol Chem 276, 37223–37229. 10.1074/jbc.M103069200.

56. Lukey, M.J., Greene, K.S., Erickson, J.W., Wilson, K.F., and Cerione, R.A. (2016). The oncogenic transcription factor c-Jun regulates glutaminase expression and sensitizes cells to glutaminase-targeted therapy. Nat Commun 7, 11321. 10.1038/ncomms11321.

57. Choudhury, D., Rong, N., Ikhapoh, I., Rajabian, N., Tseropoulos, G., Wu, Y., Mehrotra, P., Thiyagarajan, R., Shahini, A., Seldeen, K.L., et al. (2022). Inhibition of glutaminolysis restores mitochondrial function in senescent stem cells. Cell Rep 41, 111744. 10.1016/j.celrep.2022.111744.

58. DeMichele, A., Harding, J.J., Telli, M.L., Munster, P.N., McKay, R., Iliopoulos, O., Orford, K.W., Bennett, M.K., Mier, J.W., Owonikoko, T.K., et al. (2016). Phase 1 study of CB-839, a small molecule inhibitor of glutaminase (GLS) in combination with paclitaxel (Pac) in patients (pts) with triple negative breast cancer (TNBC). Journal of Clinical Oncology 34, 1011–1011. 10.1200/JCO.2016.34.15_suppl.1011.

59. Harding, J.J., Telli, M., Munster, P., Voss, M.H., Infante, J.R., DeMichele, A., Dunphy, M., Le, M.H., Molineaux, C., Orford, K., et al. (2021). A Phase I Dose-Escalation and Expansion Study of Telaglenastat in Patients with Advanced or Metastatic Solid Tumors. Clin Cancer Res 27, 4994–5003. 10.1158/1078-0432.CCR-21-1204.

60. Herr, R., Wohrle, F.U., Danke, C., Berens, C., and Brummer, T. (2011). A novel MCF-10A line allowing conditional oncogene expression in 3D culture. Cell Commun Signal 9, 17. 10.1186/1478-811X-9-17.

61. Dull, T., Zufferey, R., Kelly, M., Mandel, R.J., Nguyen, M., Trono, D., and Naldini, L. (1998). A third-generation lentivirus vector with a conditional packaging system. J Virol 72, 8463–8471. 10.1128/JVI.72.11.8463-8471.1998.

62. Komatsu, N., Aoki, K., Yamada, M., Yukinaga, H., Fujita, Y., Kamioka, Y., and Matsuda, M. (2011). Development of an optimized backbone of FRET biosensors for kinases and GTPases. Mol Biol Cell 22, 4647–4656. 10.1091/mbc.E11-01-0072.

63. Niwa, H., Yamamura, K., and Miyazaki, J. (1991). Efficient selection for high-expression transfectants with a novel eukaryotic vector. Gene 108, 193–199. 10.1016/0378-1119(91)90434-d.

64. Timpson, P., McGhee, E.J., Erami, Z., Nobis, M., Quinn, J.A., Edward, M., and Anderson, K.I. (2011). Organotypic collagen I assay: a malleable platform to assess cell behaviour in a 3-dimensional context. J Vis Exp, e3089. 10.3791/3089.

65. Perez-Riverol, Y., Bandla, C., Kundu, D.J., Kamatchinathan, S., Bai, J., Hewapathirana, S., John, N.S., Prakash, A., Walzer, M., Wang, S., and Vizcaino, J.A. (2025). The PRIDE database at 20 years: 2025 update. Nucleic Acids Res 53, D543–D553. 10.1093/nar/gkae1011.

66. Li, B., and Dewey, C.N. (2011). RSEM: accurate transcript quantification from RNA-Seq data with or without a reference genome. BMC Bioinformatics 12, 323. 10.1186/1471-2105-12-323.

67. Hastings, J.F., Latham, S.L., Kamili, A., Wheatley, M.S., Han, J.Z.R., Wong-Erasmus, M., Phimmachanh, M., Nobis, M., Pantarelli, C., Cadell, A.L., et al. (2023). Memory of stochastic single-cell apoptotic signaling promotes chemoresistance in neuroblastoma. Sci Adv 9, eabp8314. 10.1126/sciadv.abp8314.

68. Warren, S.C., Margineanu, A., Alibhai, D., Kelly, D.J., Talbot, C., Alexandrov, Y., Munro, I., Katan, M., Dunsby, C., and French, P.M. (2013). Rapid global fitting of large fluorescence lifetime imaging microscopy datasets. PLoS One 8, e70687. 10.1371/journal.pone.0070687.

69. Conway, J.R.W., Warren, S.C., Lee, Y.K., McCulloch, A.T., Magenau, A., Lee, V., Metcalf, X.L., Stoehr, J., Haigh, K., Abdulkhalek, L., et al. (2023). Monitoring AKT activity and targeting in live tissue and disease contexts using a real-time Akt-FRET biosensor mouse. Sci Adv 9, eadf9063. 10.1126/sciadv.adf9063.

70. El-Sayed, H.A., A., S.S., H., M.A., M., B.M., and and Abdel-Kader, R.T. (2016). Synthesis and Biological Evaluation of 2-Oxo/Thioxoquinoxaline and 2-Oxo/Thioxoquinoxaline-Based Nucleoside Analogues. Nucleosides Nucleotides Nucl. Acids 35, 16–31. 10.1080/15257770.2015.1114124.

71. Punna, S., Kaltgrad, E., and Finn, M.G. (2005). “Clickable” Agarose for Affinity Chromatography. Bioconjugate Chemistry 16, 1536–1541. 10.1021/bc0501496.

72. Tyanova, S., Temu, T., Sinitcyn, P., Carlson, A., Hein, M.Y., Geiger, T., Mann, M., and Cox, J. (2016). The Perseus computational platform for comprehensive analysis of (prote)omics data. Nat Methods 13, 731–740. 10.1038/nmeth.3901.

73. Huang, Q., Katt, W.P., McDermott, L.A., Cerione, R.A. (2016). Crystal structure of the clinically relevant glutaminase inhibitor CB-839 in complex with glutaminase C.

74. Friesner, R.A., Banks, J.L., Murphy, R.B., Halgren, T.A., Klicic, J.J., Mainz, D.T., Repasky, M.P., Knoll, E.H., Shelley, M., Perry, J.K., et al. (2004). Glide:_ A New Approach for Rapid, Accurate Docking and Scoring. 1. Method and Assessment of Docking Accuracy. J. Med. Chem. 47, 1739–1749. 10.1021/jm0306430.

75. Halgren, T.A., Murphy, R.B., Friesner, R.A., Beard, H.S., Frye, L.L., Pollard, W.T., and Banks, J.L. (2004). Glide:_ A New Approach for Rapid, Accurate Docking and Scoring. 2. Enrichment Factors in Database Screening. J. Med. Chem. 47, 1750–1759. 10.1021/jm030644s.

76. Friesner, R.A., Murphy, R.B., Repasky, M.P., Frye, L.L., Greenwood, J.R., Halgren, T.A., Sanschagrin, P.C., and Mainz, D.T. (2006). Extra Precision Glide:_ Docking and Scoring Incorporating a Model of Hydrophobic Enclosure for Protein−Ligand Complexes. J. Med. Chem. 49, 6177–6196. 10.1021/jm051256o.

77. Harder, E., Damm, W., Maple, J., Wu, C., Reboul, M., Xiang, J.Y., Wang, L., Lupyan, D., Dahlgren, M.K., Knight, J.L., et al. (2016). OPLS3: A Force Field Providing Broad Coverage of Drug-like Small Molecules and Proteins. J. Chem. Theory Comput. 12, 281–296. 10.1021/acs.jctc.5b00864.

78. Shelley, J.C., Cholleti, A., Frye, L.L., Greenwood, J.R., Timlin, M.R., and Uchimaya, M. (2007). Epik: a software program for pKaprediction and protonation state generation for drug-like molecules. Journal of Computer-Aided Molecular Design 21, 681–691. 10.1007/s10822-007-9133-z.

79. Michel, J., and Essex, J.W. (2010). Prediction of protein–ligand binding affinity by free energy simulations: assumptions, pitfalls and expectations. Journal of Computer-Aided Molecular Design 24, 639–658. 10.1007/s10822-010-9363-3.

80. Wahi, K., Freidman, N., Wang, Q., Devadason, M., Quek, L.E., Pang, A., Lloyd, L., Larance, M., Zanini, F., Harvey, K., et al. (2024). Macropinocytosis mediates resistance to loss of glutamine transport in triple-negative breast cancer. EMBO J 43, 5857–5882. 10.1038/s44318-024-00271-6.

81. Wang, Q., Guan, Y.F., Hancock, S.E., Wahi, K., van Geldermalsen, M., Zhang, B.K., Pang, A., Nagarajah, R., Mak, B., Freidman, N., et al. (2021). Inhibition of guanosine monophosphate synthetase (GMPS) blocks glutamine metabolism and prostate cancer growth. J Pathol 254, 135–146. 10.1002/path.5665.

82. Lu, W., Clasquin, M.F., Melamud, E., Amador-Noguez, D., Caudy, A.A., and Rabinowitz, J.D. (2010). Metabolomic analysis via reversed-phase ion-pairing liquid chromatography coupled to a stand alone orbitrap mass spectrometer. Anal Chem 82, 3212–3221. 10.1021/ac902837x.

83. Chambers, M.C., Maclean, B., Burke, R., Amodei, D., Ruderman, D.L., Neumann, S., Gatto, L., Fischer, B., Pratt, B., Egertson, J., et al. (2012). A cross-platform toolkit for mass spectrometry and proteomics. Nat Biotechnol 30, 918–920. 10.1038/nbt.2377.

