## Supplementary figures and images for "A selective inhibitor of oncogenic JNK signalling perturbs metastatic outgrowth of triple-negative breast cancer through metabolic blockade"

### Supplementary Figures

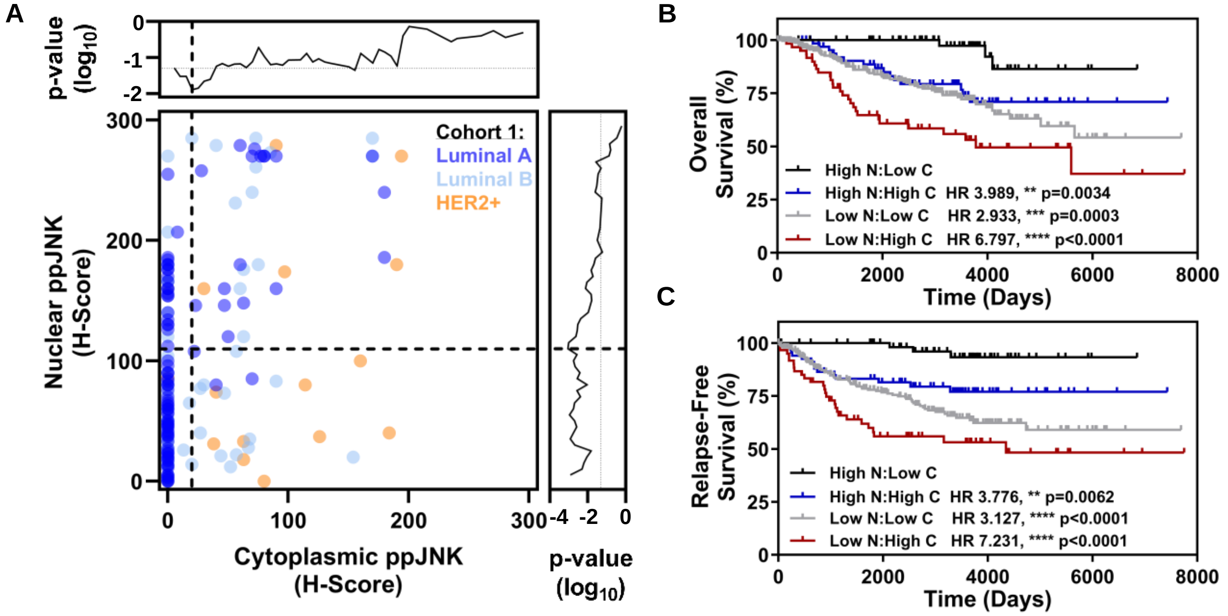

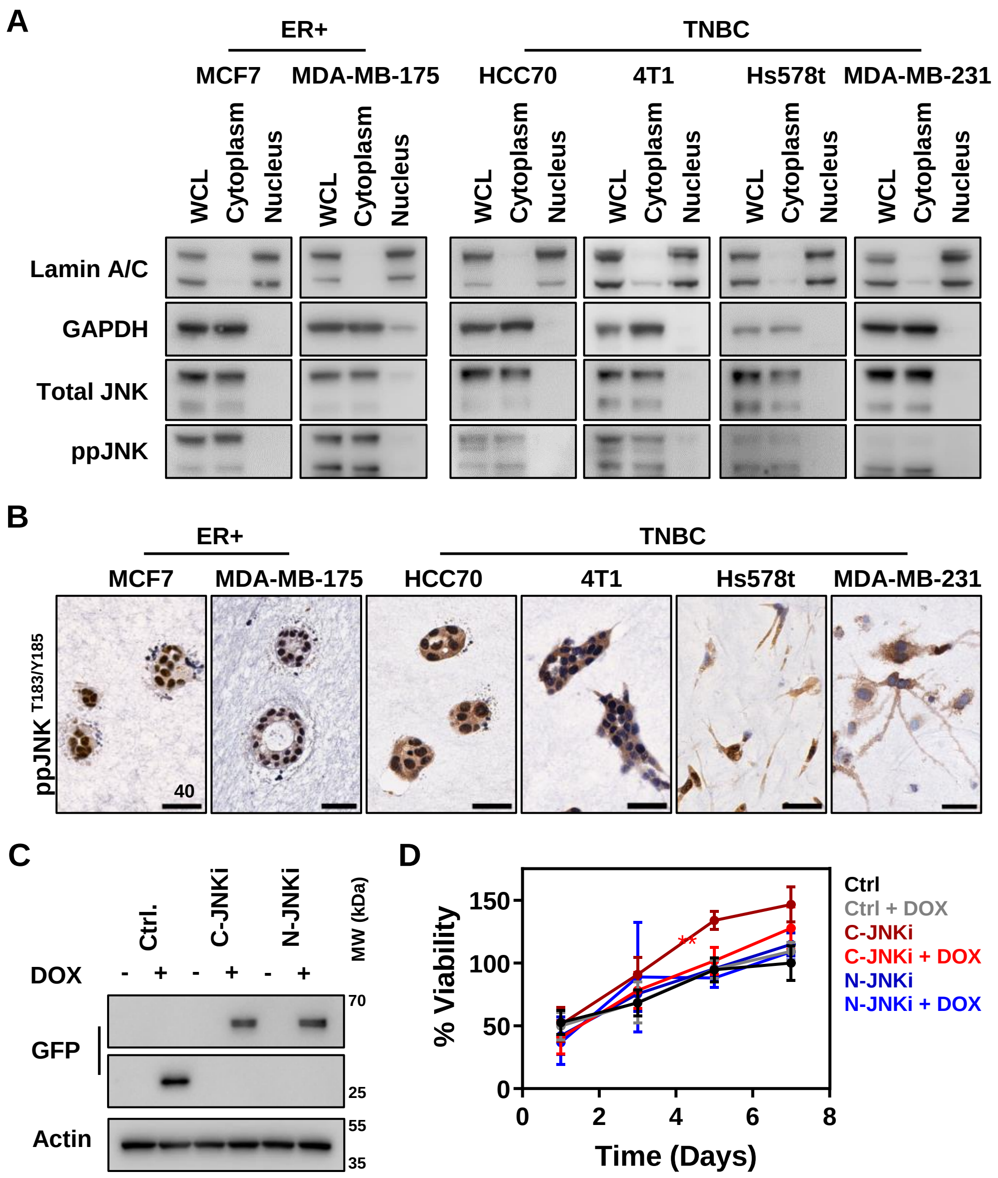

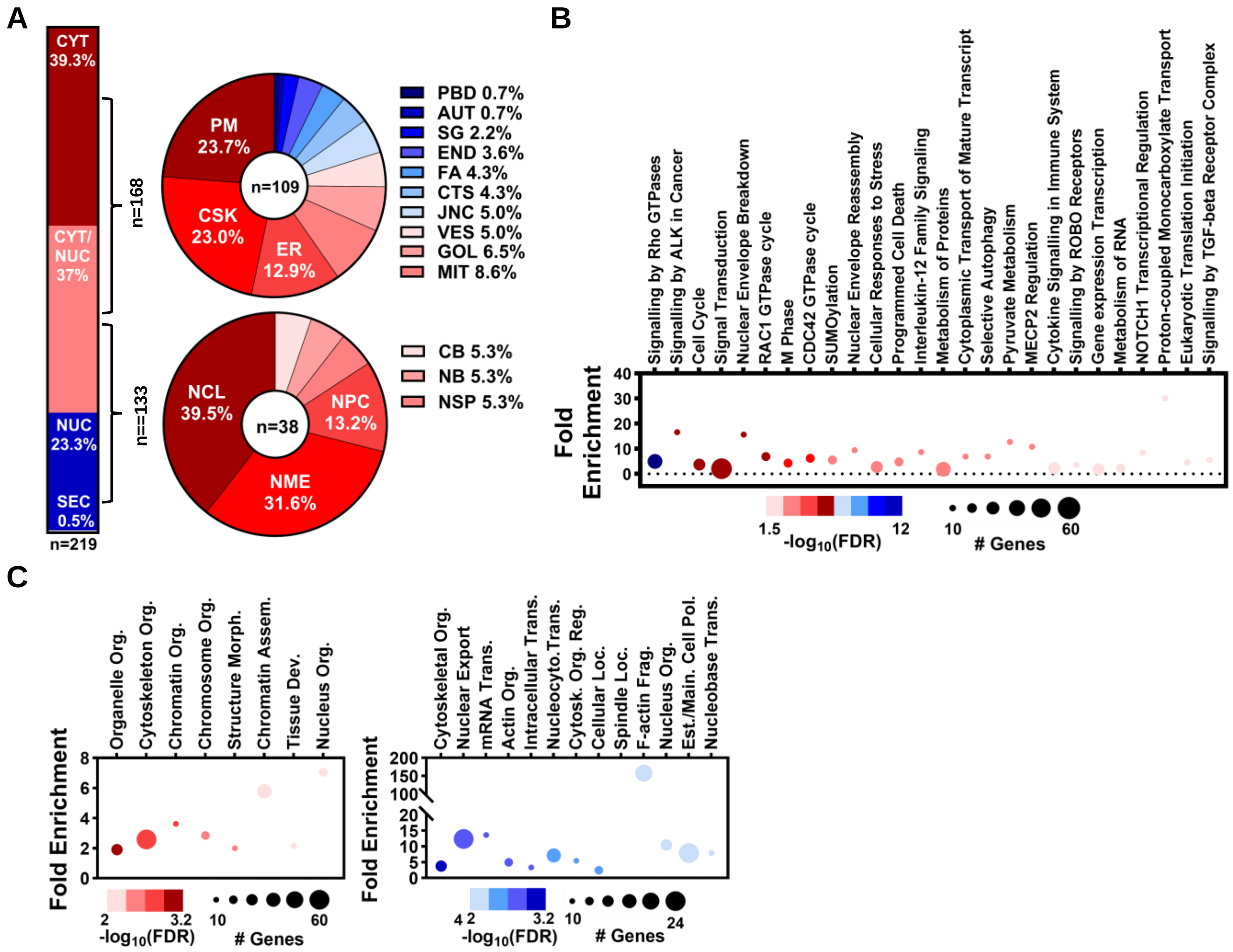

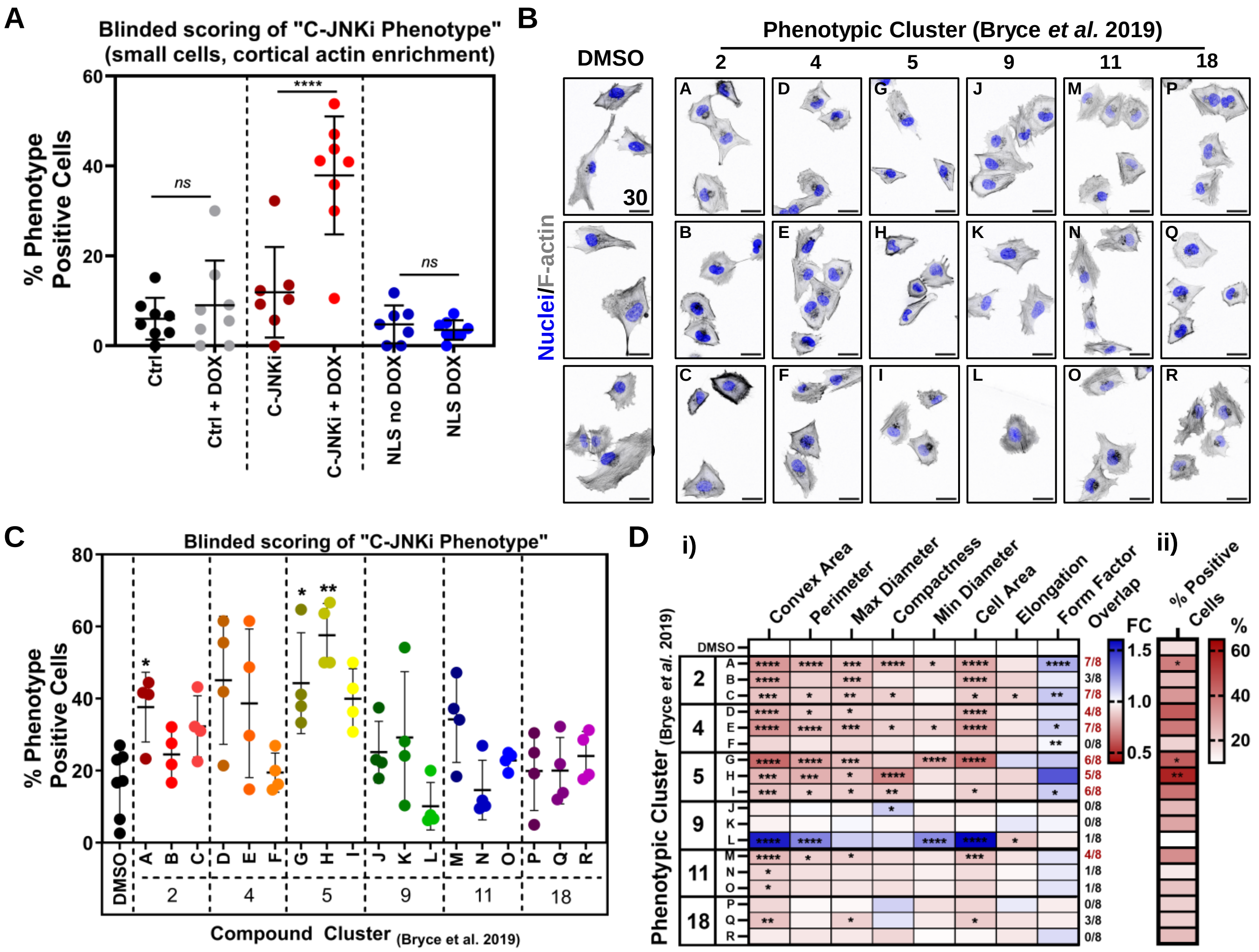

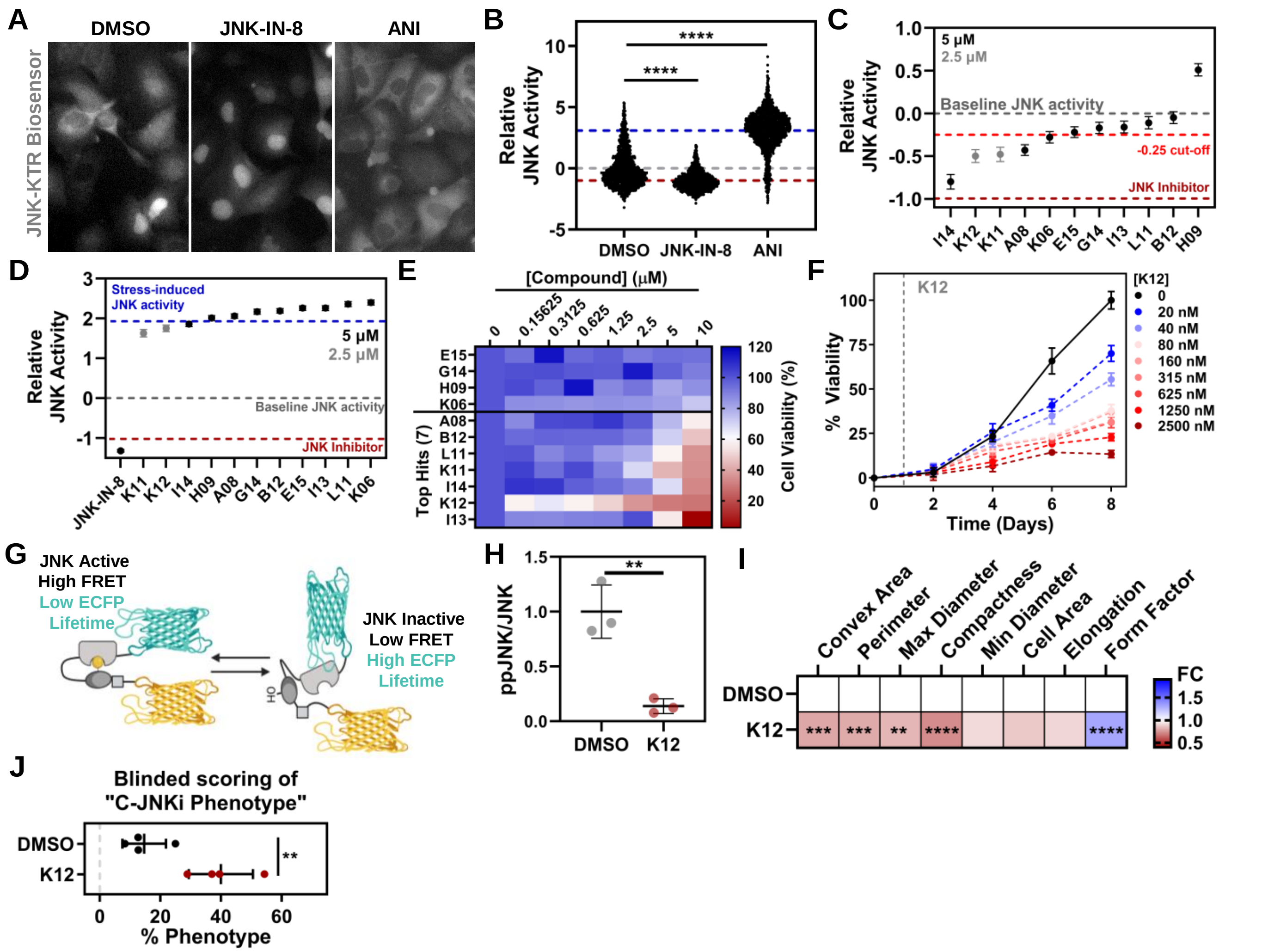

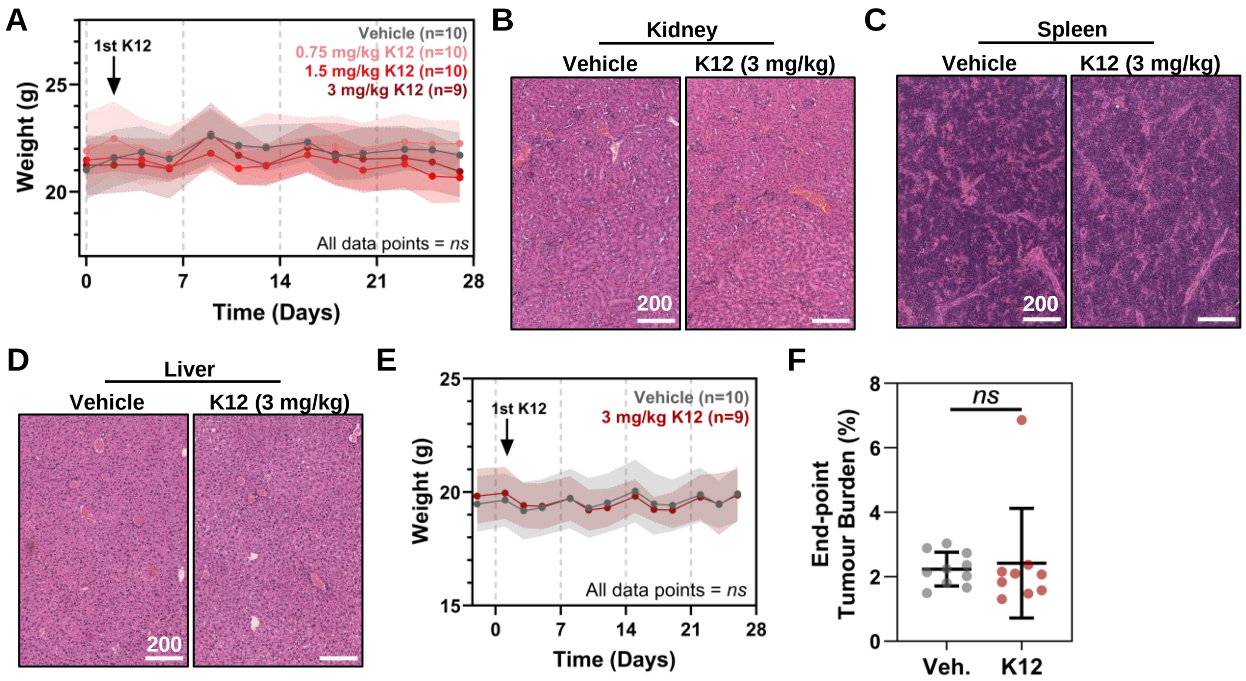

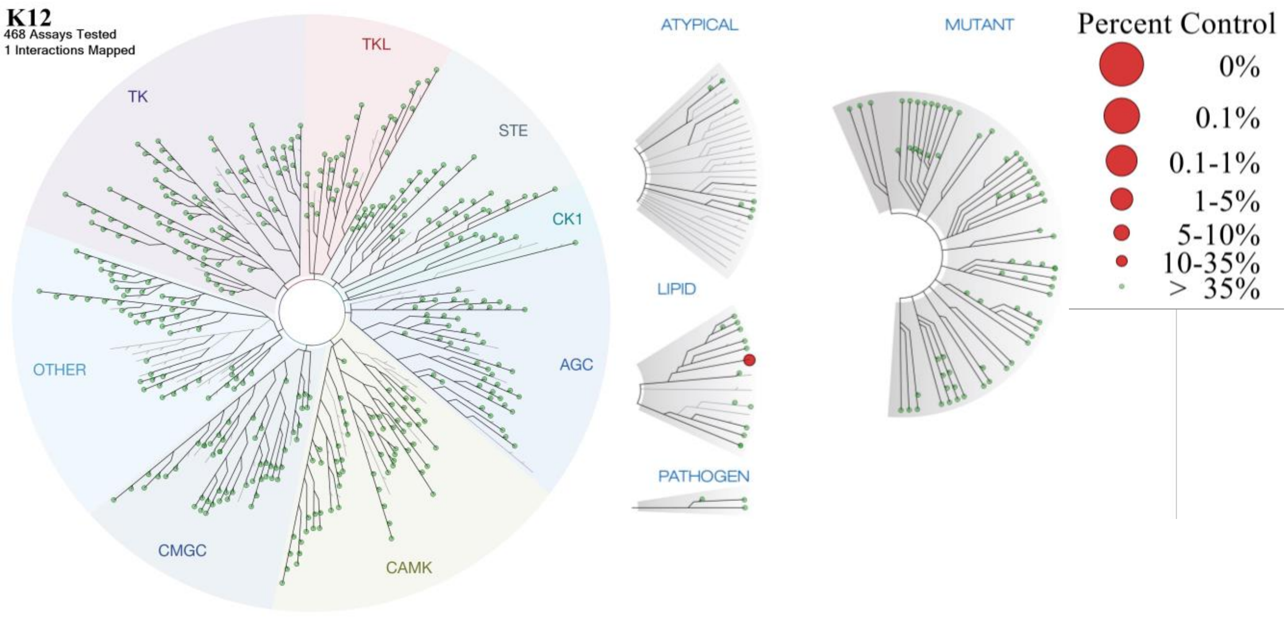

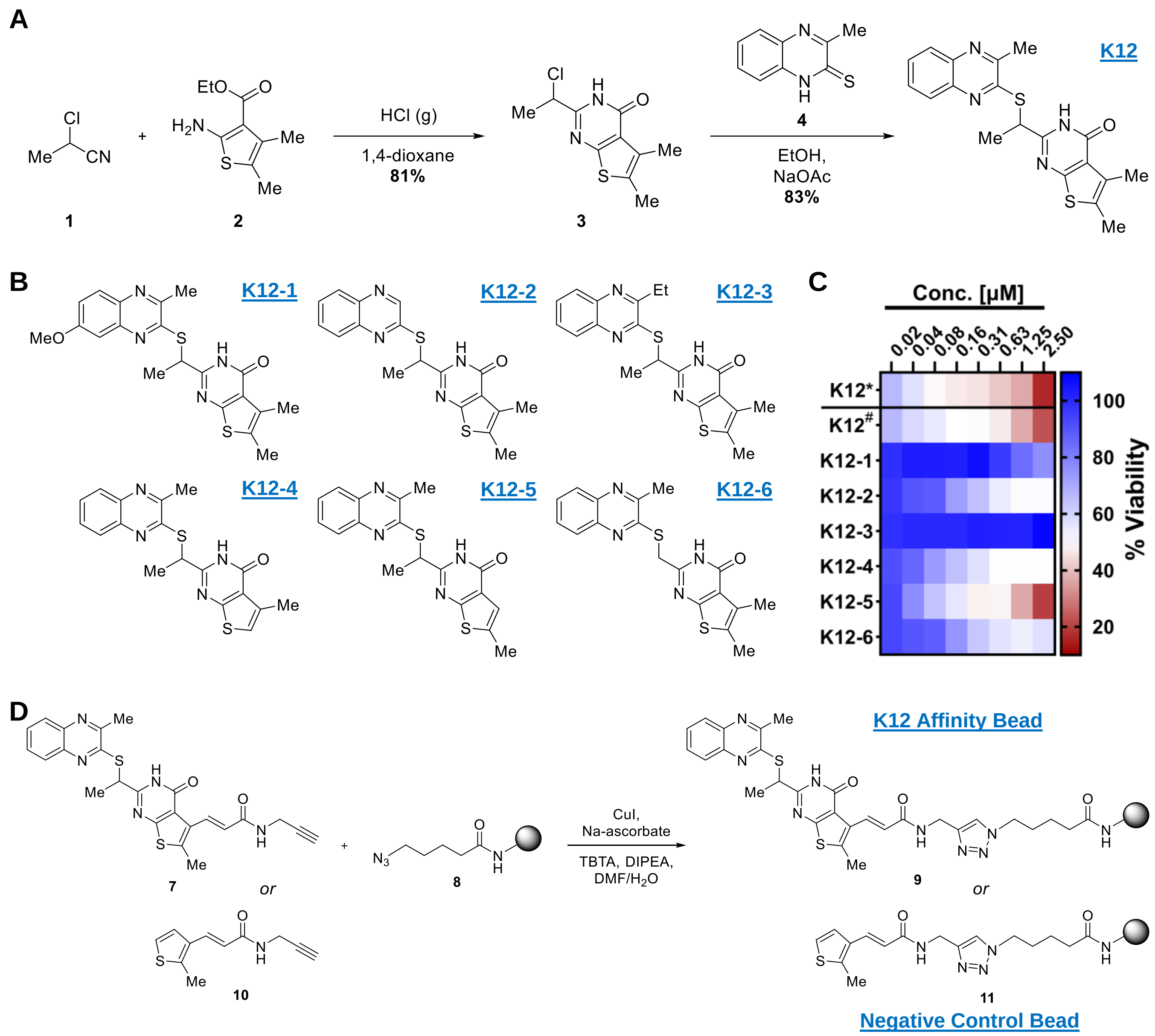

#### FIGURE S9

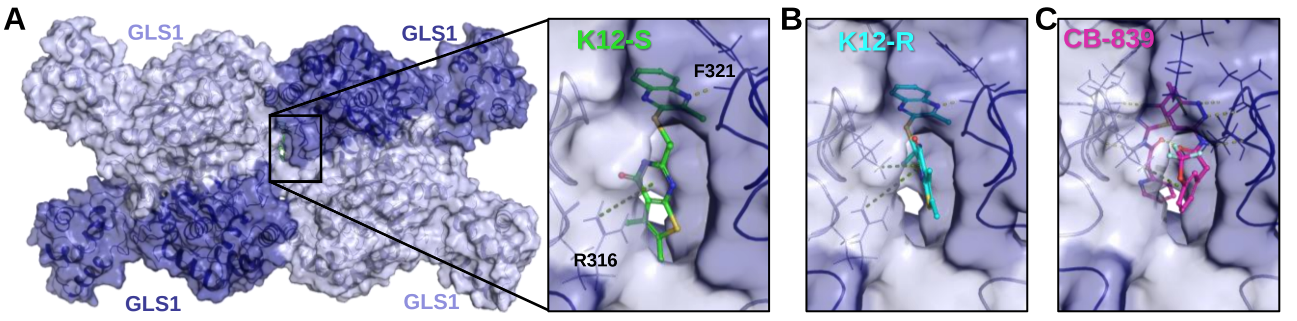

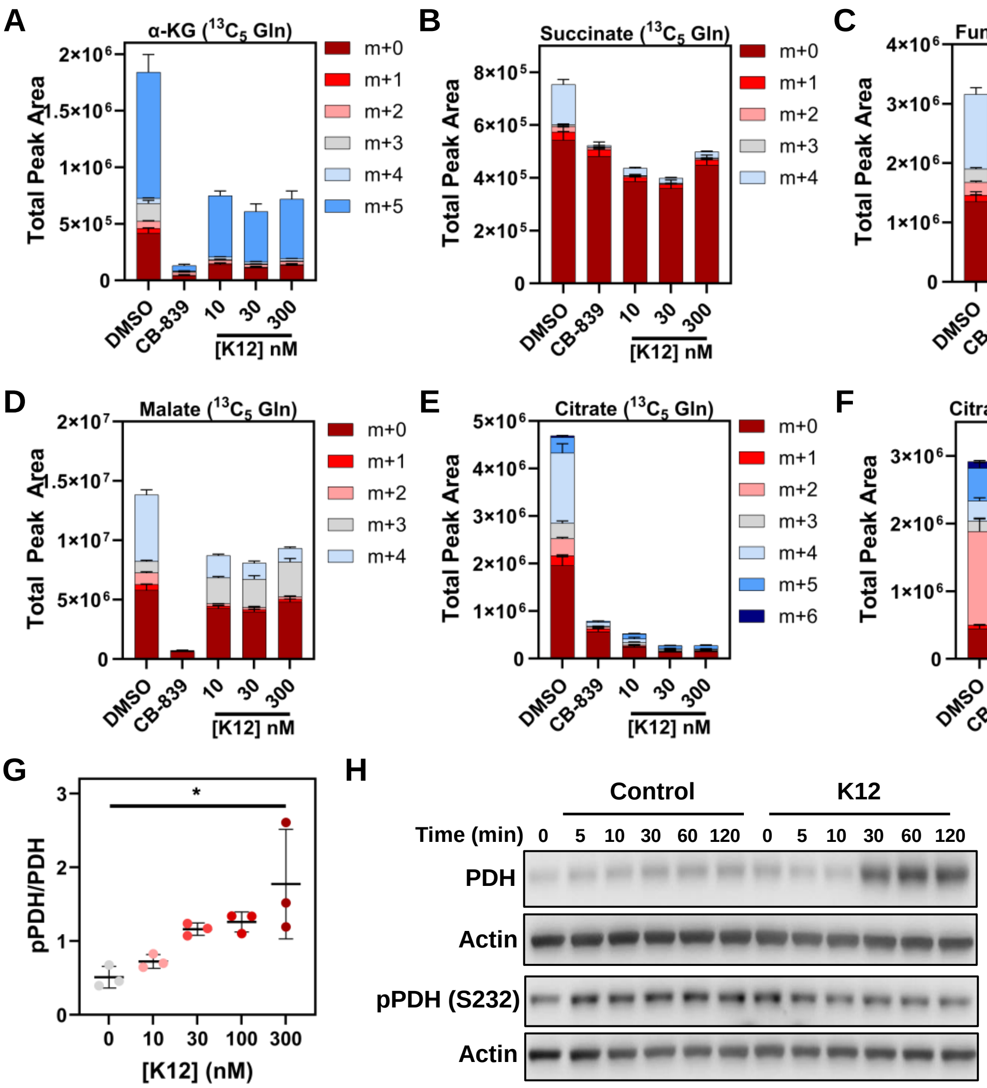
