## Supplementary Methods for "A selective inhibitor of oncogenic JNK signalling perturbs metastatic outgrowth of triple-negative breast cancer through metabolic blockade"

### Supplementary Methods – Chemical Synthesis

#### General Experimental

Melting points were obtained on an OptiMelt Automated Melting Point System with Digital Image Processing Technology and are uncorrected. $^{1}H$ NMR and $^{13}C$ NMR were recorded on a Bruker Avance III 400 (400 MHz) or Bruker Avance III 600 (600 MHz), with data acquired and processed using TopSpin 4.4 software. Chemical shifts are expressed in parts per million (ppm) on the δ scale. Chemical shifts in (a) $CDCl_3$ were referenced relative to $CHCl_3$ (7.26 ppm) for $^{1}H$ NMR and $CDCl_3$ (77.16 ppm) for $^{13}C$ NMR and (b) $(CD_3)_2SO$ were referenced relative to $(CH_3)_2SO$ (2.50 ppm) for $^{1}H$ NMR and $(CD_3)_2SO$ (39.52 ppm) for $^{13}C$ NMR.[10] Infrared spectra were obtained on a Cary 630 FTIR spectrophotometer with ATR and are reported in wavenumbers ($cm^{-1}$). Spectra were recorded from neat samples. HRMS were performed on an Orbitrap LTQ XL (Thermo Fisher Scientific, San Jose, Ca, USA) ion trap mass spectrometer using a nanospray (nano-electrospray) ionization source to generate ions from the analyte in solution. The analysis was carried out in positive ion mode using the orbitrap FTMS analyser at a resolution of 100000.

Unless otherwise stated all reactions were performed in flame dried glassware under an atmosphere of high purity argon using dry solvents. Reagents and solvents were purchased from commercial sources and used without further purification. 3-Methylquinoxaline-2(1*H*)-thione (**4**)[1], quinoxaline-2(1*H*)-thione (**12**)[11], 3-ethylquinoxaline-2(1*H*)-thione (**13**)[12] and 2-(chloromethyl)-5,6-dimethylthieno[2,3-*d*]pyrimidin-4(3*H*)-one (**14**)[13] were synthesised according to literature procedures.

#### Synthesis of K12 and K12 analogues

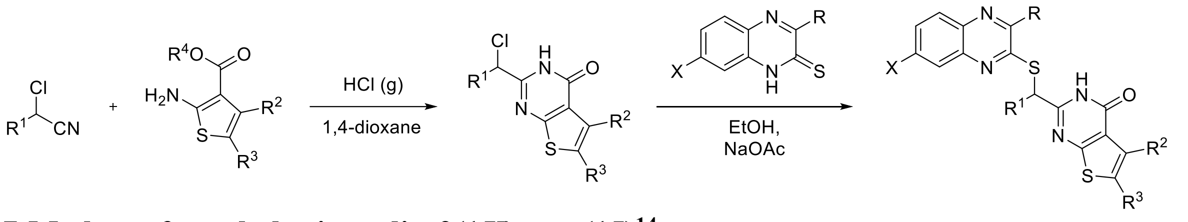


##### 7-Methoxy-3-methylquinoxalin-2(1*H*)-one (15)[14]

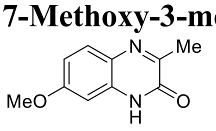


**15**

Pyruvic acid (279 mg, 3.17 mmol) was added to a suspension of 4-methoxybenzene-1,2-diamine (365 mg, 2.64 mmol) in aq. $H_2SO_4$ (1.8 M, 3.6 mL) and the reaction mixture was stirred at room temperature. After 24 h, the reaction mixture was neutralised with 1 M aqueous NaOH solution before the resulting precipitate was collected via vacuum filtration and washed with $H_2O$. The crude material was purified by flash chromatography eluting with 4% methanol/dichloromethane, to afford the product **15** as a brown solid (247 mg, 49%) with all analytical data matching that reported in the literature.[14] Mp. 220-230 °C (lit.[15] 217-218 °C); $^{1}H$ NMR (400 MHz, DMSO-$d_6$) δ 2.34 (s, 3H), 3.81 (s, 3H), 6.72 – 6.75 (m, 1H), 6.84 – 6.89 (m, 1H), 7.58 – 7.63 (m, 1H),12.19 (s, 1H).

##### 7-Methoxy-3-methylquinoxaline-2(1*H*)-thione (16)

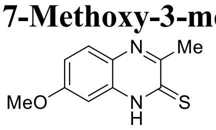


**16**

$P_2S_5$ (140 mg, 0.631 mmol) was suspended in dry pyridine (6 mL) and 7-methoxy-3-methylquinoxalin-2(1*H*)-one **15** (103 mg, 0.542 mmol) was added. The reaction mixture was stirred at reflux for 4 h. The reaction mixture was diluted with warm $H_2O$ and extracted with ethyl acetate (x5). Combined organic extracts were washed with brine (x2), dried ($Na_2SO_4$), and concentrated *in vacuo* to afford the thione **16** as a brown solid (112 mg, quant.). Mp. 245 °C (dec); $^{1}H$ NMR (400 MHz, DMSO-$d_6$) δ 2.59 (s, 3H), 3.85 (s, 3H), 7.02 – 7.10 (m, 2H), 7.71 – 7.75 (m, 1H),

14.23 (s, 1H); $^{13}$C NMR (101 MHz, DMSO-$d_6$) δ 24.3, 55.7, 97.5, 114.9, 129.3, 130.2, 133.0, 158.2, 160.4, 174.8; IR (neat) 2916, 1083 cm$^{-1}$.

<u>*General Procedure A – Formation of pyrimidinones*</u>

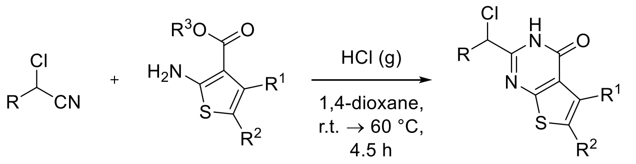


Nitrile (1.2 eq.) was added to a solution of aminothiophene (1 eq.) in freshly distilled 1,4-dioxane (0.5 M). Dry HCl gas was bubbled through the solution which was stirred at room temperature for 1.5 h then 60 °C for 3 h. The reaction mixture was poured into ice-water and basified with 30% aqueous ammonia. The resulting precipitate was collected and dried via vacuum filtration, washing with water, to give the crude material which was purified as indicated.

**2-(1-Chloroethyl)-5,6-dimethylthieno[2,3-*d*]pyrimidin-4(3*H*)-one (3)**

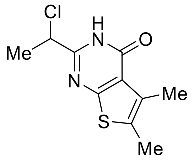

**3**

Synthesised according to *General Procedure A* using 2-chloropropionitrile (645 mg, 7.22 mmol) and ethyl 2-amino-4,5-dimethylthiophene-3-carboxylate (1.20 g, 6.02 mmol) in freshly distilled 1,4-dioxane (12 mL) to give pyrimidinone **3** as a white solid which required no further purification (1.19 g, 81%). Mp: 253 – 262 ºC; $^{1}$H NMR (400 MHz, DMSO-$d_6$) δ 1.79 (d, *J* = 6.8 Hz, 3H), 2.35 (s, 3H), 2.38 (s, 3H), 5.06 (q, *J* = 6.8 Hz, 3H), 12.56 (br s, 1H); $^{13}$C NMR (101 MHz, DMSO-$d_6$) δ 12.7, 12.7, 21.3, 54.3, 122.5, 128.8, 130.4, 154.5, 158.4, 161.1; IR (neat): 3172, 3093, 2931, 2789, 1664 cm$^{-1}$; HRMS (ESI – MS) *m/z* calcd for $C_{10}H_{11}ClN_2OS$ [M+Na]$^+$ 265.01728, found 265.01686.

**2-(1-Chloroethyl)-5-methylthieno[2,3-*d*]pyrimidin-4(3*H*)-one (17)**

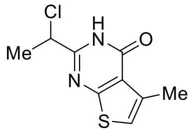

**17**

Synthesised according to *General Procedure A* using 2-chloropropionitrile (152 mg, 1.70 mmol) and ethyl 2-amino-4-methylthiophene-3-carboxylate (261 mg, 1.41 mmol) in freshly distilled 1,4-dioxane (3 mL) to give pyrimidinone **17** as a white solid which required no further purification (304 mg, 94%). Mp: 209 – 210°C; $^{1}$H NMR (400 MHz, DMSO-$d_6$) δ 1.80 (d, *J* = 6.8 Hz, 3H), 2.46 (d, *J* = 1.2 Hz, 3H), 5.07 (q, *J* = 6.8 Hz, 3H), 7.18 (q, *J* = 1.2 Hz, 1H) , 12.63 (br s, 1H); $^{13}$C NMR (101 MHz, DMSO-$d_6$) δ 16.0, 21.3, 54.2, 119.1, 121.9, 133.7, 155.5, 158.8, 164.2; IR (neat): 3092, 2803, 1666 cm$^{-1}$; HRMS (ESI – MS) *m/z* calcd for $C_9H_9ClN_2OS$ [M+Na]$^+$ 251.00163, found 251.00108.

**2-(1-Chloroethyl)-6-methylthieno[2,3-*d*]pyrimidin-4(3*H*)-one (18)**

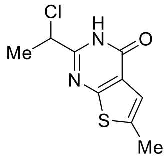

**18**

Synthesised according to *General Procedure A* using 2-chloropropionitrile (2.82 g, 31.5 mmol) and methyl 2-amino-5-methylthiophene-3-carboxylate (4.49 g, 26.2 mmol) in freshly distilled 1,4-dioxane (52 mL). Purification via trituration with ethyl acetate to afford the pyrimidinone **18** as a pink solid (3.00 g, 50%). Mp. 235 – 239 ºC; $^{1}$H NMR (400 MHz, DMSO-$d_6$) δ 1.80 (d, *J* = 6.8 Hz, 3H), 2.51 (d, *J* = 1.2 Hz, 3H), 5.08 (q, *J* = 6.8 Hz, 3H), 7.09 – 7.12 (m, 1H), 12.69 (br s, 1H); $^{13}$C NMR (101 MHz, DMSO-$d_6$) δ 15.5, 21.3, 54.4, 119.1, 124.1, 138.2, 154.9, 157.4, 162.5; IR (neat): 2924, 2793, 1676 cm$^{-1}$; HRMS (ESI – MS) *m/z* calcd for $C_9H_9ClN_2OS$ [M+Na]$^+$ 251.00163, found 250.99925.

<u>*General Procedure B – S-Alkylation Reaction*</u>

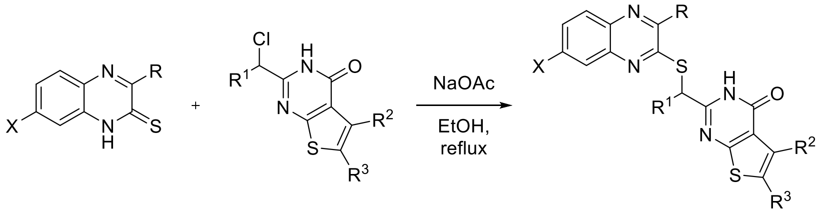

Sodium acetate (1.5 eq.) was added to a suspension of thiol (1 eq.) and chloride (1 eq.) in dry ethanol (0.25 M). The reaction mixture was heated at reflux for the indicated time before the precipitate was collected via vacuum filtration and washed with cold ethanol to afford the product.

**5,6-Dimethyl-2-(1-((3-methylquinoxalin-2-yl)thio)ethyl)thieno[2,3-*d*]pyrimidin-4(3*H*)-one (K12)**

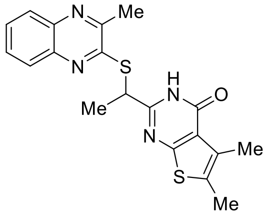

**K12**

Synthesised according to *General Procedure B* using sodium acetate (56 mg, 0.681 mmol), 3-methylquinoxaline-2(1*H*)-thione **4** (80 mg, 0.454 mmol) and 2-(1-chloroethyl)-5,6-dimethylthieno[2,3-*d*]pyrimidin-4(3*H*)-one **3** (110 mg, 0.454 mmol) in dry ethanol (1.8 mL), heating for 5 h, to afford the product (**K12**) as a beige solid (143 mg, 83%). Mp. 227 – 228 °C; $^1$H NMR (400 MHz, DMSO-$d_6$) δ 1.76 (d, *J* = 7.0 Hz, 3H), 2.32 (s, 3H), 2.37 (s, 3H), 2.60 (s, 3H), 5.19 (q, *J* = 7.0 Hz, 1H), 7.65 – 7.80 (m, 2H), 7.86 – 7.99 (m, 2H), 12.51 (br s, 1H); $^{13}$C NMR (101 MHz, DMSO-$d_6$) δ 12.2, 12.3, 18.9, 21.4, 41.4, 121.7, 126.7, 127.9, 128.2, 128.4, 128.9, 129.2, 138.7, 140.1, 151.1, 154.2, 156.7, 158.1, 161.5; IR (neat): 3055, 2919, 1685 cm$^{-1}$; HRMS (ESI – MS) *m/z* calcd for $C_{19}H_{18}N_4OS_2$ [M+H]$^+$ 383.09948, found 383.09878.

**2-(1-((7-Methoxy-3-methylquinoxalin-2-yl)thio)ethyl)-5,6-dimethylthieno[2,3-*d*]pyrimidin-4(3*H*)-one (19 – K12-1)**

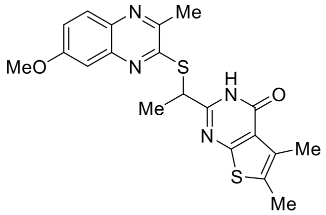

**19**

Synthesised according to *General Procedure B* using sodium acetate (20 mg, 0.24 mmol), 7-methoxy-3-methylquinoxaline-2(1H)-thione **16** (33 mg, 0.160 mmol) and 2-(1-chloroethyl)-5,6-dimethylthieno[2,3-*d*]pyrimidin-4(3*H*)-one **3** (39 mg, 0.160 mmol) in dry ethanol (1.15 mL), heating for 4 h, to afford the product **19** as a beige solid (35 mg, 53%). Mp: 235 – 244 °C; $^1$H NMR (400 MHz, DMSO-$d_6$) δ 1.76 (d, *J* = 7.1 Hz, 3H), 2.31 (s, 3H), 2.36 (s, 3H), 2.53 (s, 3H), 3.91 (s, 3H), 5.20 (q, *J* = 7.1 Hz, 1H), 7.25 – 7.27 (m, 1H), 7.28 – 7.35 (m, 1H), 7.83 (d, *J* = 9.1 Hz, 1H), 12.59 (s, 1H); $^{13}$C NMR (101 MHz, DMSO-$d_6$) δ 12.6, 12.8, 19.3, 21.3, 41.4, 55.7, 105.8, 120.5, 121.9, 128.6, 129.2, 129.2, 134.6, 141.8, 148.3, 154.3, 157.1, 158.5, 159.9, 161.8; IR (neat): 3119, 2918, 1680 cm$^{-1}$; HRMS (ESI – MS) *m/z* calcd for $C_{20}H_{20}N_4O_2S_2$ [M+Na]$^+$ 435.09199, found 435.09124.

**5,6-Dimethyl-2-(1-(quinoxalin-2-ylthio)ethyl)thieno[2,3-*d*]pyrimidin-4(3*H*)-one (20 – K12-2)**

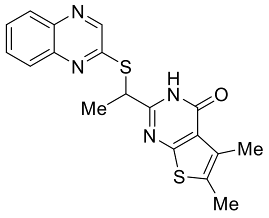

**20**

Synthesised according to *General Procedure B* using sodium acetate (38 mg, 0.462 mmol), quinoxaline-2(1*H*)-thione **12** (50 mg, 0.308 mmol) and 2-(1-chloroethyl)-5,6-dimethylthieno[2,3-*d*]pyrimidin-4(3*H*)-one **3** (75 mg, 0.308 mmol) in dry ethanol (1.2 mL), heating for 2 h, to afford the crude material which was recrystallised from ethanol to give product **20** as a beige solid (15 mg, 13%). Mp. 212 – 213 °C; $^1$H NMR (400 MHz, DMSO-$d_6$) δ 1.76 (d, *J* = 6.9 Hz, 3H), 2.32 (s, 3H), 2.36 (s, 3H), 5.17 (q, *J* = 6.9 Hz, 1H), 7.69 – 7.78 (m, 1H), 7.78 – 7.87 (m, 1H), 7.88 – 7.95 (m, 1H), 7.99 –

8.06 (m, 1H), 8.82 (s, 1H), 12.59 (br s, 1H); $^{13}$C NMR (101 MHz, DMSO-$d_6$) δ 12.6, 12.8, 19.4, 41.4, 121.9, 127.4, 128.7, 128.8, 129.0, 129.3, 130.8, 19.4, 141.7, 144.7, 154.8, 156.8, 158.5, 161.7; IR (neat): 3395, 3134, 3054, 2931, 1685 cm$^{-1}$; HRMS (ESI – MS) *m/z* calcd for $C_{18}H_{16}N_4OS_2$ [M+H]$^+$ 369.08383, found 369.08301.

**2-(1-((3-Ethylquinoxalin-2-yl)thio)ethyl)-5,6-dimethylthieno[2,3-*d*]pyrimidin-4(3*H*)-one (21 – K12-3)**

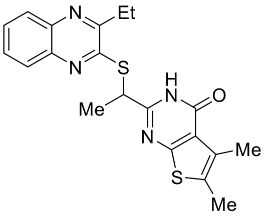


**21**

Synthesised according to *General Procedure B* using sodium acetate (16 mg, 0.197 mmol), 3-ethylquinoxaline-2(1*H*)-thione **13** (25 mg, 0.131 mmol), 2-(1-chloroethyl)-5,6-dimethylthieno[2,3-*d*]pyrimidin-4(3*H*)-one **3** (32 mg, 0.131 mmol) in dry ethanol (1.02 mL), heating for 6 h, to afford the product **21** as a white fluffy solid (27 mg, 52%). Mp. 220 – 225 °C; $^{1}$H NMR (400 MHz, DMSO-$d_6$) δ 1.33 (t, *J* = 7.4 Hz, 3H), 1.75 (d, *J* = 7.0 Hz, 3H), 2.33 (s, 3H), 2.37 (s, 3H), 2.91 (q, *J* = 7.4 Hz, 2H), 5.19 (q, *J* = 7.0 Hz, 1H), 7.67 – 7.79 (m, 2H), 7.87 – 7.91 (m, 1H), 7.95 – 8.00 (m, 1H), 12.59 (br s, 1H); $^{13}$C NMR (101 MHz, DMSO-$d_6$) δ 10.6, 12.2, 12.3, 18.9, 27.1, 41.4, 121.7, 126.7, 128.1, 128.2, 128.4, 128.9, 129.2, 138.8, 140.0, 153.9, 155.0, 156.8, 158.1, 161.6; IR (neat): 3107, 3035, 2975, 2928, 1684 cm$^{-1}$; HRMS (ESI – MS) *m/z* calcd for $C_{20}H_{20}N_4OS_2$ [M+Na]$^+$ 419.09707, found 419.09623.

**5-Methyl-2-(1-((3-methylquinoxalin-2-yl)thio)ethyl)thieno[2,3-*d*]pyrimidin-4(3*H*)-one (22 – K12-4)**

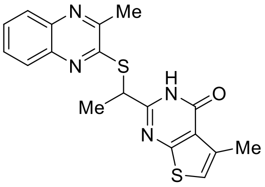


**22**

Synthesised according to *General Procedure B* using sodium acetate (15 mg, 0.187 mmol), 3-methylquinoxaline-2(1*H*)-thione **4** (22 mg, 0.125 mmol) and 2-(1-chloroethyl)-5-methylthieno[2,3-*d*]pyrimidin-4(3*H*)-one **17** (28 mg, 0.125 mmol) in dry ethanol (0.5 mL), heating for 4 h, to afford the product **22** as a pink solid (36 mg, 78%). Mp: 243 – 246 °C; $^{1}$H NMR (600 MHz, $CDCl_3$) δ 1.92 (d, *J* = 7.4 Hz, 3H), 2.52 (d, *J* = 1.3 Hz, 3H), 2.68 (s, 3H), 5.08 (q, *J* = 7.4 Hz, 1H), 6.78 (d, *J* = 1.3 Hz, 1H), 7.65 – 7.71 (m, 1H), 7.71 – 7.76 (m, 1H), 7.97 – 8.02 (m, 1H), 8.08 – 8.13 (m, 1H), 11.90 (br s, 1H); $^{13}$C NMR (151 MHz, $CDCl_3$) δ 16.5, 17.1, 22.5, 40.9, 118.0, 122.4, 126.9, 128.8, 129.2, 130.4, 135.0, 140.2, 140.3, 151.8, 155.2, 156.9, 159.2, 165.6; IR (neat): 3126, 3052, 1683 cm$^{-1}$; HRMS (ESI – MS) *m/z* calcd for $C_{18}H_{16}N_4OS_2$ [M+H]$^+$ 369.08383, found 369.08323.

**6-Methyl-2-(1-((3-methylquinoxalin-2-yl)thio)ethyl)thieno[2,3-*d*]pyrimidin-4(3*H*)-one (23 – K12-5)**

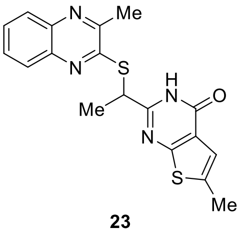


**23**

Synthesised according to *General Procedure B* using sodium acetate (49 mg, 0.596 mmol), 3-methylquinoxaline-2(1*H*)-thione **4** (70 mg, 0.397 mmol), 2-(1-chloroethyl)-6-methylthieno[2,3-*d*]pyrimidin-4(3*H*)-one **18** (91 mg, 0.397 mmol) in dry ethanol (1.6 mL), heating for 7 h. The crude material was purified on deactivated silica eluting with ethyl acetate, to afford the product **23** as a pale pink solid (95 mg, 65%). Mp. 205 °C (dec); $^{1}$H NMR (400 MHz, DMSO-$d_6$) δ 1.77 (d, *J* = 7.0 Hz, 3H), 2.47 (d, *J* = 1.2 Hz, 3H), 2.60 (s, 3H), 5.21 (q, *J* = 7.0 Hz, 1H), 7.06 (d, *J* = 1.2 Hz, 1H), 7.66 – 7.78 (m, 2H), 7.85 – 7.90 (m, 1H), 7.92 – 7.97 (m, 1H), 12.72 (br s, 1H); $^{13}$C NMR* (101 MHz, DMSO-$d_6$) δ 15.5, 19.3, 21.7, 41.7, 119.0, 123.3, 126.9, 128.1, 128.5, 129.6, 136.9, 138.9, 140.2, 151.3, 154.4, 157.4, 157.5, 163.1; IR (neat): 3104, 3054, 2930, 1696 cm$^{-1}$; HRMS (ESI – MS) *m/z* calcd for $C_{18}H_{16}N_4OS_2$ [M+Na]$^+$ 391.06577, found 391.06521.

*Due to solubility issues, $^{13}$C NMR experiment was completed at 65 °C.

##### 5,6-Dimethyl-2-(((3-methylquinoxalin-2-yl)thio)methyl)thieno[2,3-*d*]pyrimidin-4(3*H*)-one (24 – K12-6)

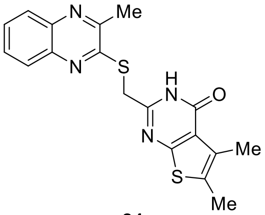

**24**

Synthesised according to *General Procedure B* using sodium acetate (33 mg, 0.408 mmol), 3-methylquinoxaline-2(1*H*)-thione **4** (48 mg, 0.272 mmol), 2-(chloromethyl)-5,6-dimethylthieno[2,3-*d*]pyrimidin-4(3*H*)-one **14** (62 mg, 0.272 mmol) in dry ethanol (1.1 mL), heating for 3.5 h, to afford the product **24** as a pink solid (75 mg, 75%). Mp. 253-255 °C; $^{1}$H NMR (400 MHz, DMSO-d$_6$) δ 2.31 (s, 3H), 2.37 (s, 3H), 2.64 (s, 2H), 4.56 (s, 2H), 7.66 – 7.78 (m, 2H), 7.86 – 7.98 (m, 2H), 12.54 (s, 1H); $^{13}$C NMR (101 MHz, DMSO-d$_6$) δ 13.1, 13.1, 22.5, 33.5, 123.1, 126.9, 128.8, 129.3, 129.7, 130.4, 130.7, 140.3, 140.4, 151.9, 152.8, 154.9, 158.8, 162.3; IR (neat) 2849, 1649 cm$^{-1}$; HRMS (ESI-MS): *m/z* calcd for $C_{18}H_{16}N_4OS_2$ [M + Na]$^+$ 391.0658, found 391.0623.

**<u>Synthesis of the K12 bead</u>**

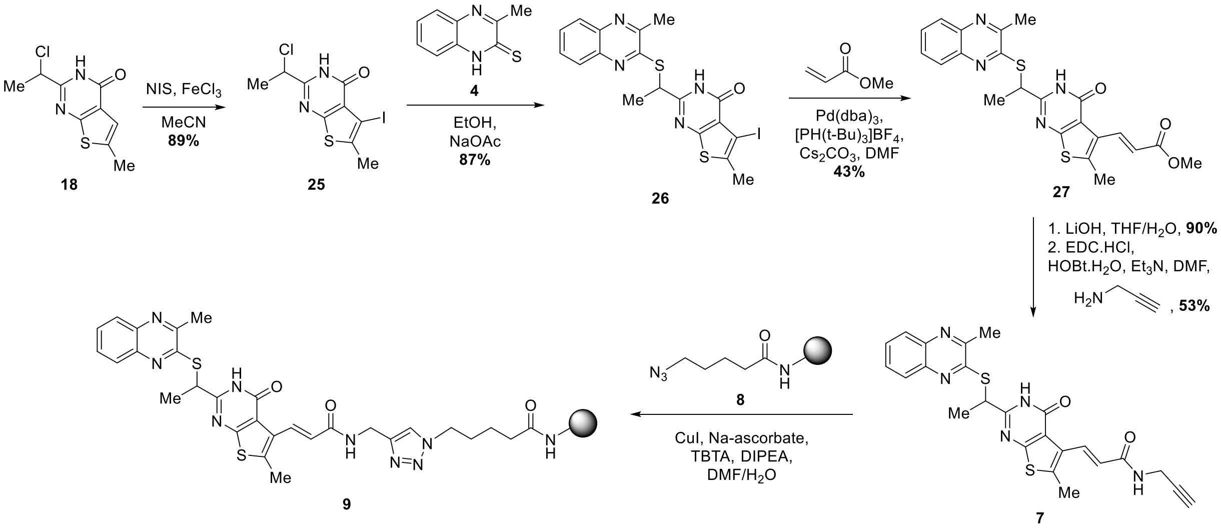


##### 2-(1-Chloroethyl)-5-iodo-6-methylthieno[2,3-*d*]pyrimidin-4(3*H*)-one (25)

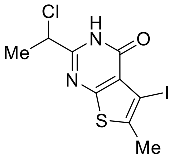

**25**

*N*-Iodosuccinimide (2.52 g, 11.2 mmol) and $FeCl_3$ (182 mg, 1.12 mmol) were added to a suspension of thienopyrimidinone **18** (1.28 g, 5.60 mmol) in acetonitrile (56 mL). The reaction mixture was stirred at reflux for 18 h. The solution was cooled to room temperature, diluted with saturated aqueous sodium bicarbonate solution, and extracted with DCM (x 6). The combined organic extracts were washed aqueous sodium bicarbonate solution, a 1M solution of sodium thiosulfate, brine and dried ($Na_2SO_4$). The solvent was removed under reduced pressure to afford the product **25** as a brown solid (1.77 g, 89%). Mp. 220 – 223 °C; $^{1}$H NMR (400 MHz, DMSO-d$_6$) δ 1.79 (d, *J* = 6.8 Hz, 3H), 2.43 (s, 3H), 5.06 (q, *J* = 6.8 Hz, 3H), 12.79 (br s, 1H); $^{13}$C NMR (101 MHz, DMSO-d$_6$) δ 18.2, 21.2, 54.1, 75.9, 122.1, 135.8, 155.3, 156.8, 161.9; IR (neat): 2919, 2856, 1657 cm$^{-1}$; HRMS (ESI – MS) *m/z* calcd for $C_9H_8ClIN_2OS$ [M+Na]$^+$ 376.89828, found 376.89526.

**5-Iodo-6-methyl-2-(1-((3-methylquinoxalin-2-yl)thio)ethyl)thieno[2,3-*d*]pyrimidin-4(3*H*)-one (26)**

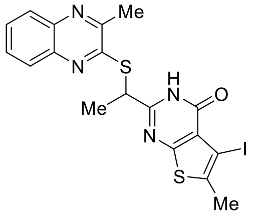

26

Synthesised according to *General Procedure B* using sodium acetate (164 mg, 2.00 mmol), 3-methylquinoxaline-2(1*H*)-thione **4** (254 mg, 1.33 mmol) and 2-(1-chloroethyl)-5-iodo-6-methylthieno[2,3-*d*]pyrimidin-4(3*H*)-one **25** (473 mg, 1.33 mmol) in dry ethanol (5.3 mL), heating for 17 h, to afford the product **26** as a brick red solid (570 mg, 87%). Mp: 212-216 °C (dec); $^1$H NMR (400 MHz, DMSO-d$_6$) δ 1.76 (d, *J* = 7.0 Hz, 3H), 2.39 (s, 3H), 2.60 (s, 3H), 5.19 (q, *J* = 7.0 Hz, 1H), 7.66 – 7.79 (m, 2H), 7.85 – 7.90 (m, 1H), 7.92 – 7.97 (m, 1H), 12.83 (br s, 1H); $^{13}$C NMR (101 MHz, DMSO-d$_6$) δ 18.1, 19.1, 21.7, 41.6, 75.8, 121.4, 126.9, 128.1, 128.5, 129.6, 134.6, 138.9, 140.2, 151.3, 154.4, 156.8, 158.0, 162.5; IR (neat): 3360, 2913, 1694 cm$^{-1}$; HRMS (ESI – MS) *m/z* calcd for $C_{18}H_{15}IN_4OS_2$ [M+Na]$^+$ 516.96242, found 516.96168.

**Methyl (*E*)-3-(6-methyl-2-(1-((3-methylquinoxalin-2-yl)thio)ethyl)-4-oxo-3,4-dihydrothieno[2,3-*d*]pyrimidin-5-yl)acrylate (27)**

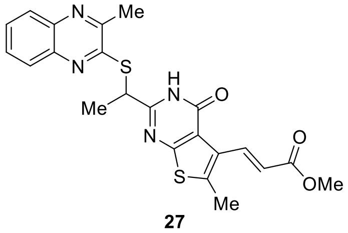

27

Iodide **26** (553 mg, 1.12 mmol), methyl acrylate (405 μL, 4.47 mmol) cesium carbonate (1.18 g, 3.38 mmol), and DMF (7.5 mL) were added to a flame-dried Schlenk tube and degassed (Freeze-Pump-Thaw x 2). $Pd_2(dba)_3$ (102 mg, 0.112 mmol) and tri-*tert*-butylphosphonium tetrafluoroborate (65 mg, 0.224 mmol) were added and the reaction mixture was degassed (Freeze-Pump-Thaw x 2) before being heated at 80 ºC for 22 h. The reaction mixture was filtered through a pad of Celite, eluting with dichloromethane. The organic layer was washed with water (x 5), brine and dried ($Na_2SO_4$). The solvent was removed under reduced pressure and the crude material was purified by flash chromatography, eluting with 10 – 20 % diethyl ether/dichloromethane, to afford the product **27** as a white solid (220 mg, 43%). Mp. 246 – 247 °C; $^1$H NMR (400 MHz, $CDCl_3$) δ 1.91 (d, *J* = 7.4 Hz, 3H), 2.60 (s, 3H), 2.69 (s, 3H), 3.76 (s, 3H), 5.07 (q, *J* = 7.4 Hz, 1H), 6.43 (d, *J* = 16.3 Hz, 1H), 7.66 – 7.78 (m, 2H), 8.00 (dd, *J* = 8.1, 1.4 Hz, 1H), 8.11 (dd, *J* = 8.1, 1.4 Hz, 1H), 8.23 (d, *J* = 16.3 Hz), 11.98 (s, 1H); $^{13}$C NMR (101 MHz, DMSO-d$_6$) δ 15.3, 19.1, 21.7, 41.5, 51.5, 120.7, 121.1, 126.9, 127.3, 128.2, 128.6, 129.6, 136.8, 137.9, 138.9, 140.2, 151.3, 154.4, 158.0, 158.5, 162.1, 168.8; IR (neat): 2950, 1691 cm$^{-1}$; HRMS (ESI – MS) *m/z* calcd for $C_{22}H_{20}N_4O_3S_2$ [M+Na]$^+$ 475.08690, found 475.08596.

**(*E*)-3-(6-Methyl-2-(1-((3-methylquinoxalin-2-yl)thio)ethyl)-4-oxo-3,4-dihydrothieno[2,3-*d*]pyrimidin-5-yl)acrylic acid (28)**

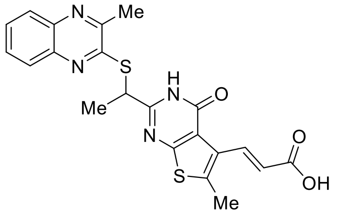

28

Lithium hydroxide (58 mg, 2.46 mmol) was added to a suspension of ester **27** (185 mg, 0.409 mmol) in tetrahydrofuran/$H_2O$ (1:1, 4.6 mL) and the reaction mixture was stirred at 60 ºC for 24 h. The volatiles were removed under reduced pressure and the reaction mixture was adjusted to pH 1 using hydrochloric acid (2 M). The resulting precipitate was collected via vacuum filtration, washed with water to afford the product **28** as a beige solid (161 mg, 90%). Mp. 168 °C; $^1$H NMR (400 MHz, DMSO-d$_6$) δ 1.77 (d, *J* = 7.0 Hz, 3H), 2.57 (s, 3H), 2.60 (s, 3H), 5.20 (q, *J* = 7.0 Hz, 1H), 6.36 (d, *J* = 16.3 Hz, 1 H), 7.66 – 7.79 (m, 2H), 7.88 (dd, *J* = 8.2, 1.3 Hz, 1H), 7.94 (dd, *J* = 8.2, 1.3 Hz, 1H), 8.19 (d, *J* = 16.3 Hz, 1H), 12.39 (br s, 1H), 12.90 (s, 1H); $^{13}$C NMR (101 MHz, DMSO-d$_6$) δ 15.4, 19.2, 21.7, 41.5, 120.7, 122.5, 127.0, 127.6, 128.2, 128.6, 129.7, 136.4, 137.3, 138.9, 140.3, 151.4, 154.4, 158.1, 158.5, 162.0, 167.7; IR (neat): 3388, 2933, 1655, 1579 $cm^{-1}$; HRMS (ESI – MS) *m/z* calcd for $C_{21}H_{18}N_4O_3S_2$ $[M+Na]^+$ 461.07125, found 461.07014.

**(*E*)-3-(6-methyl-2-(1-((3-methylquinoxalin-2-yl)thio)ethyl)-4-oxo-3,4-dihydrothieno[2,3-*d*]pyrimidin-5-yl)-*N*-(prop-2-yn-1-yl)acrylamide (7)**

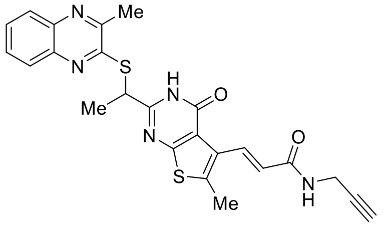

**7**

Triethylamine (50 µL, 0.359 mmol) was added to a solution of acid **28** (63 mg, 0.144 mmol) in DMF (1.4 mL). The solution was cooled to 0 ºC before 1-ethyl-3-(3-dimethylaminopropyl)carbodiimide hydrochloride (33 mg, 0.172 mmol), 1-hydroxybenzotriazole hydrate (26 mg, 0.172 mmol) and propargylamine (18 µL, 0.287 mmol) were added sequentially and the solution was allowed to warm to room temperature. After stirring for 24 h the reaction mixture was diluted with water and extracted with dichloromethane (x 3). Combined organic extracts were washed with saturated aqueous sodium bicarbonate solution, water (x 2), brine and dried ($Na_2SO_4$). The solvent was removed under reduced pressure and the crude material was purified by flash chromatography on deactivated silica gel eluting with 1 % methanol/dichloromethane, to afford the product **7** as an off-white solid (36 mg, 53%). Mp. 195 – 200 °C (dec); $^1H$ NMR (400 MHz, $CDCl_3$) δ 1.91 (d, *J* = 7.4 Hz, 3H), 2.21 (t, *J* = 2.6 Hz, 1H), 2.59 (s, 3H), 2.69 (s, 3H), 4.12 – 4.15 (m, 2H), 5.07 (q, *J* = 7.4 Hz, 1H), 5.89 – 5.95 (m, 1H), 6.89 (d, *J* = 15.9 Hz, 1H), 7.67 – 7.78 (m, 2H), 7.86 (d, *J* = 15.9 Hz, 1H), 8.00 (dd, *J* = 8.2, 1.3 Hz, 1H), 8.10 (dd, *J* = 8.2, 1.3 Hz, 1H), 11.97 (br s, 1H); $^{13}C$ NMR (101 MHz, DMSO-$d_6$) δ 15.5, 19.2, 21.7, 28.1, 41.5, 73.2, 81.1, 120.9, 125.0, 127.0, 128.1, 128.2, 128.6, 129.7, 132.2, 135.3, 138.9, 140.3, 151.3, 154.4, 158.1, 158.3, 161.9, 164.9; IR (neat): 3247, 3058, 1687 $cm^{-1}$; HRMS (ESI – MS) *m/z* calcd for $C_{21}H_{21}N_5O_2S_2$ $[M+Na]^+$ 498.10289, found 498.10178.

**Synthesis of negative control bead**

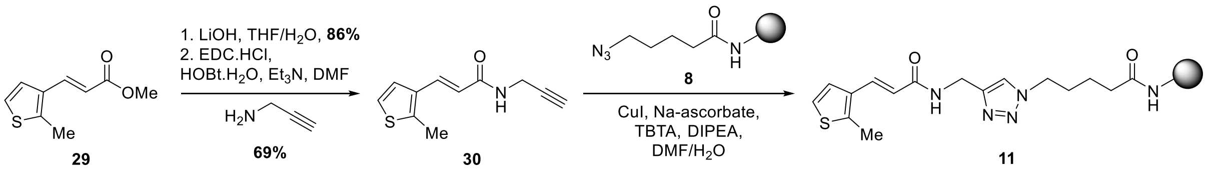


**Methyl (*E*)-3-(2-methylthiophen-3-yl)acrylate (29)[16]**

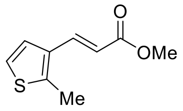

**29**

*n*-Butyllithium (2.5 M in hexanes, 813 µL, 2.03 mmol) was added dropwise to a solution of 3-bromo-2-methylthiophene (300 mg, 1.69 mmol) in dry diethyl ether (2.1 mL) at -78 ºC and the mixture was stirred at this temperature for 45 min. DMF (262 µL, 3.39 mmol) in dry diethyl ether (2.1 mL) was cooled to -78 ºC and added to the reaction mixture dropwise. The solution was allowed to warm to room temperature over 3 h and stirred at this temperature for a further 15 min before quenching with saturated aqueous ammonium chloride solution and extracting with diethyl ether (x 4). Combined organic extracts were washed with water (x 2), brine and dried ($Na_2SO_4$). The solvent was removed carefully under reduced pressure to afford the crude residue of 2-methylthiophene-3-carbaldehyde as a yellow liquid which was used without further purification. The crude residue was dissolved in dichloromethane (8.5 mL) and methyl (triphenylphosphoranylidene)acetate (848 mg, 2.54 mmol) was added and stirred at room temperature for 16 h. Volatiles were removed under reduced pressure and the crude material was purified by flash chromatography eluting with 10 % diethyl ether/petroleum spirits, to afford the product **29** as a colourless liquid (102 mg, 33% over two steps), with analytical data matching that reported in the literature.[16] $^1H$ NMR (400 MHz, $CDCl_3$) δ 2.55 (s, 3H), 3.79 (s,

3H), 6.19 (d, *J* = 15.8 Hz, 1H), 7.06 (d, *J* = 5.4 Hz, 1H), 7.17 (d, *J* = 5.4 Hz, 1H), 7.70 (d, *J* = 15.8 Hz, 1H).

**(*E*)-3-(2-Methylthiophen-3-yl)acrylic acid (30)**

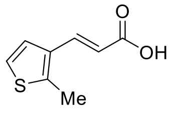

**30**

Lithium hydroxide (1 M, aq., 2.19 mL) was added to a solution of ester **29** (100 mg, 0.549 mmol) in tetrahydrofuran (12 mL) and stirred at 60 ºC for 4 h. Volatiles were removed under reduced pressure and the reaction mixture was washed with dichloromethane (x 3). The solution was adjusted to pH 2 using hydrochloric acid (2 M) and extracted with dichloromethane (x 5). Combined organic extracts were washed with brine and dried ($Na_2SO_4$). The solvent was removed under reduced pressure to afford the product **30** as a white solid (79 mg, 86%). Mp. 143 – 145 °C; $^1$H NMR (400 MHz, $CDCl_3$) δ 2.57 (s, 3H), 6.20 (d, *J* = 15.8 Hz, 1H), 7.09 (d, *J* = 5.4 Hz, 1H), 7.20 (d, *J* = 5.4 Hz, 1H), 7.79 (d, *J* = 15.8 Hz, 1H); $^{13}$C NMR (101 MHz, DMSO-$d_6$) δ 12.8, 118.0, 123.5, 125.9, 132.9, 135.6, 141.8, 168.1; IR (neat): 2818, 1670, 1608 $cm^{-1}$; HRMS (ESI – MS) *m/z* calcd for $C_8H_8O_2S$ $[M+Na]^+$ 191.01372, found 191.01343.

**(*E*)-3-(2-methylthiophen-3-yl)-*N*-(prop-2-yn-1-yl)acrylamide (10)**

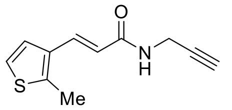

**10**

Triethylamine (81 µL, 0.580 mmol) was added to a solution of acid **30** (39 mg, 0.232 mmol) in DMF (2.3 mL). The solution was cooled to 0 ºC before 1-ethyl-3-(3-dimethylaminopropyl)carbodiimide hydrochloride (53 mg, 0.278 mmol), 1-hydroxybenzotriazole hydrate (42 mg, 0.278 mmol) and propargylamine (30 µL, 0.464 mmol) were added sequentially and the solution was allowed to warm to room temperature. After stirring for 22 h the reaction mixture was diluted with water and extracted with dichloromethane (x 3). The combined organic extracts were washed with saturated aqueous sodium bicarbonate solution, brine and dried ($Na_2SO_4$). The solvent was removed under reduced pressure and the crude material was purified on silica eluting with 40 % ethyl acetate/petroleum spirits, to afford the product **10** as a white solid (33 mg, 69%). Mp. 157 – 159 °C; $^1$H NMR (400 MHz, $CDCl_3$) δ 2.26 (t, *J* = 2.6 Hz, 1H), 2.54 (s, 3H), 4.17 – 4.20 (m, 2H), 5.69 (br s, 1H), 6.14 (d, *J* = 15.4 Hz, 1H), 7.05 (d, *J* = 5.4 Hz, 1H), 7.13 (d, *J* = 5.4 Hz, 1H), 7.68 (d, *J* = 15.4 Hz, 1H); $^{13}$C NMR (101 MHz, $CDCl_3$) δ 13.4, 29.6, 71.8, 79.7, 118.6, 122.7, 125.2, 133.1, 133.9, 142.0, 166.2; IR (neat): 3222, 3060, 2922, 1646 $cm^{-1}$; HRMS (ESI – MS) *m/z* calcd for $C_{11}H_{11}NOS$ $[M+Na]^+$ 228.04536, found 228.04497.

**Attachment of K12 and negative control substrate to affinity chromatography bead[2]**

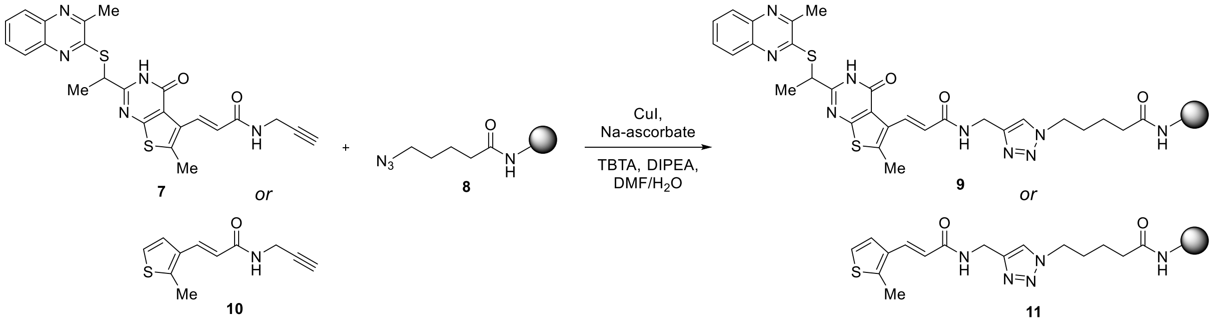


50 mM stock solutions of copper(I) iodide, *N,N*-diisopropylethylamine, TBTA and alkyne **7** or **10** in DMF were made. A 50 mM stock solution of sodium ascorbate in $H_2O$ was made. Azide-linker Carboxylink beads (prepared according to the protocol of Finn et al.[2]) were washed with DMF (10 mL) and transferred to a 20 mL vial. The vial was placed under an atmosphere of argon and the solutions of *N,N*-diisopropylethylamine (8 eq.), TBTA (4 eq.) and alkyne (3 eq.) were added. The solution was bubbled gently with argon for 15 min before the solutions of copper(I) iodide (4 eq.) and sodium ascorbate (8 eq.) were added. The vial was capped and

shaken gently (150 rpm) for 20 h. The suspension was transferred back to the Carboxylink column housing and the solution was drained. The remaining beads were washed successively with DMF (20 mL), water (20 mL), methanol (20 mL), 0.1 M aqueous NaEDTA solution (20 mL), water (20 mL) and DMF (20 mL), 1 M aqueous sodium chloride solution (6 mL) and stored under 1 M aqueous sodium chloride solution (2 mL, 0.05 % sodium azide).

**References**


1. H. A. El-Sayed, S. S. A., M. A. H., B. M. M. and R. T. and Abdel-Kader, *Nucleosides Nucleotides Nucl. Acids*, 2016, **35**, 16-31.
2. S. Punna, E. Kaltgrad and M. G. Finn, *Bioconjugate Chemistry*, 2005, **16**, 1536-1541.
3. Q. Huang, Katt, W.P., McDermott, L.A., Cerione, R.A., unpublished work.
4. R. A. Friesner, J. L. Banks, R. B. Murphy, T. A. Halgren, J. J. Klicic, D. T. Mainz, M. P. Repasky, E. H. Knoll, M. Shelley, J. K. Perry, D. E. Shaw, P. Francis and P. S. Shenkin, *J. Med. Chem.*, 2004, **47**, 1739-1749.
5. T. A. Halgren, R. B. Murphy, R. A. Friesner, H. S. Beard, L. L. Frye, W. T. Pollard and J. L. Banks, *J. Med. Chem.*, 2004, **47**, 1750-1759.
6. R. A. Friesner, R. B. Murphy, M. P. Repasky, L. L. Frye, J. R. Greenwood, T. A. Halgren, P. C. Sanschagrin and D. T. Mainz, *J. Med. Chem.*, 2006, **49**, 6177-6196.
7. E. Harder, W. Damm, J. Maple, C. Wu, M. Reboul, J. Y. Xiang, L. Wang, D. Lupyan, M. K. Dahlgren, J. L. Knight, J. W. Kaus, D. S. Cerutti, G. Krilov, W. L. Jorgensen, R. Abel and R. A. Friesner, *J. Chem. Theory Comput.*, 2016, **12**, 281-296.
8. J. C. Shelley, A. Cholleti, L. L. Frye, J. R. Greenwood, M. R. Timlin and M. Uchimaya, *Journal of Computer-Aided Molecular Design*, 2007, **21**, 681-691.
9. J. Michel and J. W. Essex, *Journal of Computer-Aided Molecular Design*, 2010, **24**, 639-658.
10. G. R. Fulmer, A. J. M. Miller, N. H. Sherden, H. E. Gottlieb, A. Nudelman, B. M. Stoltz, J. E. Bercaw and K. I. Goldberg, *Organometallics*, 2010, **29**, 2176-2179.
11. A. Makhloufi, B. M., M. M. and D. and Benachour, *Synth. Commun.*, 2011, **41**, 3532-3540.
12. G. A. El-Hiti, *Synthesis*, 2003, **2003**, 2799-2804.
13. T. J. Fyfe, B. Zarzycka, H. D. Lim, B. Kellam, S. N. Mistry, V. Katrich, P. J. Scammells, J. R. Lane and B. Capuano, *J. Med. Chem.*, 2019, **62**, 174-206.
14. A. N. Matthew, J. Zephyr, C. J. Hill, M. Jahangir, A. Newton, C. J. Petropoulos, W. Huang, N. Kurt-Yilmaz, C. A. Schiffer and A. Ali, *J. Med. Chem.*, 2017, **60**, 5699-5716.
15. M. I. Abasolo, C. H. Gaozza and B. M. Fernández, *J. Heterocycl. Chem.*, 1987, **24**, 1771-1775.
16. H. Satonaka, *Magn. Reson. Chem.*, 1986, **24**, 265-267.
